# Transition Metal Binding Drives Folding of a Metalloregulatory Riboswitch by Modulating Conformational Flexibility at Helical Junctions

**DOI:** 10.1101/2025.08.02.668262

**Authors:** Dibyendu Mondal, Sk Habibullah, Lipika Baidya, Mahesh Singh Harariya, Govardhan Reddy

**Author notes:** Contributed equally to this work (Dibyendu Mondal, Sk Habibullah).

## Abstract

Transition metal ions are crucial for bacteria’s survival. Bacteria employ metalloregulatory riboswitches to respond to varying metal ion concentrations. The czcD (NiCo) transcription riboswitch specifically senses Co^2+^, Ni^2+^, and Fe^2+^ ions at micromolar concentrations amid millimolar Mg^2+^. We used computer simulations with multi-resolution RNA models to understand how global conformational changes in the NiCo riboswitch are coupled to the remarkable specific binding of Co^2+^. We show that the riboswitch folds through an intermediate state, where a partially folded four-way junction (4WJ) creates an anionic pocket large enough to accommodate the binding of solvated divalent ions. The binding of Co^2+^ is coupled to the stability of the weak non-canonical G*·*A base pairs at the helical junction that drive the formation of native-like coaxial stacking of four helices. The Co^2+^ binding further twists the 4WJ, which locks the ions in the bound state. Electronic structure calculations show that enhanced orbital interactions between conserved guanines in the 4WJ and Co^2+^ are responsible for the high specificity of the riboswitch in binding to Co^2+^ over Mg^2+^. We provide a framework for understanding and engineering tunable RNA-based biosensors and developing antimicrobials, as metal intoxication is an evolutionary strategy to inhibit bacterial growth.

## 1 Introduction

Bacteria adapt and survive by detecting and responding to changes in the concentrations of various metal ions in their environment.^1^ Metal ions play pivotal roles, such as the stabilisation of folded biomolecular structures (Mg^2+^ and Co^2+^), redox catalysts (Fe^2+^ and Cu^2+^) and non-redox catalysts (Zn^2+^ and Mn^2+^).^2^ The scarcity of metal ions limits bacterial growth. However, excess availability leads to toxicity through mismetallation, where the metals present in excess replace the cognate ligand metal ions in proteins and RNA.^3,4^ Bacteria employ regulatory mechanisms, including metalloregulatory DNA-binding proteins and riboswitches, to maintain the balance of vital metal ions.^5,6^

Riboswitches are structured RNA elements found in the 5^*′*^-untranslated region of bacterial mRNAs that regulate gene expression by binding to their cognate ligands.^7–9^ Currently, more than 55 different classes of riboswitches have been identified^10^ that selectively bind to metal ions,^11–14^ co-factors,^15^ signaling molecules,^16^ tRNA,^17^ and metabolites.^18,19^ Metalloregulatory riboswitches selectively bind to transition metal ions such as Mn^2+^, Fe^2+^, Co^2+^, and Ni^2+^ present in micromolar concentrations in bacterial cells in the presence of monovalent (Na^+^, K^+^) and divalent (Mg^2+^, Ca^2+^) ions present in millimolar concentrations.^12,13,20,21^ The crystal structures of several metal-binding riboswitches^11,12,22^ reveal the bound pose of the metal ion in the folded riboswitch structure. However, the molecular mechanisms underlying the metal ion recognition, specificity, and signal transduction by the riboswitches remain poorly understood. It is essential to address these questions to understand how bacteria maintain metal homeostasis, and these insights will help design metal ion sensors and antibacterial molecules.^23^

The NiCo riboswitch aptamer (NRA) is a single-stranded noncoding RNA that resides upstream of czcD genes that encode for a subfamily of metal export and resistance proteins. The folded state of NRA consists of four helices (labeled P1, P2, P3, and P4) and a four-way junction (4WJ) at the core of the structure (Fig. 1A-C).^12^ An excess concentration of Fe^2+^, Co^2+^, or Ni^2+^ leads to the binding of these ions to the binding pocket in the 4WJ, which results in conformational transitions that promote mRNA transcription by RNA polymerase, and subsequent production of proteins that aid in the efflux of excess ions. X-ray crystallography showed that four Co^2+^ ions bind to the 4WJ through a combination of inner-shell (IS) contacts with N7 and O2^*′*^ atoms in six conserved guanosines and one adenosine.^12^ In-line probing experiments showed that the binding of Co^2+^ ions to the 4WJ is cooperative. ^12^ Single-molecule fluorescence resonant energy transfer (smFRET) studies on NRA demonstrated that in the presence of Mg^2+^ and/or high concentration of Na^+^, NRA folds to an intermediate state from unfolded conformations.^20^ The experiments further show that when cognate transition metal ions are added, the intermediate state folds to the native state. However, the following questions remain unclear: (i) What is the structure of the intermediate state and the key interactions that govern the formation of the native-like helical arrangement? (ii) At what stage of the folding process do transition metal ions bind to the riboswitch, and how does their binding drive the structural transitions to the folded state? (iii) What interactions are critical for the selective binding of multiple Co^2+^ over Mg^2+^, and how is their binding cooperative?

**Figure 1.**
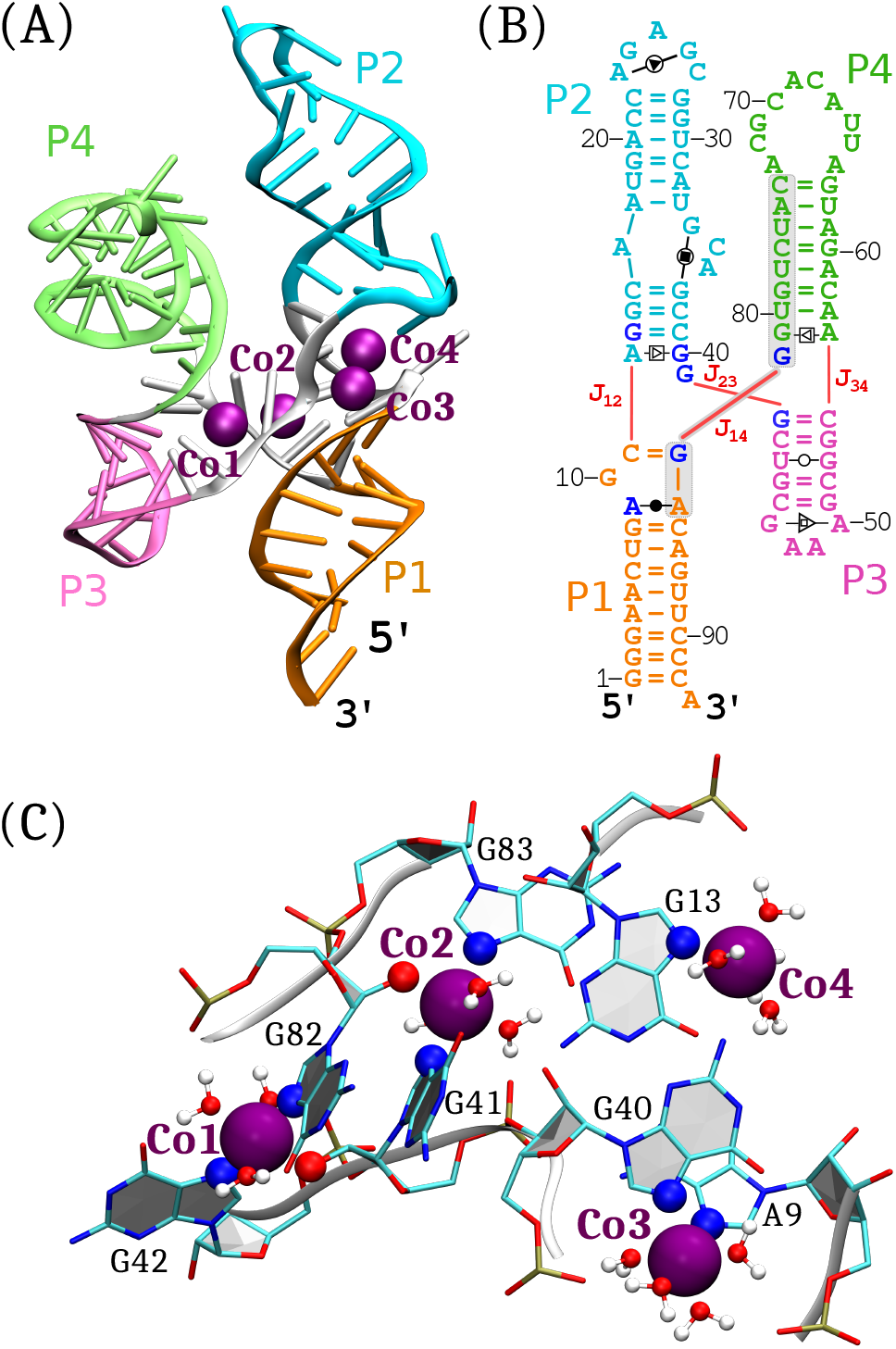
**(A)** The NRA crystal structure with the four bound Co^2+^ ions (Co1, Co2, Co3 and Co4 shown as purple spheres). The structure has four helices - P1 (orange), P2 (cyan), P3 (mauve), and P4 (green), and a four-way junction (4WJ) (gray) (PDB ID: 4RUM^12^). **(B)** The secondary structure of NRA. Nucleotides that are directly bound with Co^2+^ ions are shown in blue. Different non-canonical H-bonds are shown in black^27^ (Table S1). Junctions between two consecutive helices are shown in red. The anti-terminator or the switching strand (C74 to A84) is highlighted in gray.^12^ **(C)** The ion-binding nucleotides at the 4WJ. The base nitrogens (N7, blue spheres) and sugar oxygens (O2^*′*^, red spheres) that form IS contacts with Co^2+^ ions (purple spheres) are highlighted.

Experimentally, it is challenging to parse the role of different metal ions in riboswitch folding and metal ion binding, as multiple time scales are involved in these events. Identifying the structures of short-lived intermediate states crucial for cognate metal ion recognition in experiments is also challenging. To overcome these problems, we used molecular dynamics (MD) simulations using coarse-grained^24^ and all-atom RNA models^25,26^ to investigate riboswitch folding and Co^2+^ ion sensing in the presence of K^+^ and Mg^2+^ ions. We also performed density functional theory (DFT) calculations to understand the origin of riboswitch specificity for binding Co^2+^ ion over Mg^2+^.

Combining the results from MD simulations using coarse-grained and all-atom RNA models and DFT calculations, we show that NRA folds through an intermediate state, where divalent metal ions bind and stabilize the four-way junction (4WJ). The formation of weak non-canonical G*·*A base pairing is essential for the coaxial stacking of helices and the folding of NRA. Interestingly, divalent ions tune these base pair interactions depending on their concentration. We also show that the four crystal-bound Co^2+^ (Co1, Co2, Co3, and Co4) are not equally stable in their bound state, and their binding stabilities are correlated (Fig. 1C). Additionally, the enhanced orbital interactions between conserved guanine N7 atoms and Co^2+^ contribute to the selectivity of Co^2+^ over Mg^2+^. This work comprehensively establishes the folding energy landscape of the transcriptional NiCo riboswitch. It further illustrates the origin of selective transition metal ion binding and how it is coupled to the global conformational changes in the riboswitch that drive gene regulation.

## 2 Methods

### 2.1 Simulation Details

#### 2.1.1 Coarse-Grained (CG) MD Simulations

There are multiple RNA CG models to study various problems related to RNA.^28–35^ We used the three interaction site (TIS-RNA)^24,28^ model to study the folding of the NRA riboswitch. In the TIS-RNA model, each nucleotide is modeled using three beads, which represent the phosphate, sugar, and nucleobase groups. The initial structure for the simulations was prepared using the crystal structure of the NRA riboswitch (PDB ID: 4RUM). ^12^ The missing nucleotide U71 was added using COOT software.^36^ We performed Langevin dynamics simulations using OpenMM^37^ at temperature (*T*) 300 K using a cubic box of length 200 Å. Simulations were performed at various ion concentrations: (i) [K^+^] = 50, 100, and 150 mM. (ii) [Mg^2+^] = 2, 4, 6, and 8 mM with a fixed background [K^+^] = 150 mM. (iii) [Co^2+^] = 2, 4, and 8 mM with a fixed background [K^+^] = 150 mM and [Mg^2+^]= 8 mM. The K^+^, Mg^2+^, Co^2+^, and Cl^−^ ions were randomly inserted into the box to maintain the required concentration. The equations of motion were integrated using the LangevinIntegrator in OpenMM with a timestep, *τ* = 2 fs. The cumulative simulation time using CG models for all ion concentrations is 11 *µ*s. The hydrogen bonds and tertiary stacks used are shown in Table S1. The CG parameters for RNA and ions are given in Table S2. A detailed description of the TIS RNA model and simulation methodology is in the SI.

#### 2.1.2 All-Atom (AA) MD Simulations

We performed unbiased all-atom simulations to study selective metal ion binding to NRA using the GROMACS program (version 2018.6).^38,39^ The initial conformation for the simulations was prepared by removing the terminal phosphate group at the 5^*′*^-end and the three bound Co^2+^ ions (labeled as Co1, Co2, and Co3) from the crystal structure (PDB ID: 4RUM^12^). This system is labeled as F-NRA. We also performed AA simulations of the intermediate state (I state) populated in the CG simulations. The conformations of the I state from the CG simulations are converted to the AA description using the TIS2AA^40^ program. This system is labeled as I-NRA.

We performed additional simulations to understand the cooperative binding of ions using the crystal structure of NRA with (i) Co1, Co2, and Co3 ions bound to the NRA (labeled as NRA(1,2,3)), (ii) only Co1 and Co3 bound to NRA (labeled as NRA(1,3)), (iii) only Co2 and Co3 bound to NRA (labeled as NRA(2,3)). The simulations were performed in a cubic box of length 100 Å. The details of ion concentrations for different systems are provided in Table 1. There are multiple AA RNA and ion forcefields^41,42^ available in the literature. We used a modified AMBER ff14 forcefield (DESRES ff) for RNA^25^ and TIP4P-EW^43^ water model. The monovalent and divalent ion parameters are taken from ref.^44^ and Table 8 of ref.,^45^ respectively. Periodic boundary conditions were implemented in all directions. A detailed description of the AA simulations and justification for the chosen RNA and ion force fields are in the SI.

**Table 1:**
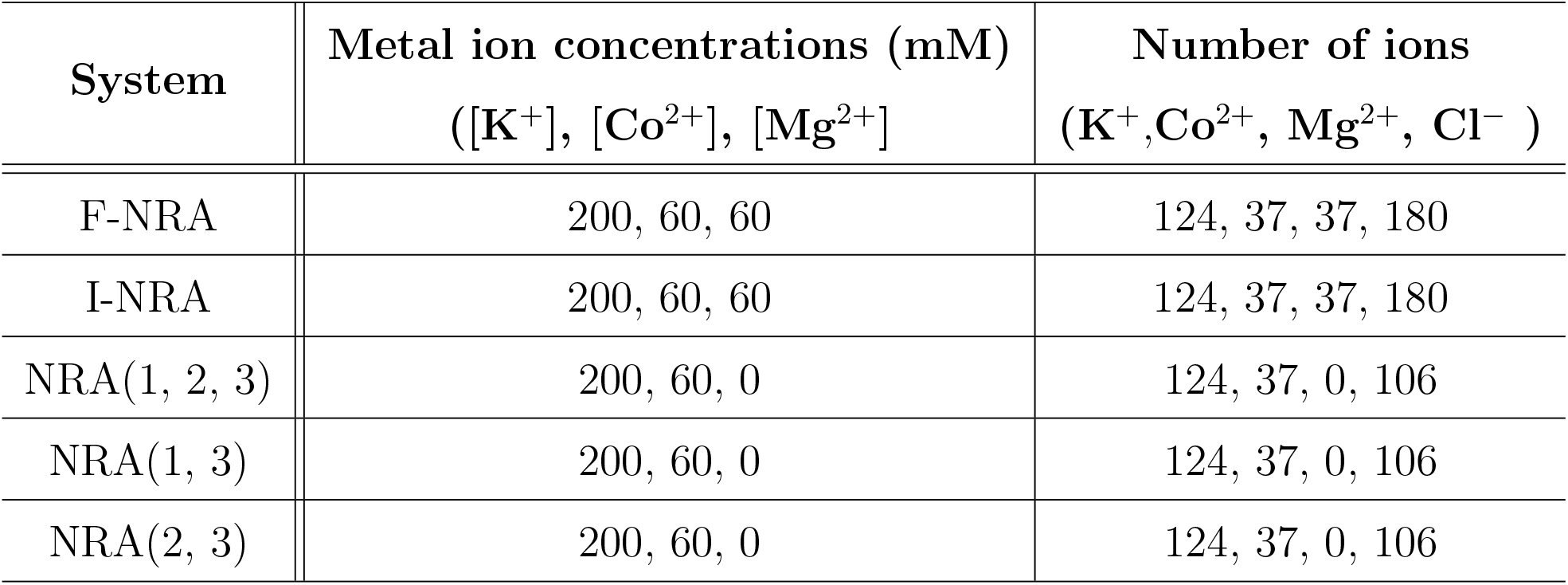
Description of AA simulation systems.

#### 2.1.3 DFT Calculations

We performed electronic structure calculations to investigate the orbital interactions between conserved guanine nucleotides in the ion-binding domain (IBD) (Table 2) and metal ions. We used Gaussian 16^46^ to perform density functional theory (DFT) calculations and geometry optimization of four complexes – Co_2_G_4_, Mg_2_G_4_, Co_1_G_2_, and Mg_1_G_2_. All stationary points were optimized using the unrestricted B3LYP hybrid density functional with Grimme’s dispersion correction and Becke-Johnson damping (GD3BJ).^47^ We used the Def2-SVP basis set for geometry optimizations and frequency calculations (the electronic energies are shown in Table S3), and Def2-TZVPP for single-point calculations. ^48^ The atomic coordinates for each complex are given in SI. The solvent effects are modeled using the self-consistent reaction field (SCRF) method applied to the density-based solvation model (SMD).^49^ To compare the interaction strengths of IBD for different metal ions, we performed Bader’s atoms in molecules (AIM)^50^ analysis using Multiwfn software^50,51^ and natural bond orbital (NBO) analysis using the NBO 3.1 program^52^ as implemented in Gaussian 16.^46^

**Table 2:** Description of nucleotides present in different sub-parts of NRA.

| Region | Nucleotides |
| --- | --- |
| Four-way-junction (4WJ) | A9-G14, C39-C43, G54-A57, and G81-A84 |
| Ion-binding-domain (IBD) | A9, G40, G41, G42, G81, and G82 |
| pocket-12 (poc12) | G41, G42, G82, and G83 |
| pocket-3 (poc3) | A9 and G40 |

### 2.2 Data Analysis

#### 2.2.1 Structural Overlap Function (*χ*)

The structural overlap function (*χ*)^53^ is computed using 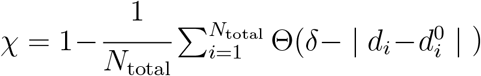, where *d*_*i*_ and 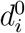 are the distances between the *i* pair of beads in a RNA conformation and NRA crystal structure. Θ is the Heaviside step function, and *δ* = 5 Å. *N*_*total*_ (= 35245) is the total number of pairs of beads in the TIS model of NRA. Here, only pairs of beads that are at least 3 nucleotides separated from each other are used in the calculation.

#### 2.2.2 Coordination Number

The coordination number between divalent ions (Co^2+^ or Mg^2+^) and the IBD nucleotides is calculated using the equations,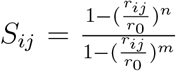 and *CN* =∑ _*i*∈*A*_ ∑ _*j*∈*B*_ *S*_*ij*_ where group A consists of all the M^2+^ ions (M = Co or Mg). Group B consists of the phosphate beads of IBD nucleotides to compute 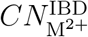 (Table 2). For computing 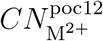 and 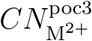, group B consists of all the atoms of poc12 and poc3 nucleotides (Table 2). *r*_0_ (= 5.0 Å) is the cutoff distance and *r*_*ij*_ is the distance between atoms *i* and *j* from groups *A* and *B*, respectively. The steepness of the switching function is modulated by the *n* (= 6) and *m* (= 12).

#### 2.2.3 Fraction of Tertiary Contacts

The fraction of native tertiary contacts (TC) between a particular set of nucleotides (*λ*) in a conformation *i*, is computed using 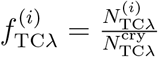 where 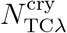 and 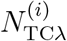 are the number of native contacts present between the nucleotides belonging to set *λ* in the crystal structure and the *i*^*th*^ conformation, respectively. ^54,55^

#### 2.2.4 Effect of Divalent Ions on Tertiary Contact Formation

To probe whether the binding of divalent ions (M^2+^, M = Mg and Co) to the nucleotides belonging to the junction loops (J_23_ and J_14_) (Fig. 1B) affect the tertiary contact formation between these loops and leads to the folding of the NRA, we calculated the free-energy change between these two states 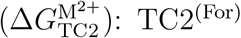 and TC2^(Rup)^. If the tertiary contacts between the junction loops J_23_ and J_14_ (labeled as TC2) are formed i.e. the fraction of native contacts in TC2, 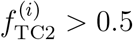 and at least one divalent ion M^2+^ is bound to both the junctions, then we designate the state as TC2^(For)^ (Fig. S1). If 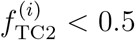 and no divalent ion M^2+^ is bound to both the junctions, then the state is TC2^(Rup)^. The free energy difference 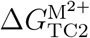 is given by 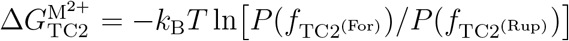 where 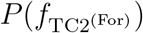 and 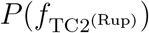 are the probability of observing states TC2^(For)^ and TC2^(Rup)^, respectively.

#### 2.2.5 Local Ion Concentration Around RNA

Local concentration of a specific type of ion *j*, around a specific RNA atom is given by,^24^ 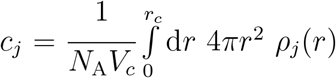, where *ρ*_*j*_(*r*) is the number density of ion *j* at a distance *r* from the considered RNA atom, *V*_*c*_ is the spherical volume with a cutoff radius *r*_*c*_ (= 5 Å), and *N*_A_ is the Avogadro number. See Data Analysis in SI for details.

## 3 Results

### 3.1 Divalent Metal Ions Drive NRA Folding

We performed MD simulations using the coarse-grained TIS model of RNA (see Methods) to investigate the effects of monovalent ([K^+^]) and divalent ([Mg^2+^] and [Co^2+^]) ion concentration on NRA folding. ^12,20^ To quantify the structural changes induced by the ions, we projected the free energy surface (FES) onto the structural overlap function, ^53^ *χ*, for different cation concentrations. Here, *χ* = 0 and 1 for a conformation indicates that its structure is highly similar and dissimilar to the crystal structure (PDB ID: 4RUM ^12^), respectively. In the absence of Mg^2+^, the FES showed only a single minimum at *χ* ≈ 0.57 corresponding to the unfolded state (denoted as U state) as we varied [K^+^] from 50 to 150 mM indicating that K^+^ alone cannot induce NRA folding (Fig. S2). Subsequently, we studied NRA folding by varying [Mg^2+^] from 2 to 8 mM while keeping the background concentration of K^+^ ion constant at 150 mM (physiological concentration) (Fig. 2A). At low [Mg^2+^] (= 2 mM), the unfolded U state (*χ* ≈ 0.54) is the dominantly populated state. In contrast, at higher [Mg^2+^] (4 to 8 mM), two additional states, an intermediate state (I state) (0.37 < χ ≤ 0.5) and the folded state (F state) (*χ* ≤ 0.37) are populated. Multiple transitions between the F, I, and U states are observed (Fig. 2B).

**Figure 2.**
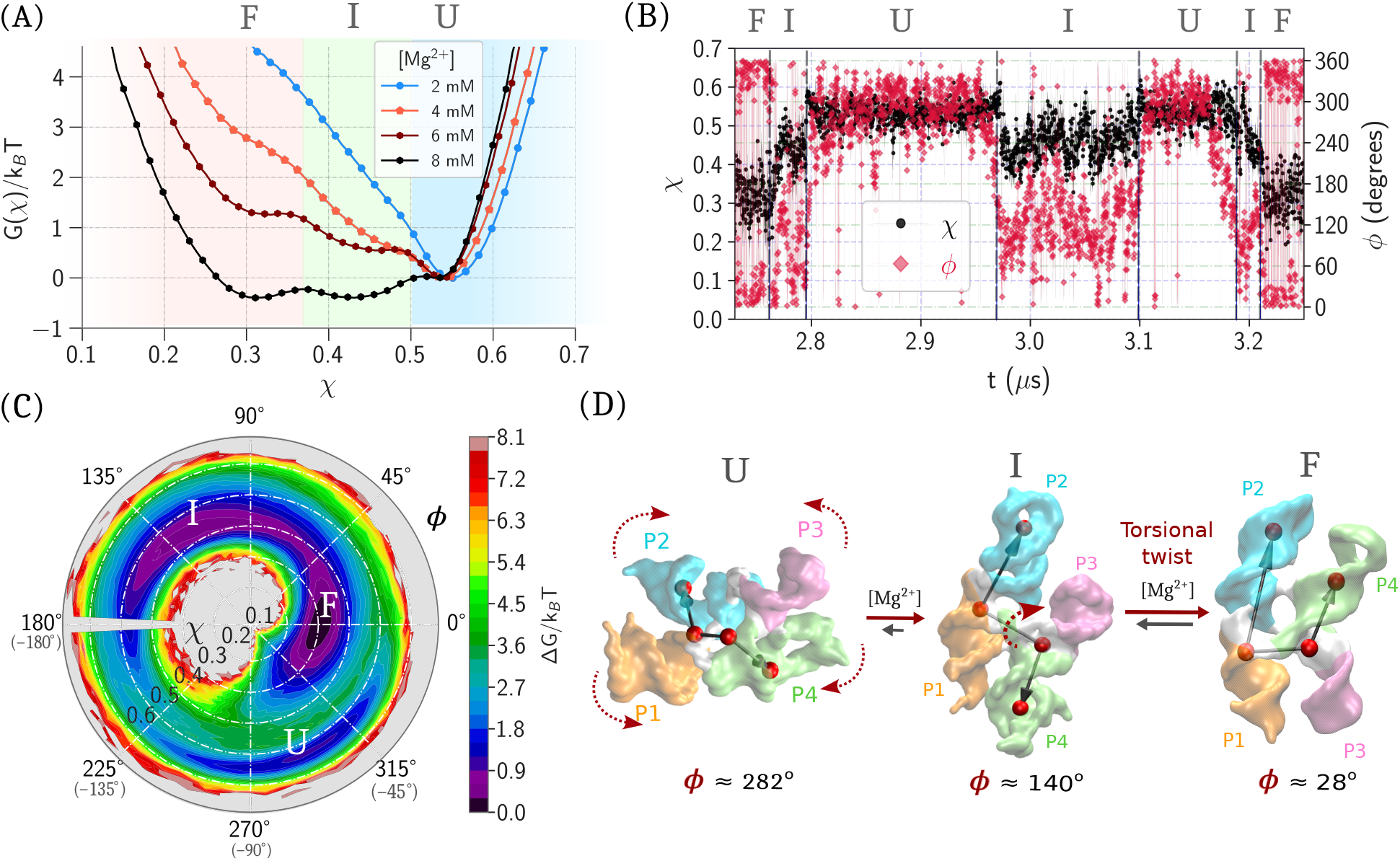
**(A)** FES projected onto χ for different [Mg^2+^] at 300 K. The folded (F), intermediate (I), and unfolded (U) states are highlighted in pink, green, and sky-blue shaded regions, respectively. **(B)** Simulation trajectory showing the evolution of χ and ϕ as a function of time ([Mg^2+^] = 6 mM). The vertical dashed lines (black) are a guide to the eye showing the transitions between the three states (F, I, and U states). **(C)** 2D polar FES (pFES) projected onto χ (along the radius) and ϕ (along the azimuth) for [Mg^2+^] = 8 mM. **(D)** Transition from U → I → F shown by representative structures and torsion angles (ϕ) between P2 and P4 helices are shown for states F, I, and U (See Movie S1).

The smFRET experiments^20^ suggested the existence of three thermodynamic states in the NRA folding process by probing the dynamics of P2 and P4 helices. The experiments showed that at high [Na^+^] (≈ 500 mM) and low [Mg^2+^] (≈ 2 mM), an intermediate state is populated, which is in agreement with our observations. Interestingly, our CG simulations show that at [Mg^2+^] = 8 mM, the F and I states are more stable than the U state, suggesting that beyond a critical threshold Mg^2+^ concentration ([Mg^2+^] ≈ 6 mM), the I and F states are global minima (Fig. 2A).

To further probe whether the CG TIS RNA model of NRA can distinguish the Mg^2+^ and Co^2+^ ions in a mixture, we performed additional simulations by fixing [Mg^2+^] at 8 mM and [K^+^] at 150 mM, and varied [Co^2+^] from 2 to 8 mM. The FES projected onto *χ* shows that increasing the divalent ion concentration by adding Co^2+^ stabilized the F state, making it the global minimum in agreement with the smFRET experiments^20^(Fig. S3). In the later sections, we show that the CG TIS model has limitations in distinguishing between the Mg^2+^ and Co^2+^ ions in a mixture.

The simulation trajectories show that during transitions between the U, I, and F states, NRA undergoes diverse conformational changes by reorganizing the orientation of the four helices (P1 to P4). Since *χ* cannot quantify the relative orientations of the helices in different states, we used the torsion angle, *ϕ*, between P2 and P4 helices to distinguish different states. *ϕ* is computed using the center of mass positions of the phosphate backbone beads of nucleotides present at P2 and P4 helix head and P1|P2 and P3|P4 junction (Fig. S4). The order parameter *ϕ* complements *χ* in distinguishing the structural features of different NRA states (Fig. 2B). The 2D polar FES (pFES) projected onto *χ* (along the radial axis) and *ϕ* (along the azimuthal axis) shows three basins (F, I, and U states) with different *χ* and *ϕ* values (Fig. 2C).

In the U state (*χ ≥* 0.5), P2 and P4 point along different directions as *ϕ* spans a broad range (225^*°*^ < *ϕ* < 335^*°*^) (Fig. 2D). In the I state (0.37 ≤ *χ* < 0.50), P2 and P4 are approximately anti-parallel (80^*°*^ < *ϕ* < 190^*°*^), suggesting a significant structural rearrangement during the U to I transition. The comparison between the structures from the U and I basins in the pFES suggests that NRA undergoes a coaxial stacking rearrangement between helices while transitioning from U to I states (Fig. 2D and Movie S1).^56,57^ In the F state (*χ* < 0.37), P2 and P4 helices become approximately parallel (−55^*°*^ < *ϕ* < 45^*°*^) as observed in the crystal structure. The comparison between conformations in the I and F basins from the pFES shows that NRA only undergoes a simple torsional twist about the 4WJ in the I to F transition (Fig. 2D, and Movie S1). The small barrier heights (∼ *k*_B_*T*) (Fig 2C) are due to the absence of tertiary contacts between the helices (PDB ID: 4RUM^12^).

### 3.2 Tertiary Contacts in the 4WJ Drive the Formation of Native-like Helical Stacking

Since the four helices in the NRA interact only through the four-way junction (4WJ) (Table 2), their structural dynamics must be strongly correlated to changes in the 4WJ. The interactions between nucleotides in the 4WJ (Table 2) aided by the metal ions (Co^2+^, Ni^2+^ and Fe^2+^) should drive the transitions between different states (U ⇌ I ⇌ F).^12,20^ Contact maps and pair-distance matrices (PDM) show that 4WJ in the F state is compact due to the crossed topology that brings the helix P1 near P3, and P2 near P4 stabilizing five of the six tertiary contacts (TC1–TC6, except TC4) (Fig. S5, S6 and S7, left panels). The conformations in state F with coaxial stacking of helices P1|P2 and P3|P4 resemble the native crystal structure (Fig. S8). In state I, the contact probabilities of the tertiary contacts (TC1–TC6, except TC4) decrease due to the untwisting of the 4WJ retaining the coaxial stacking of helices (P1|P2 and P3|P4) (Fig. S5, S6 and S7, middle panels). The untwisted 4WJ resembles a DNA Holiday junction.^58^ In state U, there is a further loss of 4WJ contacts leading to a widely open 4WJ. In these conformations, there is a new coaxial stacking arrangement of helices (P1|P4 and P2|P3) (Fig. S5, S6 and S7, right panels). In particular, TC4 contacts (A9-A12 and G54-A57) are observed in this state, which were absent in states F and I. The conformations in state U look like a distorted X-like structure, indicating large-scale conformational transitions from state I. The transitions between F, I, and U are marked by changes in coaxial stacking between helices and tertiary contacts within the 4WJ.

### 3.3 Terminal Non-canonical G*·*A Base Pairs in Helices Control Loop Lengths Critical for Coaxial Stacking of Helices

The coaxially stacked helices are common to all three states observed in NRA folding. The quality of coaxial stacking between consecutive helices depends on the number of nucleotides in the loop RNA strand connecting the helices. The loop RNA strand that connects two consecutive helices *i* and *j* forms the junction, J_*ij*_.^59^ A smaller RNA strand constituting the junction leads to better coaxial stacking. ^59^ In the crystal NRA structure, the number of nucleotides in the RNA strands constituting junctions J_12_, J_23_, J_34_, and J_14_ are 0, 1, 0, and 1, respectively (Fig. 3A). However, the junction length can be dynamic depending on the stability of the base pairs present near the junctions at the helix terminals that connect two consecutive junctions. The terminal base pairs in helices P1, P2, P3 and P4 are G83*·*C11, G40*·*A12, G42*·*C55, and G81*·*A56, respectively (Fig. 3A and S9). To probe the effect of these terminal base pairs on coaxial stacking of helices, we calculated the probability of base pair formation using the equation,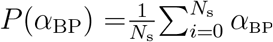, where *s* = F, I, or U states, *N*_*s*_ = total number of conformations in state *s* and *α*_BP_ = 1 if there is a hydrogen bond between the bases and 0 if the hydrogen bond is absent. A hydrogen bond is present between the bases if its hydrogen bond energy *U*_HB_ < −*k*_B_*T* (Eq. S4).

**Figure 3.**
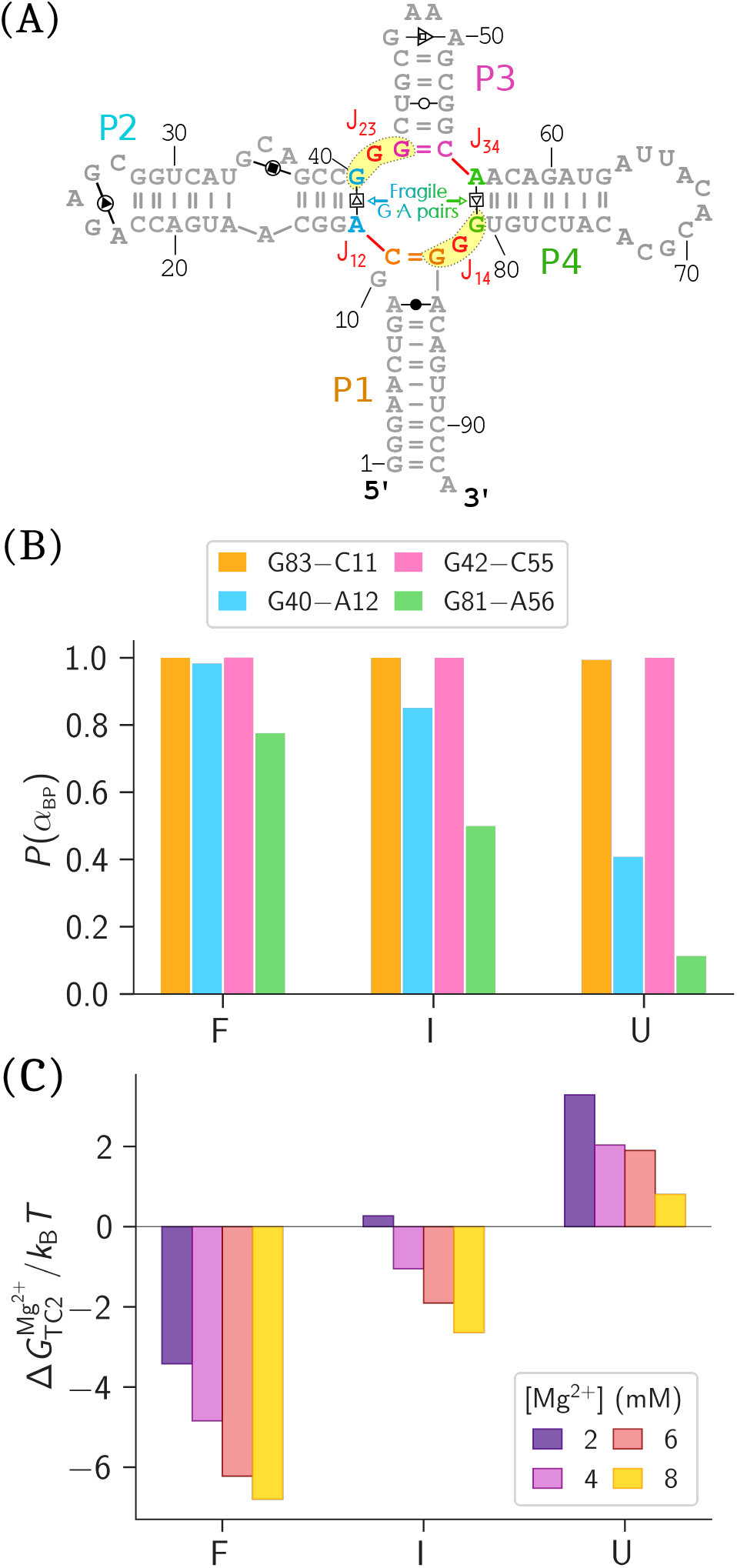
**(A)** The terminal base pairs of the helices P1 (G83·C11: orange), P2 (G40·A12: cyan), P3 (G42·C55: mauve), and P4 (G81·A56: lime) in NRA secondary structure corresponding to states F and I. The junction loop (J_*ij*_) nucleotides (red), the nucleotides corresponding to TC2 (yellow shade), and non-canonical H-bonds(black) are highlighted. **(B)** The probability of terminal base pair formation, P(α_BP_), is plotted for states F, I, and U. **(C)** The 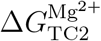(see Data analysis) with varying [Mg^2+^] for states F, I, and U.

The terminal canonical G*·*C base pair hydrogen bonds of helices P1 (G83*·*C11) and P3 (G42*·*C55) remain intact (*P*(*α*_BP_) ≈ 1) in all states (U, I and F) (Fig. 3B). In contrast, the stability of the terminal non-canonical G*·*A base pair hydrogen bonds in P2 (G40*·*A12) and P4 (G81*·*A56) helices depends on the state of NRA (Fig. 3B). These base pairs are least stable in state U, resulting in the elongation of the loops at the junctions in state U (Fig. 3B and S10). In states I and F, the stability of the G*·*A pairs is high, resulting in the shortening of the junction loop regions (Fig. 3A,B), enhancing the coaxial stacking of the helices, which strongly depends on the loop length. Generally, Watson-Crick G*·*C base pairs (G83*·*C11 and G42*·*C55) possess three hydrogen bonds and are more stable than the sheared non-canonical G*·*A base pairs (G40-A12 and G81-A56), which possess only two hydrogen bonds each.^59–61^ The shortening of all junction lengths (J_12_, J_23_, J_34_, and J_14_) due to the formation of the terminal G*·*A base pairs in helices P2 (G40*·*A12) and P4 (G81*·*A56) will enhance the helical stacking propensity (Fig. 3A,B). However, since all the junction lengths are shortened, coaxial stacking of the two possible combinations of consecutive helices (P1|P2:P3|P4 or P1|P4:P2|P3) is possible. The following questions remain: (1) Why is the P1|P2:P3|P4 coaxial stacking arrangement preferred in the I and F states of NRA? (2) What drives the formation of the terminal G*·*A base pairs in P2 and P4 helices?

### 3.4 Divalent Ion-binding Drives Specific Coaxial Stacking of Helices

The crystal structure shows that most of the guanine nucleotides involved in Co^2+^ binding belong to or are adjacent to J_23_ (G40, G41, and G42) and J_14_ (G81, G82, and G83) junction loops. We hypothesized that the interaction of the divalent metal ions with the guanine nucleotides facilitates the formation of terminal G*·*A base pairs in helices P2 (G40*·*A12) and P4 (G81*·*A56) and the specific coaxial stacking of P1|P2 and P3|P4 helices in the I and F states.

To verify the hypothesis, we calculated the effect of divalent ion (M^2+^, where M = Mg and Co) binding on the stability of the TC2 tertiary contacts present in the 4WJ by computing the free energy 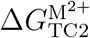 (see Data Analysis). TC2 tertiary contacts involve contacts between nucleotides present in the junctions J_23_ (G40, G41, and G42) and J_14_ (G81, G82, and G83) (Fig. S5). The negative (positive) value of 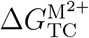 implies that the specific binding of M^2+^ stabilizes (destabilizes) the formation of tertiary contacts between J_23_ and J_14_ (Fig. 3C). In the state U, 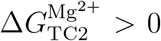 for all [Mg^2+^], indicating that TC2 ruptured state with divalent ions not bound to the J_23_ and J_14_ nucleotides is dominantly populated. In contrast, in state I, 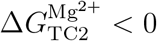 for [Mg^2+^] > 2 mM, showing that TC2 contacts are formed with divalent ions bound to the J_23_ and J_14_ nucleotides and their stability increased with [Mg^2+^] (Fig. 3C). In state F, 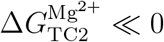, suggesting that TC2 contacts are stable for all [Mg^2+^] and also their stability increased with [Mg^2+^] (Fig. 3C). We found a similar trend in 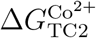 when Co^2+^ is used as a divalent ion (Fig. S11).

The analysis using 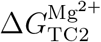 supports the hypothesis that divalent ion binding to the nucleotides of J_23_ and J_14_ helps in structural rearrangements that pull these junctions closer to each other to form TC2 contacts in I and F states, which facilitates coaxial stacking. In this process, the terminal G*·*A base pairs also form hydrogen bonds in helices P2 and P4, providing additional stability.

### 3.5 Structure of the Ion Binding Domain (IBD) Modulates Ion-binding Affinity

The equilibrium between different NRA states (U ⇌ I ⇌ F) is dependent on the divalent ion concentration (Fig. 2A and S3). However, not all nucleotides are directly involved in the Co^2+^ sensing activity (Fig. 1C). To identify the nucleotides sensitive to divalent ion binding and their role in stabilizing the three distinct states (U, I, and F), we computed the local concentration of Mg^2+^ 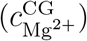 around each phosphate backbone bead for the conformations in the three states (see Data Analysis) (Fig. 4A). Unlike the state U, the states I and F showed significant 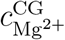 around the nucleotides A9, G40, G41, G42, G81, and G82, which are interestingly the ion binding domain (IBD) nucleotides (Fig. 1C and 4A). The spatial distribution of Mg^2+^ around the NRA backbone also showed that the ions preferably bind around the IBD region (Fig. 4B).

**Figure 4.**
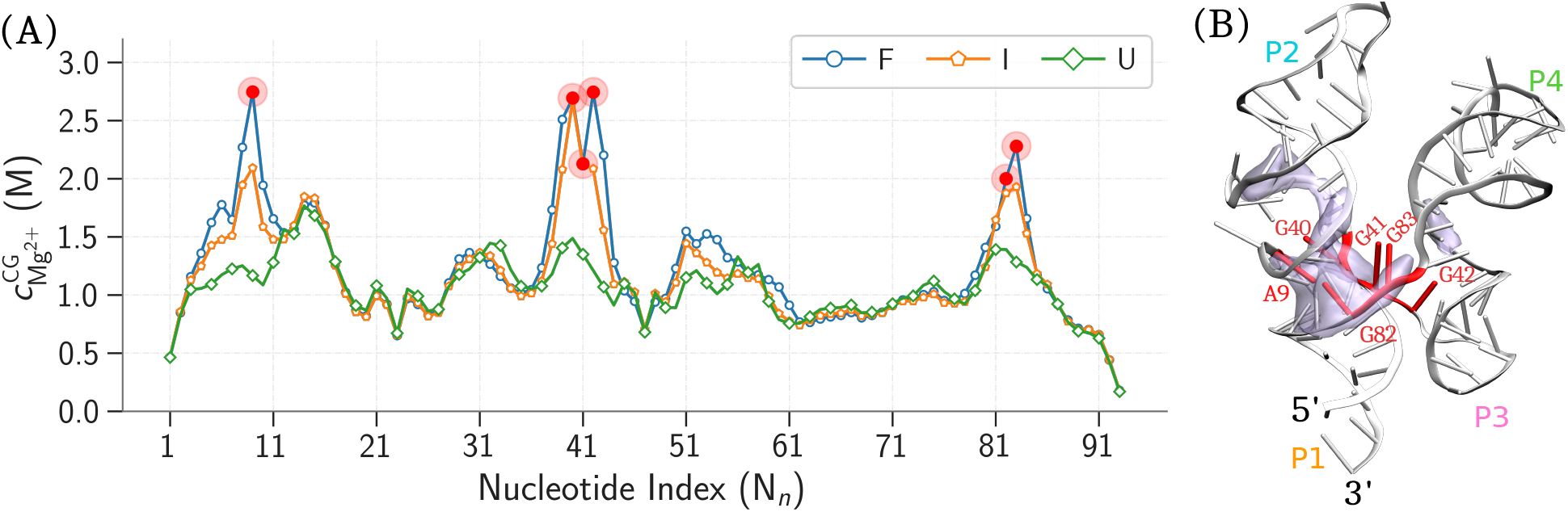
**(A)** Average local Mg^2+^ ion concentration,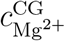, around the phosphate beads for states F (blue), I (orange), and U (green). IBD nucleotides (A9, G40, G41, G42, G82, and G83) are pointed out as red circles. **(B)** NRA folded structure is shown in grey, and IBD nucleotides are shown in red. The surface plot in violet color shows the spatial distribution of Mg^2+^ ions around RNA. The isosurface corresponding to ISO value 0.008 is plotted for the high-charge-density region^62^ (see SI).

To probe the Mg^2+^-induced IBD formation, we calculated the average structural overlap parameter for the IBD (⟨*χ*^IBD^⟩) (Fig. S12). To monitor the ion binding affinity of the IBD, we computed the average number of Mg^2+^ ions coordinated to the IBD nucleotides (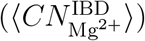) (see Data Analysis) (Fig. S12). The decrease in ⟨*χ*^IBD^⟩ with increasing [Mg^2+^] indicates that higher [Mg^2+^] promotes IBD formation. Correspondingly, 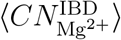 increased gradually with increasing [Mg^2+^], highlighting the importance of Mg^2+^ binding to stabilize IBD (Fig. S12).

In the three states (F, I, and U), 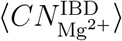 increased with [Mg^2+^] due to the availability of more ions (Fig. S13A). In contrast, for a fixed [Mg^2+^], 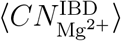 shows a gradual increase from U →I →F (Fig. 5 and S13A) as it should be correlated to the IBD cavity size, which determines the effective interactions between the IBD beads and ions. As expected, in all three individual states (F, I, and U), the average radius of gyration of IBD, 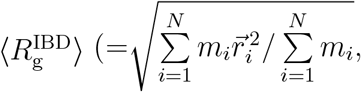, where *m*_*i*_ is the mass of bead *i* and 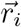 is the distance of bead *i* from the center of mass of the IBD beads) decreased with increasing [Mg^2+^] (Fig. 5 and S13B). For example, at [Mg^2+^] = 6 mM, transitioning from U → I → F, 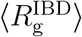 decreased from ≈ 12.2 Å to 7.9 Å, implying compaction of the IBD cavity volume by approximately three-fold (Fig. 5). Previously, we observed the change in the number of nucleotides associated with the lengths of the junction loops (J_*ij*_), which also contributed to the change in IBD size (Fig. 3A-C).

**Figure 5.**
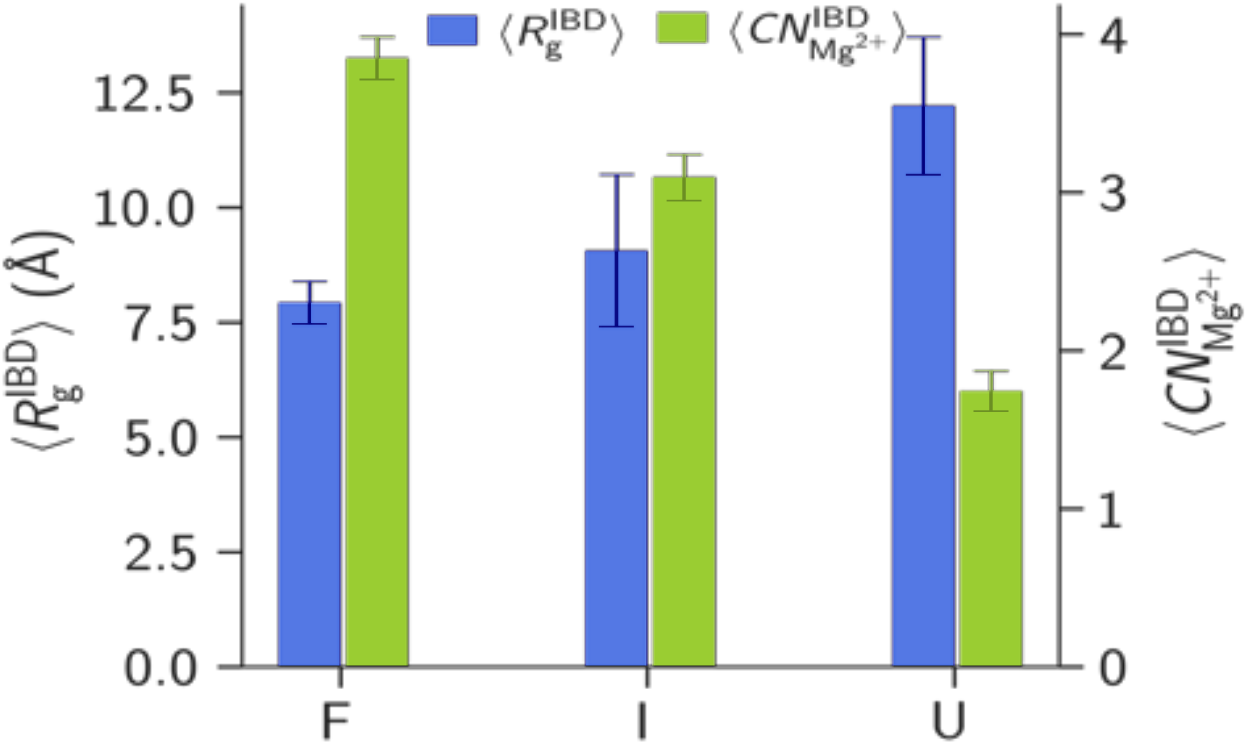
At [Mg^2+^] = 6 mM, the average coordination number associated with the IBD, 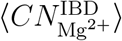, is shown in green and the average radius of gyration of IBD, 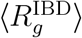, are shown in blue for states F, I, and U.

In addition to the average cavity size, the flexibility of IBD is also important in determining its binding affinity. We quantified the flexibility by measuring the fluctuation in 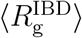, which indicates that the IBD of state F 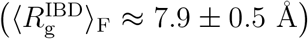 is rigid and compact, resulting in the highest 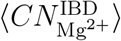 (Fig. 5, S13, and S14). Unlike state F, the IBD in state U is wide open and flexible 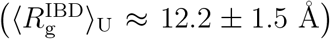. However, the IBD in state I 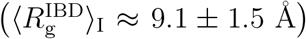 is slightly larger compared to F but has flexibility comparable to state U. The moderate ion-binding capacity of state I is therefore attributed to the combined effects of moderate cavity size and high flexibility of IBD. Similarly for Co^2+^, we observed similar trends in 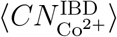 and 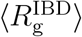 (Fig. S15A,B, S16). In conclusion, not only state F but also state I is capable of binding ions, and the global folding of IBD is cooperatively coupled to the ion binding affinity of NRA.

### 3.6 IBD Structure Creates an Anionic Cavity that Attracts Divalent Metal Ions

In the CG model, Co^2+^ ions (*R* = 1.288 Å) have a higher charge-to-size ratio compared to Mg^2+^ ions (*R* = 1.353 Å) (Table S2). In addition to the electrostatic interaction, the ions interact with the RNA beads only through excluded volume interaction (see SI). From the CG simulations, we observed that state F is more stabilized compared to states I and U, as we increased [Co^2+^] keeping [Mg^2+^] and [K^+^] fixed at 8 mM and 150 mM, respectively (Fig. S3). We also observed that when [Mg^2+^] = [Co^2+^] = 8 mM, Co^2+^ ions are more condensed in the IBD compared to Mg^2+^ ions as it has a higher charge-to-size ratio. However, we did not observe any unique patterns for 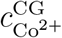 over 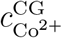 indicating a preferred binding specificity for one particular divalent ion over the other (Fig. S17 and S18). Co^2+^ condenses relatively more compared to Mg^2+^ due to its higher charge-to-size ratio.

From the CG simulations, we learn that as the NRA folds to the I state, the partially folded 4WJ forms the IBD with an anionic cavity that has a negative electrostatic potential surface, which has a strong affinity for the binding of positively charged metal ions (Fig. S19 and S18). The anionic cavity is primarily due to the arrangement of the phosphate groups, as these are the only CG beads that are charged in the CG TIS RNA model. In the CG simulations, the IBD has no strong specificity for binding Co^2+^ compared to Mg^2+^. We attribute this to the absence of explicit water molecules as the crystal structure^12^ shows ions bound to the IBD using both IS and OS water-mediated interactions (Fig. 1A-C). The other drawback could be that Co^2+^ is a transition metal ion that can form coordination complexes with the IBD nucleotides, which the CG simulations cannot model. To overcome the limitations of the CG TIS model where water is not explicitly present and to incorporate the water-mediated interactions between RNA heavy electronegative atoms and metal ions, we performed unbiased all-atom (AA) simulations to explore the role of water in the specificity of Co^2+^ ion binding to the IBD over Mg^2+^ (see Methods).

From the AA simulation trajectories of the native NRA (F-NRA) system (see Methods and Table 1), we computed the local ion concentration of Co^2+^ 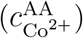 and Mg^2+^ 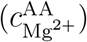 around the phosphate backbone atoms (O1P and O2P) of NRA (see Data Analysis in Methods) (Fig. S20 and S21). On comparing local Co^2+^ ion concentration around NRA computed from CG 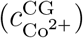and AA 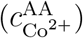 simulations, we find that the local ion concentration pattern exhibited remarkable similarity for the IBD nucleotides (A9, G40, G41, G42, G82, and G83) (Fig S21). This further supports that the ion-binding cavity in the IBD is anionic due to the specific arrangement of negatively charged phosphate groups aided by the local structure of the 4WJ that drives the attraction of the metal ions to these nucleotides.

### 3.7 Selective OS Coordinated Binding of Co^2+^ in the IBD Pocket

The local ion concentration 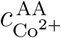 shows that Co^2+^ ions are more condensed compared to Mg^2+^ near nucleotides G8 to G10, C39 to C43 and G81 to G82 in the vicinity of IBD (Fig. 6A). In the crystal structure,^12^ Co^2+^ ions Co1 and Co2 interact with G41, G42, G82, G83, and Co3 interacts with A9 and G40 (Fig. 1C). The higher local concentration of Co^2+^ over Mg^2+^ in the IBD region in the AA simulations is due to its higher charge-to-size ratio and also from the van der Waals interaction modeled through the Lennard-Jones (LJ) potential, which is absent in the CG simulations where only excluded volume of ions is present. The LJ parameters of ions in the AA simulations (Table 8 in Ref.^45^) are optimized to reasonably reproduce the ion hydration-free energies (HFE) and ion-oxygen distance (IOD).

**Figure 6.**
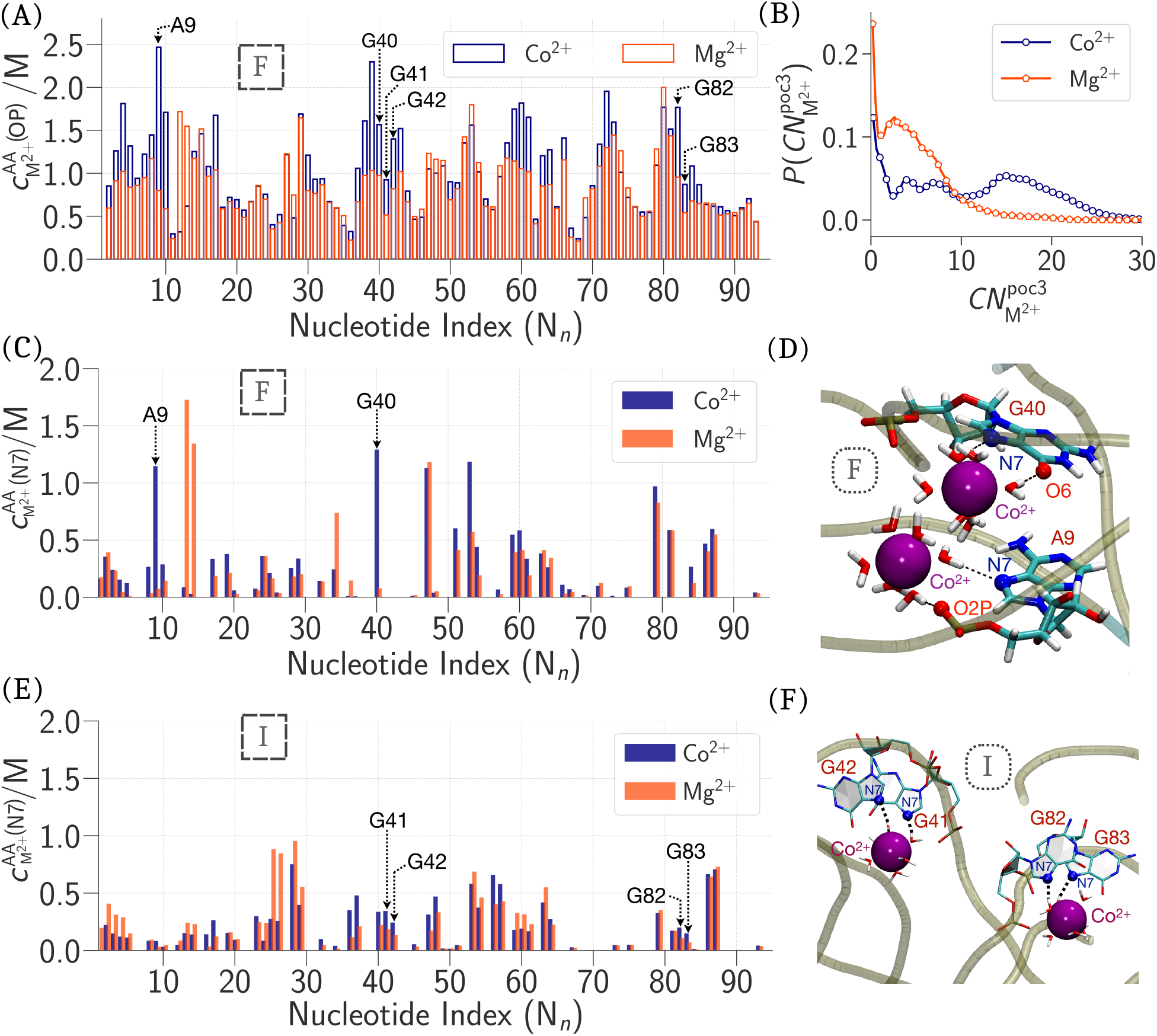
**(A)** Local ion concentration (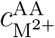, in M units) of both Co^2+^ (blue) and Mg^2+^ (orange) ions around phosphate oxygen atoms (OP) of all the nucleotides (N_*n*_). We highlighted IBD nucleotides (A9, G40, G41, G42, G82, G83) for F-NRA from AA simulation. **(B)** Probability distribution plot (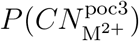) of coordination number 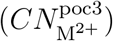 between poc3 nucleotides (A9, G40) and each divalent metal ion (Co^2+^ (blue) and Mg^2+^ (orange)) for F-NRA. **(C)** Local ion concentration (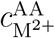, in M units) of both Co^2+^ (blue) and Mg^2+^ (orange) ions around N7 atoms of all the nucleotides (N_*n*_) where we highlight poc3 nucleotides for F-NRA from AA simulation. **(D)** Represenatative snapshot of OS interactions (dotted lines) between Co^2+^ and N7 atoms of poc3 nucleotides from AA simulation of F-NRA. **(E)** Local ion concentration (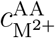, in M units) of both Co^2+^ (blue) and Mg^2+^ (orange) ions around N7 atoms of all the nucleotides (N_*n*_) where we highlight poc12 nucleotides (G41, G42, G82, G83) for I-NRA from AA simulation. **(F)** A representative snapshot of OS interactions (dotted lines) between Co^2+^ and N7 atoms of poc12 nucleotides from AA simulation of I-NRA.

To further gain insights into the interaction of Co^2+^ and Mg^2+^ with the IBD nucleotides, we computed the coordination number of these ions with specific IBD nucleotides. The crystal structure shows that four Co^2+^ ions are bound to the IBD via IS mode. Three (Co1, Co2 and Co3) out of the four Co^2+^ ions interact with common nucleotides (A9, G40, G41, G42, G82, G83) in IBD (Fig. 1B,C). Experiments^12^ report that the binding site of the fourth Co^2+^ ion (Co4) lacks selectivity. Therefore, we only focused on the selectivity of binding pockets where Co1, Co2, and Co3 ions bind and coordinate with the IBD nucleotides (A9, G40, G41, G42, G82, G83). In the crystal structure, Co1 and Co2 ions are IS-coordinated to the nucleotides G41, G42, G82, and G83 (poc12), and the Co3 ion forms IS contacts with G40 and A9 (poc3) (Fig. 1C, Table 2). To understand the origin of the ion selectivity, we computed two coordination numbers: (i) coordination number, 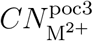, between divalent ions (M = Co or Mg) and poc3 nucleotides, (ii) coordination number 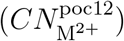 between divalent ions and poc12 nucleotides (G41, G42, G82, and G83) (See Data Analysis).

The probability distribution of 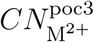 shows that at 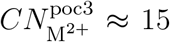 there is a peak for Co^2+^ ions that is not present for Mg^2+^ ions (Fig. 6B). We also observed that the 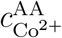 values around the N7 atoms are significantly higher for nucleotides A9 and G40 compared to the 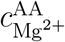 values. This was determined by calculating the local ion concentration 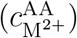 for both metal ions surrounding the N7 atoms of the poc3 nucleotides (Fig. 6C). This suggests that the high specificity for Co^2+^ binding over Mg^2+^ can be attributed to the strong water-mediated OS interactions between the N7 atoms in A9 and G40 nucleotides and the Co^2+^ ions (Fig. 6D). These observations are also consistent with experimental findings regarding the preferred interactions between Co^2+^ ions and N7 atoms.^12^ The Co^2+^ ions also interact with the highly negatively charged phosphate oxygen atoms (O2P) of poc3 nucleotides through OS coordination mode (Fig. 6D and S22) as the binding pocket is large enough to accommodate water-solvated Co^2+^ ions. The radial distribution function, *g*(*r*), for water around ion, also confirms that the size of hydrated Co^2+^ ions (≈ 1.96 Å) is smaller compared to hydrated Mg^2+^ (≈ 2.04 Å)^63^ (Fig. S23). The larger size of hydrated Mg^2+^ ions could also contribute to their lower occupancy in the binding pocket compared to the Co^2+^ ions.

To investigate the selectivity of metal ions at the poc12 nucleotides, we computed the probability distributions of 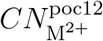. We found a peak for Co^2+^ ions at 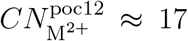, which is absent for Mg^2+^ ions (Fig. S24). Therefore, the poc12 nucleotides in IBD are also more likely to interact with Co^2+^ ions compared to Mg^2+^ ions. However, we did not observe significant local ion concentration of both the metal ions 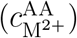 near the N7 atoms of the poc12 nucleotides (Fig. 6C). To conclude, we did not observe any IS/OS interactions between Co^2+^ ions and the N7 atoms of guanine nucleotides (G41, G42, G82, G83) in poc12 (Fig. 6C). This is because the binding pockets formed by the four nucleotides (G41, G42, G82, G83) in the folded state are compact due to the twisted topology of the 4WJ, and they cannot accommodate the solvated divalent ions. We also did not observe any OS-to-IS transitions^63^ as the energy barrier associated with this transition is high (≈ 12 kcal mol^−1^), and it is rare to observe these transitions in unbiased all-atom simulations on a time scale of *µ*s.

### 3.8 Co^2+^ Ions Bind in the Intermediate State and Stabilize the Folded State

In the AA simulations spawned using the crystal structure as the initial conformation but with all the bound Co^2+^ removed (F-NRA, Table 1), we observed that the Co^2+^ and Mg^2+^ ions in solution neither interact in IS nor OS mode with the N7 atoms of poc12 guanine nucleotides (G41, G42, G82, G83) as these nucleotides are buried and are inaccessible for the hydrated ions due to the twisted topology of the 4WJ (Fig. 6C). This is also supported by the radial distribution functions, *g*(*r*), between the divalent metal ions (Co^2+^ and Mg^2+^) and the N7 atoms of poc12 guanine nucleotides, which showed no peak at the OS or IS coordination distances (*r*_IS_ ≈ 2.0 Å and *r*_OS_ ≈ 4.2 Å) (Fig. S25). We only observed OS interactions between the metal ions and N7 atoms poc3 nucleotides (A9, G40) (Fig. 6C,D). The CG simulations showed that I state is capable of binding metal ions (Co^2+^ and Mg^2+^) in the IBD domain (Fig. 4A and 5). We took the state I conformations obtained from the CG simulations and used them as initial conformations for AA simulations (Table 1, see SI) in the presence of Co^2+^ and Mg^2+^ to probe if poc12 can preferentially bind to Co^2+^ ions (I-NRA, Table 1). We computed the local ion concentration 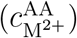 around the N7 atoms of nucleotides to investigate the ion condensation in the I state (Fig. 6E). The *g*(*r*) between the divalent metal ions (Co^2+^ and Mg^2+^) and the N7 atoms of poc12 guanine nucleotides show that metal ions interact with the N7 atoms of poc12 nucleotides predominately via OS coordination in the I state (Fig. S25). We observed that in the I state, the IBD domain is more open compared to the F state, allowing the preferential binding of hydrated Co^2+^ and Mg^2+^ ions with the N7 atoms of the poc12 nucleotides (Fig. 6F and S26). The data also shows that Co^2+^ ions condense more near the N7 atoms of poc12 nucleotides compared to Mg^2+^ ions (Fig. 6E). We infer that the Co^2+^ ions (Co1 and Co2) bind in the I state as the binding pocket is inaccessible to the ions in the F state.

The local ion concentration 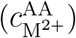 around the N7 atoms in the I state also show significant peaks for Co^2+^ ions at A56 and A57 nucleotides, suggesting the preferential OS coordination between the N7 atoms and Co^2+^ ions (Fig. 6E). Since these nucleotides are present at J_34_ junction (Fig. 3A), Co^2+^ ions binding in this region facilitate their interactions with the proximal poc12 nucleotides through OS coordinations in the I state. Therefore, we predict that NRA captures two hydrated Co^2+^ ions in poc12 of state I (Fig. 6F), followed by a torsional twist at the 4WJ that takes NRA from state I to state F. Due to the twist, the bound hydrated Co^2+^ ions (Co1, Co2) lose water molecules from their first hydration shell and get trapped inside the IBD cavity via OS to IS transition.

### 3.9 Co^2+^ Binding Specificity is due to Enhanced Orbital Interactions with the Guanine Nucleotides

To further investigate the role of N7 atoms in the binding selectivity of Co^2+^ in F-NRA, we examined the orbital interactions between the metal ions and the poc12 nucleotides involved in octahedral coordination with the metal ions in the native 4WJ geometry. We performed DFT-based geometry optimization on a segment of the IBD, which includes the bound metal ions (Co1 and Co2), poc12 nucleotides (G41, G42, G82, and G83), and the water molecules involved in IS coordination with Co1 and Co2 (Fig. S27A). We refer to this complex as Co_2_G_4_. Following the same protocol, we constructed another complex by replacing Co^2+^ ions in Co_2_G_4_ with Mg^2+^ ions, and this complex is referred to as Mg_2_G_4_. We independently optimized both the complexes (Fig. S27A,B). In the optimized Co_2_G_4_ complex, we observed that the coordination environment is the same for Co1 and Co2 ions. This is also true for the Mg_2_G_4_ complex. Based on this result, we constructed and optimized Co_1_G_2_ and Mg_1_G_2_ complexes (half of Co_2_G_4_ and Mg_2_G_4_ complexes), which are also octahedral (Fig. S27C,D).

To compare the ion-dependent stability of the complexes, we formulated a hypothetical isodesmic ion-exchange reaction:

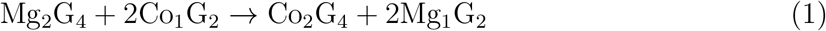

In this reaction, the negative free-energy change, Δ*G*_*iso*_ = -4.8 kcal mol^−1^ suggests that replacement of Mg^2+^ by Co^2+^ ions from Mg_2_G_4_ complex to form Co_2_G_4_ is thermodynamically feasible process (Table S4). To further investigate the origin of the higher thermodynamic stability of Co_2_G_4_ over Mg_2_G_4_ complex, we performed Bader’s atom-in-molecule (AIM) analysis for both the complexes^50,51^ and compared the potential energies, *V*_BCP_, at the metal-ligand bond-critical points (BCP) (Table S5).^64,65^ A more negative *V*_BCP_ value corresponds to a stronger interaction. We find that the *V*_BCP_ between N7 atoms and Co^2+^ is higher in magnitude compared to Mg^2+^ indicating a significantly enhanced stability of the Co_2_G_4_ coordination complex over Mg_2_G_4_ (Fig. 7A, S28A, Table S5). This finding further corroborates our conclusion from all-atom simulations that the N7 atoms of guanine in the poc12 are crucial for the selective sensing of Co^2+^ over Mg^2+^. We also observed that the bond energies between the inner-shell coordinated oxygen atoms (O2^*′*^) of the sugar moieties and Co^2+^ are also approximately twice as large compared to Mg^2+^, further highlighting the preferential interaction of Co^2+^ with the nucleotides in poc12.

**Figure 7.**
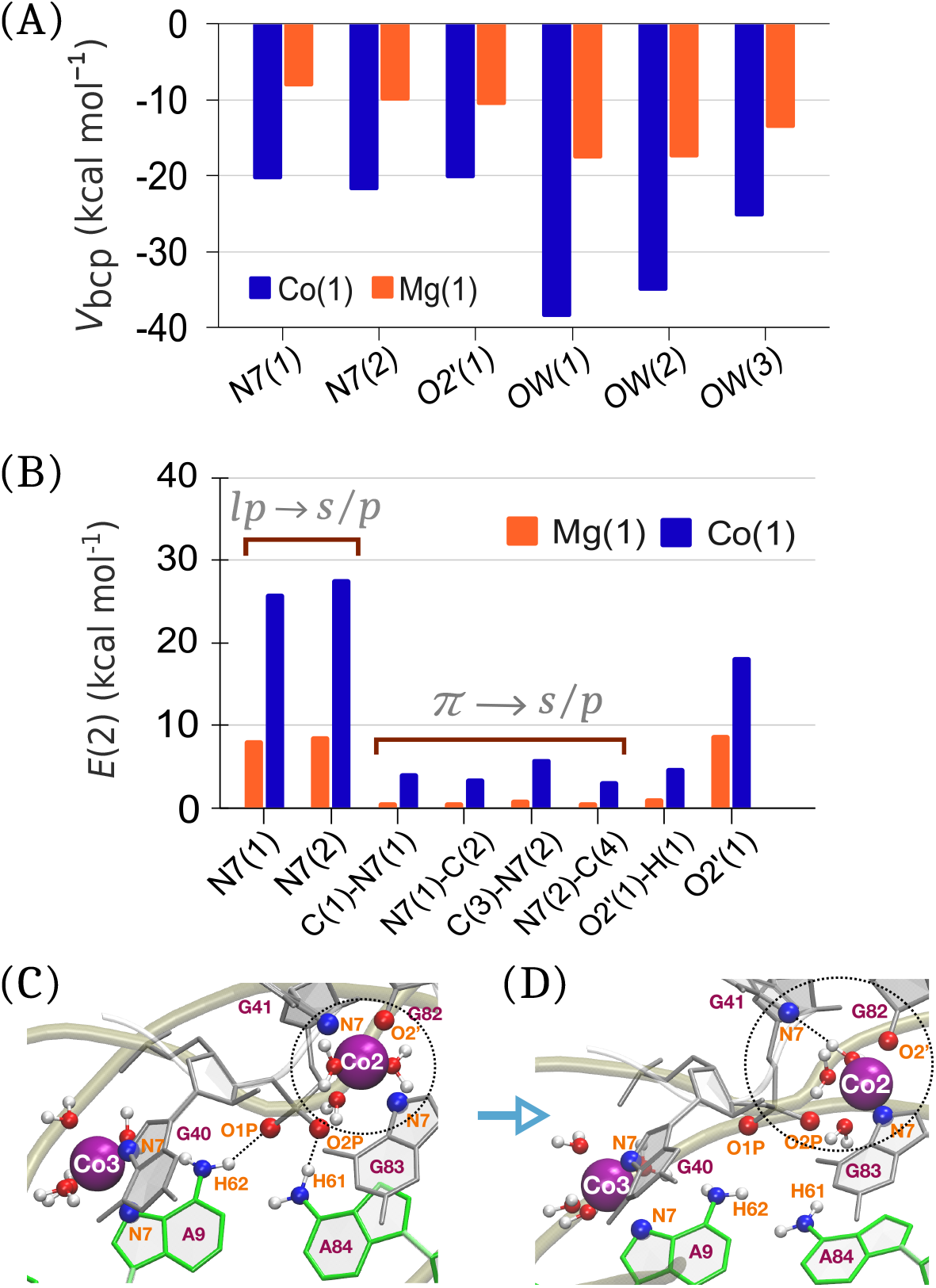
**(A)** The comparison of *V*_bcp_ (kcal mol^−1^) between Co(1) and Mg(1) in Co_2_G_4_ and Mg_2_G_4_ complexes, respectively (Table S5, Fig. S27A,B). **(B)** The comparison of cumulative interaction energies, *E*(2) (kcal mol^−1^), between NBOs for Co(1) and Mg(1) ions in Co_2_G_4_ and Mg_2_G_4_ complexes, respectively (Fig. S27A,B). Individual components are in Table S6. **(C)** and **(D)** Representative snapshots of IS to OS transition of Co2 from G41(N7) inside the binding pocket (dotted circle). This transition expanded the pocket and broke the base phosphate H-bonds (dotted line) (A9 and A84 with G41).

To further explore the orbital interactions involved in the metal-RNA coordination, we performed a natural bond orbital (NBO) analysis to calculate the second-order perturbation interaction energies (*E*(2)).^52,64^ We observed that the NBO stabilization energies, *E*(2), between the lone pairs of the N7 atoms with Co^2+^ in Co_2_G_4_ complex are 3-fold higher than that of the Mg_2_G_4_ complex. This suggests that poc12 nucleotides interact more strongly with Co^2+^ than with Mg^2+^ (Fig. 7B, S28B, S29, Table S6 and S7). Additionally, the *π* orbitals associated with the N7 nucleotides strongly interact with the empty orbitals of Co^2+^, which is not the case for Mg^2+^ (Fig. 7B, S30). These exclusive orbital interactions involving N7 atoms and Co^2+^ further support our observations from the all-atom simulations.

### 3.10 Co2 Coordination Regulates Co3 Binding Stability

The in-line probing analyses performed on G81, G82, and G83 by conditional N7 substitutions suggest that the binding of Co^2+^ ions in poc12 and poc3 occurs through the interconnecting conserved nucleotides (G40, G41, G42, G82, and G83) in a cooperative manner.^12^ To assess the correlation between Co^2+^ ions (Co1, Co2, and Co3) in poc12 and poc3, we performed additional simulations summarized in Table 1 (see Methods). We performed unbiased molecular dynamics simulations on the NRA(1,2,3) system to probe whether three Co^2+^ ions are stable in their binding pockets. We observed that Co1 and Co2 ions are stable in their binding pockets (Fig. S31). However, the Co3 ion broke two of its IS contacts with N7 (A9 and G40), forming an OS contact, and eventually left poc3. (Fig. S31, S32A,B).

To understand the influence of Co1 and Co2 on Co3 unbinding, we performed additional simulations of NRA(1,3) and NRA(2,3) (Table 1) and measured the average residence time of Co3, ⟨*τ*⟩. ⟨*τ*⟩ is the average time taken by Co3 to leave the crystal binding mode and go to the bulk. To track the position of Co3, we computed the distances between Co3 and N7 atom of A9 (*d*_Co3-A9N7_) and G40 (*d*_Co3-G40N7_). The results showed a significantly higher ⟨*τ*⟩ for NRA(2,3) (584 ± 77 ns) compared to NRA(1,3) (219 ± 81 ns), indicating that Co2 plays an important role in stabilizing Co3. The ⟨*τ*⟩ value will depend on the forcefields. ^66^ However, we believe that the overall trend will be consistent across all force fields. Structural analysis revealed that in NRA(2,3), the transition of Co2-G41(N7) coordination from IS to OS increased *d*_Co2-G41(N7)_. As a result, *d*_A9(H62)-G41(O1P)_ and *d*_A84(H61)-G41(O2P)_ increases, weakening base-phosphate hydrogen bonds (A9-G41 and A84-G41) and increasing thermal fluctuations in Co3-binding nucleotides A9 and G40 (Fig. 7C,D and S33). Additionally, the increased fluctuation in the 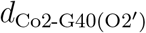, also contributes to the exit process of Co3 (Fig. S34A,B).

Subsequently, the increasing *d*_Co3-A9(N7)_ and *d*_Co3-G40(N7)_ confirms the departure of Co3 from the poc3. In NRA(1,3), the absence of Co2 led to rapid disruption of A9-G41 and A84-G41 base-phosphate hydrogen bonds, causing higher nucleotide fluctuations and allowing Co3 to exit more easily(Fig. S35). Interestingly, a K^+^ ion diffused into the pocket in the absence of Co2. We did not find any influence of Co1 ion on the Co3 unbinding events because of the absence of interconnecting nucleotides between Co1 and Co3. These findings demonstrate that Co2 coordination regulates nucleotide fluctuations inside the pockets, ultimately controlling Co3 binding stability.

## 4 Discussion

### Mechanism of NRA Folding and Co^2+^ Binding

In the unfolded state U, the helical pairs P1|P4 and P2|P3 in NRA are coaxially stacked. With an increase in divalent ion concentration, the ions bind to the IBD nucleotides from *J*_14_ and *J*_23_ and shorten all the junction loops, driving the change in the coaxial stacking of the helical pairs to P1|P2 and P3|P4 populating the intermediate state I (Fig. 8A,B). The change in coaxial stacking arrangement reduces the cavity volume in the 4WJ and creates an anionic pocket, where two hydrated Co^2+^ ions preferentially bind to the IBD nucleotides (poc12: G41, G42, G82, G83) in OS coordination (Fig. 8B). A torsional twist at the 4WJ further leads to compaction of the anionic cavity, and the NRA transitions to the folded state F (Fig. 8C). During this transition, bound hydrated Co^2+^ ions (Co1, Co2) lose water molecules and transition to IS coordination with Guanine nucleotides in the IBD (Fig. 8D). The torsional twist compacts the cavity volume, locks the bound Co^2+^ ions, and also configures the poc3 ion binding site (A9, G40). Finally, a Co^2+^ ion (Co3) binds to poc3 with the cooperative assistance of bound Co2 (Fig. 8E).

**Figure 8.**
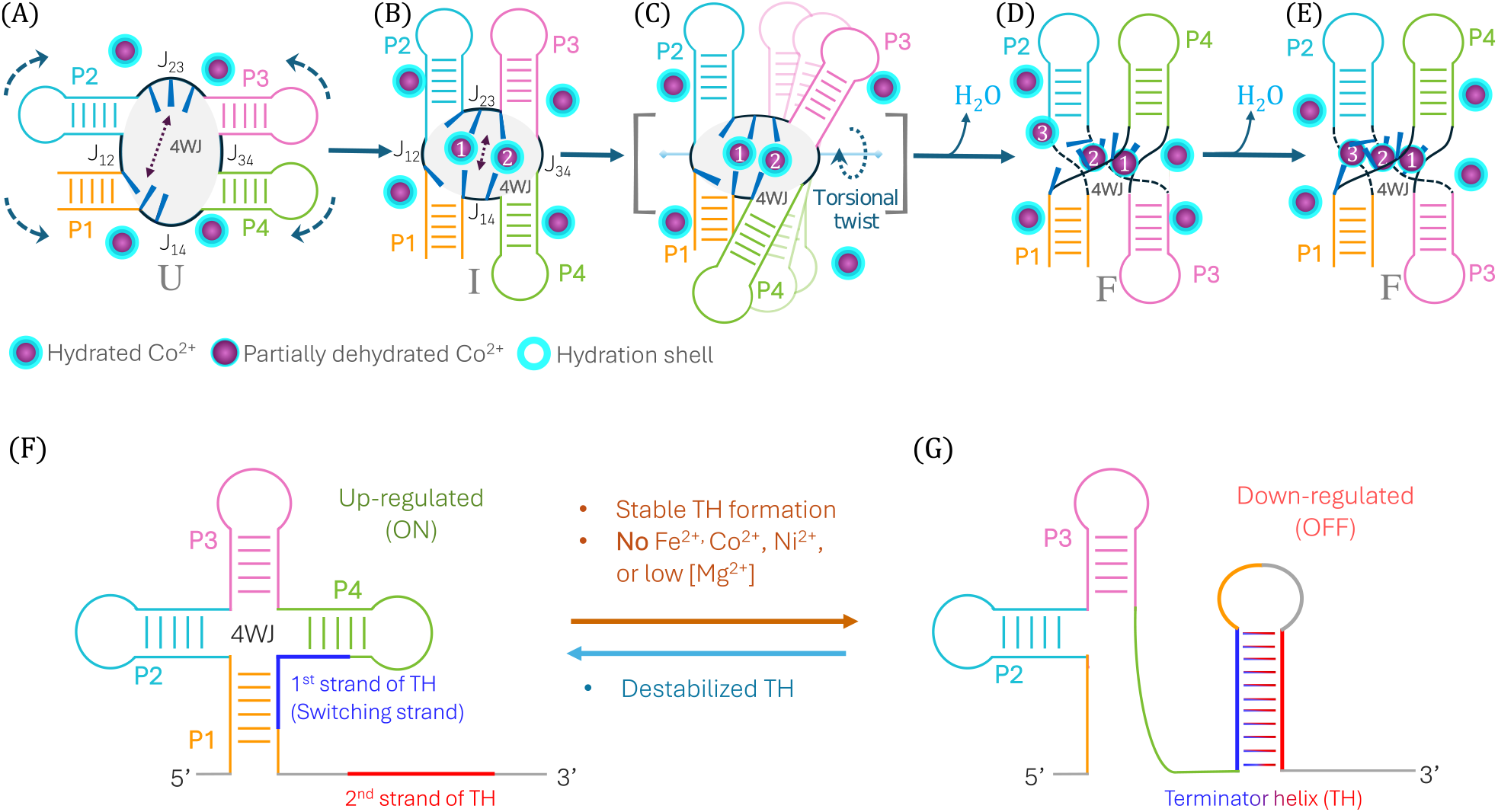
Mechanism of Co^2+^ binding to the IBD in NiCo riboswitch. The NRA domain with four helices (P1 (orange), P2 (cyan), P3 (mauve), and P4 (green) and the 4WJ (gray shade). The junctions (J_*ij*_) are shown as black lines. Six IBD nucleotides are marked with deep-blue solid sticks. The Co^2+^ ions in purple circles are shown with their hydration shell in cyan around them. **(A)** State U with a widely open IBD. **(B)** State I with relatively compact IBD and OS coordinated Co^2+^ ions. **(C)** Probable transition state where the 4WJ undergoes a torsional twist. **(D)** The coordination of Co2 in poc12 pocket helps another Co^2+^ ion (Co3) to bind at poc3 pocket in state F after eliminating two water molecules from the first solvation shell of Co3. **(E)** Folded state F with the three bound Co^2+^ ions. Schematic of a competitive regulation mechanism involving terminator helix (TH), showing **(F)** up-regulated (ON) and **(G)** down-regulated (OFF) form of full-length NiCo riboswitch.

NRA binds to three Co^2+^ ions and uses the switching strand (C74 to A74) embedded inside P1 and P4 to detect transition metal ions (Fig. 1B).^12^ This mechanism differs from other metalloregulatory riboswitches such as Mn^2+^-sensing riboswitch.^13^ The Mn^2+^ sensing riboswitch binds only to a single Mn^2+^ ion to a structure, whose folding is driven by Mg^2+^ ions, and the bound Mn^2+^ ion does not interact directly with the switching strand.

### Weak Non-Canonical Base-pairs at the Helix-junctions are Pivotal for RNA Functions

The coaxial stacking of helices is a ubiquitous tertiary interaction in RNA junctions, and it is essential to adopt functionally active structures. ^59,67^ We show that at low divalent ion concentration, the formation of native-like coaxial stacking of NRA helices is less stable due to the breakage of weak non-canonical G*·*A base pair hydrogen bonds at the helix terminals present near the 4WJ, making it highly flexible. However, at high divalent ion concentrations, the ions bound to the 4WJ help stabilize the weak G*·*A pairs, induce the native-like coaxial stacking, and reduce the flexibility of the 4WJ domain. Therefore, we infer that weak base pairs are key to tuning the divalent ion concentration-dependent native-like coaxial stacking of helices and the formation of functional RNA structures.

Weak base pairs, such as G*·*A, A*·*U, A*·*A, etc. are commonly found in various RNA structures.^59,61^ For example, in Mn^2+^-riboswitch, the 4WJ and ligand (Mn^2+^) binding domains are distant from each other (PDB ID: 6N2V^13^). The Mn^2+^-binding domain should be formed for the riboswitch to perform its function, which is possible only if the helices are in native-like coaxial stacking. The NRA simulations suggest that the stable weak base pairs at the junctions can strongly influence the stability of the native-like coaxially stacked helices. Interestingly, Mn^2+^-riboswitch also has two weak base-pairs, A*·*U and U*·*U near its 4WJ.^13^ Therefore, these weak base pairs could be disrupted in low divalent ion concentrations, leading to a flexible 4WJ domain and poor coaxial stacking of helices. As a result, the Mn^2+^-binding domain will be unavailable for Mn^2+^-binding, making the riboswitch dysfunctional. In contrast, higher divalent ion concentrations stabilize these weak base pairs at junctions, allowing the RNA to adopt the native-like coaxial stacking and function effectively. Similarly, the 16s and 23s ribosomal subunits, hairpin ribozyme, FMN riboswitch, SAM-III riboswitch, etc., also possess weak base pairs at different junctions.^18,56,59,61,68^ Thus, weak base pairs near the junctions can be critical for controlling the RNA functions.

### Terminator Helix Formation for Gene Regulation

Transcriptional riboswitches start folding as they are synthesized by RNA polymerase, and the final folded structure, which depends on whether the cognate ligand is bound to it or not, signals whether transcription is to proceed or be terminated.^69^ For the full-length NiCo riboswitch comprising the aptamer domain and expression platform, the regulatory mechanism depends on the availability of transition metal ions for binding. In the absence of Co^2+^, Ni^2+^, and Fe^2+^, the full-length NiCo RNA downregulates genes by terminating the transcription process through the formation of a terminator helix (TH).^12^ However, in the presence of cognate ions during transcription, the initially transcribed aptamer domain (NRA: A1 to U93) folds and tightly captures the switching strand (first strand of TH: C74-A84) inside the helices P1 and P4 (Fig. 1B, and S36), which prevents its pairing with the second TH strand (U95-G105) (Fig. S36) when it is transcribed eliminating the possibility of TH formation leaving the riboswitch in the ON state.

We found that in the three thermodynamic states of NRA (U, I, and F) the switching strand (C74-A84) is sequestered within P1 and P4 even without metal ion binding (Fig. 8F, 1B) suggesting that once the aptamer transcript folds, it should keep the gene ON regardless of metal binding. If the switching strand is sequestered within P1 and P4, even without metal ions, the question arises of how TH forms to repress transcription.

We propose that in the absence of transition metal ions, even if the first strand of TH is sequestered into the P1 and P4 helices in the U state (Fig. 8F), the synthesis of the second strand should initiate a competitive structural reorganization disrupting the P1 and P4 helices to form thermodynamically stable TH, which will halt transcription and down-regulate the downstream gene (Fig. 8G). However, in the presence of cognate metal ions (Co^2+^, Ni^2+^, and Fe^2+^), ion binding stabilizes the first strand of TH sequestered in the P1 and P4 helices and inhibits TH formation. Experiments showed that even if the stability of TH is slightly reduced, the NRA will remain up-regulated irrespective of the concentration of transition metal ions.^12^ This suggests that a minimal destabilization of the TH pushes the NRA population back to the ensemble of U, I, and F states (Fig. 8F-G), depending on the metal ion concentrations. This observation substantiates the proposed competitive transcription termination mechanism. We also propose NMR or SHAPE (Selective 2^*′*^-Hydroxyl Acylation analyzed by Primer Extension) experiments to probe the formation of TH using the sequence from A1 to U106 in the presence and absence of cognate metal ions. This competitive gene regulation protocol of NiCo riboswitch can be compared to riboswitches such as the fluoride-riboswitch, Mn^2+^-riboswitch, where the ligand or Mg^2+^ induced anti-terminator formation and TH formation compete with each other. ^13,70^ Therefore, we expect that a slight destabilization of TH can also keep the corresponding genes up-regulated (ON), even in the absence of the cognate ligands.

### N7 Atoms in Gaunines Present in the IBD are Vital for Selective Binding of Co^2+^, Ni^2+^, and Fe^2+^

All-atom simulations showed that in states F and I, the N7 atoms on the conserved G nucleotides in IBD exhibited a preference for Co^2+^ binding over Mg^2+^ (Fig. 6B-D). Transition metal ions are known to form complexes better than *s*-block metal ions. In the case of Co^2+^ sensing proteins, the occurrence of Histidine is highest, and it is directly involved in Co^2+^ binding via IS coordination with nitrogen atoms.^71^ In NRA, the G nucleotides have a nitrogen-rich imidazole moiety similar to Histidine, which it exploits for specific binding to the transition metal ions Co^2+^, Ni^2+^, and Fe^2+^.

Previous studies showed that Mg^2+^ ions preferably interact with phosphate groups compared to the nucleobases,^72–75^ and interaction of Mg^2+^ with base nitrogens is less common. However, transition metal ions interacted more with the base nitrogen.^75,76^ The DFT calculations we performed to probe the influence of nitrogen atoms on Co^2+^ binding showed that the NRA bound with two Co^2+^ ions (Co1 and Co2) is thermodynamically more stable than that of the Mg^2+^-NRA complex. AIM analysis supported stronger binding of Co^2+^ compared to Mg^2+^ as it has lower potential energies at the bond critical points (BCP) (Fig 7A, S28A, and Table S5). NBO analysis further suggested that in addition to the direct coordinate bonds, the *π*-orbitals of the nitrogen atoms (N7) interact significantly with the *p*-orbitals of Co^2+^ (Fig. 7B, Table S6, S7). The in-line probing analyses performed by substituting the N7 atoms in G81, G82, and G83 by Furukawa and co-workers showed diminished ion-binding affinity in NRA, which supports our observation.^12^ We performed the same electronic structure analyses to understand the affinity of Ni^2+^ and Fe^2+^ binding to NRA. The calculations show that the binding affinity followed the Irving-Williams series (Fe^2+^ < Co^2+^ < Ni^2+^) as expected^12,21,77^ (data not shown).

## 5 Conclusion

We showed that the four helices (P1 to P4) and the 4WJ present in a czcD riboswitches adopt three broad subensembles (U, I, and F states) of distinct, rapidly interconverting conformations in the presence of divalent metal ions (Mg^2+^ and Co^2+^) without altering the overall hydrogen bond network in agreement with the standard two-color smFRET experiment.^20^ The strong selectivity of czcD riboswitches towards the cognate transition metal ions over alkaline earth metal ions is cooperatively controlled by the N7 atoms in the conserved guanine nucleotides. We further provided insights into the structure of the I state, demonstrated the importance of the coaxial stacking arrangement of helices in each state, and the ion binding mechanism. The predicted U and I states can be verified by NMR experiments and advanced three-color smFRET experiments.^17^ Using electronic structure calculations, we quantified the relative binding affinity of Co^2+^ over Mg^2+^ to the IBD. We show that the imidazole moiety in the guanine nucleotides is critical for selective transition metal ion binding due to their enhanced orbital interaction. The insights from this work can be exploited to engineer tunable biosensors for real-time monitoring of transition metal ion concentrations and to develop antimicrobials by disrupting the gene regulation mechanism that helps maintain the homeostasis of toxic transition metal ions.

## Supporting information

NiCo Riboswitch Folding

## 6 Competing interests

No competing interest is declared.

## 7 Author contributions

D.M., S.H., and G.R. designed research; D.M., S.H. performed research; D.M., S.H., L.B., M.H., and G.R. contributed analytic tools; D.M. and S.H. analyzed data; and D.M., S.H., L.B., and G.R. wrote the paper.

## 8 Supporting Information

Simulation details and data analysis; Coarse-grained parameters (Table S1 and S2); DFT calculations (Table S3−S7); *f*_*T C*2_ distribution (Fig. S1); FES (Fig. S2 and S3); dihedral angle definition (Fig. S4); contact maps and PDMs (Fig. S5, S6 and S8); schematic of helixcoaxiality in states (Fig. S7); terminal base-pairs in PDB (Fig. S9); secondary structure with broken G*·*A pairs in state U (Fig. S10); free-energy change on ion-induced TC2 contact formation (Fig. S11); CN and *χ* with [Mg^2+^] (Fig. S12); ⟨*CN*⟩ and ⟨*R*_*g*_⟩ of IBD (Fig. S13 and S15); distribution *R*_*g*_ of IBD (Fig S14 and S16); local ion concentrations and electrostatic potential surface (Fig. S17−S21); simulation snapshot (Fig. S22); RDF, *g*(*r*) and CN (Fig. S23−S25); state I with bound Co^2+^ at IBD (Fig. S26); DFT optimized structures (Fig. S27); AIM and NBO analyses data (Fig. S28 and S29); N7 NBO interaction with Co^2+^ and Mg^2+^ (Fig. S30); change in distances and Snapshot of OS interactions in NRA(1,2,3) (Fig. S31 and S32); change in distances for NRA(2,3) and NRA(1,3) (Fig. S33−S35); secondary structure of the E. bacterium NRA (Fig. S36); movie of transitions from U→I→F (Movie S1); DFT Geometry Optimized Coordinates.

## 9 Abbreviations

NRA: NiCo riboswitch aptamer
PDM: Pair distance matrix
FES: Free energy surface
CG: Coarse-grained
AA: All-atom
DFT: Density functional theory
TC: Tertiary contacts
IBD: Ion binding domain
IS: Inner-shell
OS: Outer-shell
AIM: Atoms in molecule
NBO: Natural bond orbitals

## 10 Acknowledgement

We thank Prof. Garima Jindal for guidance on electronic structure calculations and data analyses. GR acknowledges funding from the Science and Engineering Research Board (SERB) through the grant CRG/2023/002817. DM and MH acknowledge the research fellowship from the Indian Institute of Science, Bangalore. SH acknowledges the Prime Minister’s Research Fellowship (PMRF). We acknowledge the National Supercomputing Mission (NSM) for providing computing resources of “Param Pravega” at IISc and “Param Brahma” at IISER Pune, supported by the Ministry of Electronics and Information Technology (MeitY) and the Department of Science and Technology (DST), Government of India.

## For Table of Contents Use Only

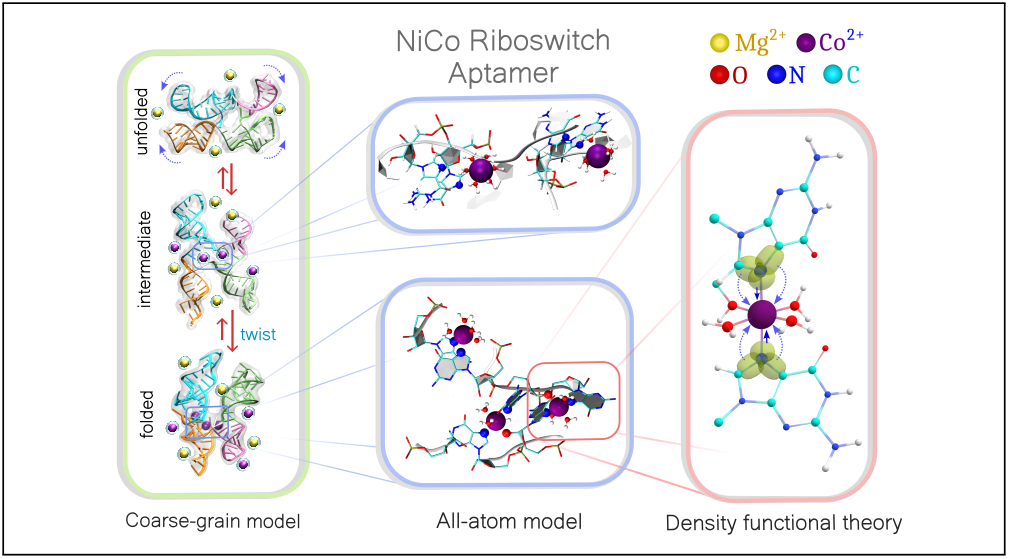

## Notes

### Competing Interest Statement

The authors have declared no competing interest.

