## Supplementary Information for "Transition Metal Binding Drives Folding of a Metalloregulatory Riboswitch by Modulating Conformational Flexibility at Helical Junctions"

§*Contributed equally to this work*

### 1 Coarse-grained (CG) Simulations

#### 1.1 Three-Interaction-Site (TIS) Model

The CG simulations were performed using the three interaction site (TIS) RNA model.[1,2] In the TIS model, a nucleotide is modeled using three beads positioned at the center of mass of the phosphate group (P), sugar (S), and nucleobase (B). The energy function in the TIS model[1] is given by

$$U_{\mathrm{TIS}} = U_{\mathrm{BL}} + U_{\mathrm{BA}} + U_{\mathrm{EV}} + U_{\mathrm{EL}} + U_{\mathrm{ST}} + U_{\mathrm{HB}} + U_{\mathrm{TST}} \tag{S1}$$

The TIS energy function consists of potentials to model bond length ($U_{\mathrm{BL}}$), bond angle ($U_{\mathrm{BA}}$), excluded volume interactions ($U_{\mathrm{EV}}$), stacking between adjacent bases ($U_{\mathrm{ST}}$), tertiary stacking (stacking between non-adjacent bases in the structure) ($U_{\mathrm{TST}}$), electrostatic interactions ($U_{\mathrm{EL}}$) and native hydrogen bond ($U_{\mathrm{HB}}$) interactions present in the folded structure.

The bond length and bond angle interactions are modeled using harmonic potentials given by,

$$U_{\mathrm{BL}} = k_\rho(\rho - \rho_0)^2 \tag{S2}$$

$$U_{\mathrm{BA}} = k_\alpha(\alpha - \alpha_0)^2 \tag{S3}$$

where $\rho_0$ and $\alpha_0$ are the bond distance and bond angles at equilibrium taken from the ideal A-form RNA helix. The $k_\rho$ values are 23, 64, and 10 in kcal.mol$^{-1}$·Å$^{-2}$ for P→S, S→P (′ →' implies 5′→3′) and S−B bonds, respectively. The $k_\alpha$ values are 5 kcal mol$^{-1}$ rad$^{-2}$ for angles involving the base beads, and 20 kcal mol$^{-1}$ rad$^{-2}$ otherwise.

The potential used for mimicking the hydrogen bonding network present in the RNA is given by,

$$U_{\mathrm{HB}} = U^0_{\mathrm{HB}} \exp(-u) \tag{S4}$$

where,

$$u = 5.0(r-r_0)^2+1.5\{(\theta_1-\theta_{1,0})^2+(\theta_2-\theta_{2,0})^2\}+0.15\{(\psi_0-\psi_{0,0})^2+(\psi_1-\psi_{1,0})^2+(\psi_2-\psi_{2,0})^2\}.$$

The value of $U^0_{\mathrm{HB}}$ is -2.9 kcal·mol$^{-1}$. The parameters $r_0$, $\theta_{1,0}$, $\theta_{2,0}$, $\psi_{0,0}$, $\psi_{1,0}$, and $\psi_{2,0}$ for all the canonical hydrogen bonds are calculated from the coarse-grained structure of an ideal A-form RNA helix and for all the non-canonical hydrogen bonds are calculated from the coarse-grained model of the riboswitch crystal structure (PDB-ID: 4RUM).[1,3] The non-canonical hydrogen bonds used in the force field are mentioned in Table S1. The definitions of $r$, $\theta_1$, $\theta_2$, $\psi_0$, $\psi_1$, and $\psi_2$ are defined in the original work of Denesyuk et al.[2] To identify the hydrogen bond network in the crystal structure of the riboswitch, we have used the WHAT IF server (https://swift.cmbi.umcn.nl),[4] RNApdbee 2.0,[5] and the Nico crystal structure (PDB-ID: 4RUM).[3]

The stacking interactions between two consecutive nucleotides are sequence-dependent ($U_{\mathrm{ST}}$) and are given by

$$U_{\mathrm{ST}} = \frac{U^0_{\mathrm{ST}}}{1 + k_r(r - r_0)^2 + k_\phi(\phi_1 - \phi_{1,0})^2 + k_r(\phi_2 - \phi_{2,0})^2} \tag{S5}$$

where, $k_r$ is 1.4 Å$^{-2}$ and $k_\phi$ is 4.0 rad$^{-2}$. The $r_0$, $\phi_{1,0}$, and $\phi_{2,0}$ are calculated from the coarse-grained structure of an ideal A-form RNA helix. The values of $U^0_{\mathrm{ST}}$ and all the definitions of $r$, $\phi_1$, and $\phi_2$ are defined in the original work of Denesyuk et al.[1]

The tertiary stacking interaction between two non-adjacent nucleobases ($U_{\mathrm{TST}}$) is given by

$$U_{\mathrm{TST}} = \frac{U^0_{\mathrm{TST}}}{1 + u} \tag{S6}$$

where,

$$u = 5.0(r-r_0)^2+1.5\{(\theta_1-\theta_{1,0})^2+(\theta_2-\theta_{2,0})^2\}+0.15\{(\psi_0-\psi_{0,0})^2+(\psi_1-\psi_{1,0})^2+(\psi_2-\psi_{2,0})^2\}.$$

The value of $U^0_{\mathrm{TST}}$ is taken to be -6.5 kcal·mol$^{-1}$. The definitions of the other variables in the potential are the same as the hydrogen bonding interaction potential used in the work of Denesyuk et al.[2]

The electrostatic interaction between the phosphate beads is computed using the Coulomb potential given by.

$$U_{\mathrm{EL}} = \frac{Z_i Z_j}{4\pi\epsilon_0\epsilon_r r_{ij}} \tag{S7}$$

The dielectric constant of water is a function of temperature ($\epsilon_r(T)$) and is given by

$$\epsilon_r(T) = 87.740 - 0.4008T + 9.398\times10^{-4}T^2 + 1.410\times10^{-6}T^3 \tag{S8}$$

The excluded volume interactions between the beads is given by a modified LJ potential,

$$U_{\mathrm{EV}} = \begin{cases} \epsilon_{ij}\left[\left(\frac{d_s}{r+d_s-d_{ij}}\right)^{12} - 2\left(\frac{d_s}{r+d_s-d_{ij}}\right)^{6} + 1\right] & \text{if } r \le d_{ij} \\ 0 & \text{if } r > 0 \end{cases} \tag{S9}$$

where $d_s$ is the diameter of the smallest ion present in the system, $d_{ij} = R_i + R_j$, $\epsilon_{ij} = \sqrt{\epsilon_i\epsilon_j}$. The values of $R_i$ and $\epsilon_i$ for phosphate (P), sugar (S), and bases (A, G, C, and U) are taken from the work of Denesyuk et al.[2] We used the same ion parameters in both all-atom and CG simulations. The values of $R_i$ and $\epsilon_i$ for the ions are given in Table S2.

#### 1.2 Simulation Details

We performed all simulations using the TIS RNA model of the NiCo riboswitch using OpenMM.[6] The equation of motion for the particles in the simulation is given by

$$m_i \ddot{\vec{r_i}} = -m_i \gamma \dot{\vec{r_i}} + \vec{F_i} + \vec{\Gamma}_i \tag{S10}$$

where, $\vec{r_i}$ and $\gamma$ are coordinates and friction coefficient of the $i$th particle, respectively. The deterministic force on particle $i$ is given by $\vec{F_i} = -\dfrac{\partial U_{\mathrm{TIS}}(\vec{r_i})}{\partial \vec{r_i}}$, and $\vec{\Gamma}_i$ is an uncorrelated random force with a white noise spectrum. The autocorrelation function of the random force in the discretized form is $\big\langle \Gamma(t)\Gamma(t+nh)\big\rangle = \frac{2\gamma m_i k_{\mathrm{B}} T}{h}\delta_{0,n}$, where n = 0, 1, ... and $\delta_{0,n}$ is Kronecker delta function. We have used the *LangevinIntegrator* module in OpenMM to integrate the equations of motion. For better conformational sampling, we used a low friction coefficient ($\gamma$ = 1 ps$^{-1}$) with an integration time step of 2 fs. The Particle Mesh Ewald (PME) algorithm was used to compute the long-range Coulomb interactions.

We used a cubic simulation box of length 200 Å. The water solvent is implicitly modeled. The RNA and ions are explicitly present. All simulations are performed at 300 K. We performed simulations with the following ion concentrations to understand the effect of monovalent and divalent ions on the folding of NiCo riboswitch: (i) [$K^+$] = 50, 100, and 150 mM (ii) [$Mg^{2+}$] = 2, 4, 6, and 8 mM with a fixed background [$K^+$] = 150 mM, (iii) [$Co^{2+}$] = 2, 4, and 8 mM with a fixed background [$K^+$] = 150 mM and [$Mg^{2+}$]= 8 mM. To ensure adequate sampling for RNA folding thermodynamics, we performed four independent simulations for each ion concentration, and each trajectory is $\approx$ 11 $\mu$s long. We discarded the initial 1 $\mu$ s of the data from each CG trajectory for analysis.

To generate the all-atom representations of the coarse-grained structures, we used the TIS2AA program,[7] and for visualization and representation of the 3D structures, we used VMD.[8]

#### 2 All-Atom (AA) MD Simulations

We performed unbiased all-atom molecular dynamics simulations of multiple NRA systems using the GROMACS program (version 2018.6).[9,10] We extracted the initial coordinates from the X-ray crystal structure of NRA (PDB ID: 4RUM[3]). The terminal phosphate group at the 5′-end is deleted. We also removed the three $Co^{2+}$ ions labeled as Co1, Co2, and Co3 bound to the IBD. This native NRA system is labeled as F-NRA. We have converted the I state obtained from the CG simulations to the AA description using the TIS2AA[7] program. We performed AA simulations of the I state (designated as I-NRA) using the same protocol as F-NRA.

We also performed simulations of three additional systems to investigate the origin of cooperative binding of $Co^{2+}$ ions to NRA. The three systems are (i) the crystal structure of NRA with three bound $Co^{2+}$ ions (Co1, Co2, and Co3) to the IBD labeled as NRA(1,2,3). (ii) Crystal structure of NRA with two bound $Co^{2+}$ ions (Co1 and Co3) to the IBD labeled as NRA(1,3). (iii) Crystal structure of NRA with two bound $Co^{2+}$ ions (Co2 and Co3) to the IBD labeled as NRA(2,3). The details of AA simulation systems are given in Table 2 in the main text.

All the systems were solvated with TIP4P-EW[11] water molecules in a cubic box of edge length 100 Å with periodic boundary conditions implemented in all directions. We used a modified AMBER ff14 force field[12] (DESRES ff) for RNA because it was shown that this force field accurately reproduces the experimental structural and thermodynamic characteristics of single-stranded RNAs, RNA duplexes, RNA hairpins, and riboswitches.[12] The parameters for monovalent and divalent ions are taken from ref.[13,14] The parameters for the divalent ions were derived from the final optimized set for $Mg^{2+}$ and $Co^{2+}$ presented in Table 8 of Li et al.[14] study. Due to the inability of the individual hydration-free energy (HFE) and ion-oxygen distance (IOD) parameter sets to accurately reproduce both experimental HFEs and IODs simultaneously, we opted for the final optimized set. This set serves as a compromise between the HFE and IOD parameters, achieving ± 2 kcal/mol accuracy of experimental relative HFE while maintaining the coordination number (CN) for most divalent metal ions.

We used the following protocols for the simulations. The initial solvated NRA structures with the ions are energy minimized using the steepest descent minimization algorithm with a maximum force cutoff of 100 kJ $mol^{-1}$ $Å^{-1}$) followed by equilibration. The systems were equilibrated in three sequential steps: (i) NVT simulation of the solvent molecules at a temperature $T = 300$ K for 10 ns with the positions of the RNA atoms and ions restrained using a harmonic potential with a force constant of 10 kJ $mol^{-1}Å^{-2}$), (ii) NVT simulation of both the solvent molecules and ions at $T = 300$ K for 10 ns, with the positions of the RNA atoms restrained and (iii) NPT simulation of the solvent molecules and ions at $T = 300$ K and pressure $P = 1$ atm for 10 ns with the RNA atom positions restrained. Following the equilibration process, we performed the production run in the NPT ensemble at a $T = 300$ K and $P = 1$ atm without any restraints on the atoms. The temperature and pressure were controlled using a velocity rescaling thermostat[15] with time constant $\tau_t = 0.1$ ps and Parrinello-Rahman barostat[16] with time constant $\tau_p = 2$ ps. Lennard-Jones interactions are truncated at 10 Å. The particle-mesh Ewald (PME)[17] method is used to calculate electrostatic interactions, with a grid size of 1.6 Å and a real space cutoff of 10 Å. LINCS algorithm[18] is employed to maintain the rigidity of all covalent bonds containing hydrogen atoms. The leapfrog integrator is used to integrate the equations of motion with a time step of 2 fs.

### 3 Density Functional Theory (DFT) Calculations and Analyses

We used electronic structure calculations to investigate the orbital interactions between conserved guanine nucleotides in the ion binding domain (IBD) and metal ions (Table 2). We performed the DFT calculations using the Gaussian 16 suite of quantum chemical programs[19] for geometry optimizations. We used the unrestricted B3LYP hybrid density functional with Grimme's dispersion correction and Becke-Johnson damping (GD3BJ)[20] to optimize all stationary points. We used the Def2-SVP basis set for geometry optimization and Def2-TZVPP for the single-point calculations,[21] and the self-consistent reaction field (SCRF) method was incorporated with the SMD model to include solvent effect.[22]

The geometry optimization was performed on a segment of the ion-binding domain (IBD), named poc12, that comprises only water molecules and the nucleotides G41, G42, G82, and G83, which are involved in inner-shell coordination with the cobalt ions Co1 and Co2. This configuration is referred to as $Co_2G_4$, where $Co_2$ indicates the presence of two $Co^{2+}$ ions, and $G_4$ denotes the four guanine (G) bases (Fig. S27A). The overall system is charge-neutral. We have optimized the complexes with various possible multiplicities (1, 3, 5, and 7) for the two $Co^{2+}$ ions, and the configuration with a multiplicity of 7 was found to be the most stable. To facilitate a comparison with $Mg^{2+}$, we substituted $Co^{2+}$ with $Mg^{2+}$ in the crystal structure (designated as $Mg_2G_4$) and performed a separate optimization (Fig. S27B). The calculation protocol was identical for both $Co^{2+}$ and $Mg^{2+}$. We assigned a multiplicity of 1 to the $Mg^{2+}$ systems. For the comparison between the two ions, we focused solely on their most stable structures.

To formulate the isodesmic reaction, we further chopped the $M_2G_4$ complex to create two $M_1G_2$ complexes, where M = Co or Mg, $M_1$ implies one ion, and $G_2$ implies the two guanine (G) bases. We performed geometry optimization using the same protocols as described earlier (Fig. S27). The overall system is charge-neutral in all cases. Considering all the possible multiplicities (2 and 4) for one $Co^{2+}$, we have optimized the structure, and the system with multiplicity 4 was the most stable system. We considered the multiplicity of $Mg^{2+}$ systems to be 1. For comparison between the two ions, we only considered the most stable structures.

We performed AIM (Atoms in molecule)[23] analysis with the Multiwfn software on the wave functions of $Co_2G_4$ and $Mg_2G_4$ generated using Gaussian16.[19] The protocol used for the single-point calculations[24] was also used to calculate wave functions. The NBO calculations were performed using Gaussian16 with the in-built NBO 3.1 package at the same level of theory used for the single-point calculations. We used Chemcraft software to visualize the NBO interactions.

### 4 Data Analysis

#### 4.1 Structural Overlap Function ($\chi$)

To quantify the folding of NRA to the native state, we computed the structural overlap function ($\chi$)[25] given by

$$\chi = 1 - \frac{1}{N_{\text{total}}} \sum_{i=1}^{N_{\text{total}}} \Theta(\delta - \mid d_i - d_i^0 \mid) \tag{S11}$$

where $d_i$ is the distances between $i^{\text{th}}$ pair of beads in an RNA conformation, and $d_i^0$ is the corresponding distance in the crystal structure. $N_{total}$ (= 35245) is the total number of bead pairs in the TIS model of NRA. Here, we only considered pairs of beads that are at least 3 nucleotides separated from each other. $\Theta$ is the Heaviside step function, and $\delta = 5$ Å. The values $\chi = 0$ and 1 correspond to the structure of a conformation that is highly similar or dissimilar to the crystal structure, respectively.

#### 4.2 Dihedral Angle of Between Helices ($\phi$)

To quantify the relative orientation of the helices, we defined the dihedral angle $\phi$ between the P2 and P4 helices (Fig. S4 and Fig.2D). To calculate $\phi$, we used the center of mass positions of the phosphate backbone beads of nucleotides C21, C22, G28, and G29 for the P2 helix head; U62, G63, C74, and A75 for the P4 helix head; A9, C11, G40, A84, and C85 for the P1-P2 junction; G42, A56, A57, G81, G82 for the P3-P4 junction (Fig. S4).

#### 4.3 Free Energy Surface (FES) from CG Simulations

We projected the free energy surface (FES) onto $\chi$ to identify the various thermodynamic states populated during NRA folding. We computed the FES $G(\chi)$ using the equation

$$G(\chi) = -k_{\mathrm{B}}T \ln\bigl(P(\chi)\bigr) \tag{S12}$$

where $k_{\mathrm{B}}$ is the Boltzmann constant, and $P(\chi)$ $\bigl(= \int \mathrm{d}\boldsymbol{r}\ \delta[\chi - \chi(\boldsymbol{r})]P(\boldsymbol{r})\bigr)$ is the probability that RNA conformations given by coordinates $\boldsymbol{r}$ have a structural overlap function value $\chi$.

#### 4.4 Computation of Pair distance matrices (PDMs) and Contact maps

To compute the average pair distance matrices (PDMs), we have calculated the pair distances between all the RNA CG beads, excluding the beads that are less than three nucleotides apart from each other in the RNA sequence. To compute the average contact map, we used the pair distances from the PDMs. If a pair distance is less than the cutoff distance $r_{cut}$ (= 15 Å),[2,26] then it is considered a contact between the given pair of beads. The averaging is performed over all the conformations in the subensemble (U, I, and F).

#### 4.5 Probability of Base Pair Hydrogen Bond Formation

We calculated the probability of a base pair (BP) hydrogen bond using the equation

$$P(\alpha_{\mathrm{BP}}) = \frac{1}{N_{\mathrm{s}}}\sum_{i=0}^{N_{\mathrm{s}}} \alpha_{\mathrm{BP}} \tag{S13}$$

where

$$\alpha_{\mathrm{BP}} = \begin{cases} 1 & \text{if } U_{\mathrm{HB}} < -k_{\mathrm{B}}T \\ 0 & \text{otherwise} \end{cases} \tag{S14}$$

Here, $s$ = F, I, or U states, and $N_s$ is the total number of conformations in state $s$. $U_{\mathrm{HB}}$ is the hydrogen bond energy of the base pair (Eq. S4). We assume that there is a hydrogen bond between a base pair if its $U_{\mathrm{HB}} < k_{\mathrm{B}}T$.

#### 4.6 Coordination Number

The coordination number between two groups − divalent ions ($Co^{2+}$ or $Mg^{2+}$) and the IBD nucleotides (A9, G40, G41, G42, G81, and G82) is calculated using the equations

$$S_{ij} = \frac{1 - (\frac{r_{ij}}{r_0})^n}{1 - (\frac{r_{ij}}{r_0})^m}, \tag{S15}$$

$$CN = \sum_{i \in A} \sum_{j \in B} S_{ij} \tag{S16}$$

Here, group A consists of all the $M^{2+}$ ions (M = Co or Mg) for both CG and AA models. Group B consists of the phosphate beads of IBD nucleotides to compute $CN^{\mathrm{IBD}}_{\mathrm{M}^{2+}}$ from the CG model. Whereas, for the AA model, to compute $CN^{\mathrm{poc12}}_{\mathrm{M}^{2+}}$, group B consists of all the atoms of poc12 nucleotides (G41, G42, G82, and G83), and to compute $CN^{\mathrm{poc3}}_{\mathrm{M}^{2+}}$ group B consists of all the atoms of poc3 nucleotides (A9 and G40). $r_0$ (= 5.0 Å) is the cutoff distance for the CG and AA models, and $r_{ij}$ is the distance between atoms $i$ and $j$ from groups $A$ and $B$, respectively. The steepness of the switching function is modulated by the $n$ and $m$ values. We chose the default values of $n = 6$ and $m = 12$ for both the models.

#### 4.7 Radius of Gyration of Ion Binding Domain

To investigate the change in cavity volume of the IBD in different states with varying [$Mg^{2+}$], we computed the average radius of gyration ($\langle R_g^{\mathrm{IBD}} \rangle$) using the equation

$$R_{\mathrm{g}}^{\mathrm{IBD}} = \left( \frac{\sum_{i=1}^{N} m_i \vec{r}_i^{\,2}}{\sum_{i=1}^{N} m_i} \right)^{1/2} \tag{S17}$$

where $m_i$ is the mass of bead $i$ and $\vec{r_i}$ is the distance of bead $i$ from the center of mass of the ion-binding domain (IBD) beads (A9, G40, G41, G42, G81, and G82). $m_i$ values are in Table S2.

##### 4.8 Fraction of Tertiary Contacts

In a conformation $i$, we computed the fraction of native tertiary contacts between a particular set of nucleotides designated as $\lambda$ (TC$\lambda$) using

$$f^{(i)}_{\mathrm{TC}\lambda} = \frac{N^{(i)}_{\mathrm{TC}\lambda}}{N^{\mathrm{cry}}_{\mathrm{TC}\lambda}} \tag{S18}$$

where $N^{\mathrm{cry}}_{\mathrm{TC}\lambda}$ and $N^{(i)}_{\mathrm{TC}\lambda}$ are the number of native contacts present between the nucleotides belonging to set $\lambda$ in the crystal structure and the $i^{th}$ conformation, respectively.[27,28]

##### 4.9 Effect of Divalent Ions on Tertiary Contact Formation

To probe whether the binding of divalent ions ($M^{2+}$, where M = Mg and Co) to the nucleotides belonging to the junction loops ($J_{23}$ and $J_{14}$) affect the tertiary contact formation between these loops leading to the folding of the NRA, we calculated the free-energy change between these two states ($\Delta G^{\mathrm{M}^{2+}}_{\mathrm{TC2}}$): (i) The tertiary contacts between the junction loops $J_{23}$ and $J_{14}$ (labeled as TC2) are formed, and at least one divalent ion $M^{2+}$ is bound to both the junctions. We designate this state as $\mathrm{TC2}^{(\mathrm{For})}$. (ii) The tertiary contacts between $J_{23}$ and $J_{14}$ are ruptured and $M^{2+}$ is not bound to these junctions. We designate this ruptured state as $\mathrm{TC2}^{(\mathrm{Rup})}$.

To check whether TC2 contacts are formed or ruptured in a conformation, we initially computed the probability distribution of the fraction of native contacts in TC2 ($f_{\mathrm{TC2}}$), $P(f_{\mathrm{TC2}})$, and it is bimodal (Fig. S1). We used the condition that if in a conformation $f_{\mathrm{TC2}} > 0.5$ and $M^{2+}$ is bound we designate that it belongs to state $\mathrm{TC2}^{(\mathrm{For})}$, and if $f_{\mathrm{TC2}} < 0.5$ and $M^{2+}$ is not bound, then the conformation belongs to state $\mathrm{TC2}^{(\mathrm{Rup})}$. The free energy difference $\Delta G^{\mathrm{M}^{2+}}_{\mathrm{TC2}}$ is given by

$$\Delta G^{\mathrm{M}^{2+}}_{\mathrm{TC2}} = -k_{\mathrm{B}}T \ln\left(\frac{P(\mathrm{TC2}^{(\mathrm{For})})}{P(\mathrm{TC2}^{(\mathrm{Rup})})}\right) \tag{S19}$$

where $P(\mathrm{TC2}^{(\mathrm{For})})$ and $P(\mathrm{TC2}^{(\mathrm{Rup})})$ are the probability of observing states $\mathrm{TC2}^{(\mathrm{For})}$ and $\mathrm{TC2}^{(\mathrm{Rup})}$, respectively.

#### 4.10 Local Ion Concentration Around RNA

For both AA and CG simulations, we computed the local ion concentration ($c_j$) to investigate the condensation of positively charged divalent metal ions around RNA atoms ((a) phosphate beads for CG simulation, (b) phosphate oxygens (O1P and O2P) for AA simulation, and (c) base nitrogens (N7) for AA simulation) using the equation[2]

$$c_j = \frac{1}{N_{\mathrm{A}}V_c}\int_0^{r_c} \mathrm{d}r\ 4\pi r^2\ \rho_j(r) \tag{S20}$$

where, $\rho_j(r)$ is the number density of a specific type of ion $j$ at a distance $r$ from the specific RNA atom, $V_c$ is the spherical volume with a radius $r_c$, and $N_{\mathrm{A}}$ is the Avogadro number.

For the AA simulations, the cutoff radius is defined as $r_c^{(\mathrm{AA})} = R_{\mathrm{M}^{2+}} + R_{\mathrm{O/N}} + R_{\mathrm{IS}}$, where $R_{\mathrm{M}^{2+}}$ is the radius of the divalent metal ion ($R_{\mathrm{Mg}^{2+}} = 1.353$ Å and $R_{\mathrm{Co}^{2+}} = 1.288$ Å), $R_{\mathrm{O/N}}$ ($= 1.7$ Å) is the radius of the RNA oxygen/nitrogen atom, and $R_{\mathrm{IS}}$ ($\approx 2.0$ Å) is the inner-shell distance between metal ion and RNA atoms, which leads to $r_c^{(\mathrm{AA})} \approx 5$ Å. For the CG simulations the cutoff radius is defined as $r_c^{(\mathrm{CG})} = R_{\mathrm{M}^{2+}} + R_{\mathrm{P}} + \Delta r$ ($\approx 5$ Å), where $R_{\mathrm{P}}$ ($= 2.1$ Å) is the CG radius of a phosphate bead[2] and $\Delta r$ ($= 1.6$ Å) is the margin distance, which also leads to $r_c^{(\mathrm{CG})} \approx 5$ Å. Therefore, we set the cutoff radius $r_c = 5$ Å for both AA and CG simulations.

#### 4.11 Spatial Distribution of Ions Around RNA

We used the volmap plugin in the VMD software[29] to generate spatial density maps that illustrate the distribution of metal ions ($Co^{2+}$ and $Mg^{2+}$) around RNA nucleotides. Initially, we aligned the RNA beads from all conformations from the CG trajectory for a state to one of the representative conformations. Consequently, the positions of the metal ions in each simulation frame were adjusted accordingly. The alignment process utilized the Kabsch method[30] implemented in VMD. In our analysis, metal ions were modeled as spheres with their respective van der Waals radii. A grid with a size of 10 $Å^3$ was constructed to partition the space surrounding the RNA nucleotides. A grid point was designated as occupied and assigned a value of 1 if the sphere corresponding to a specific metal ion overlapped with it; otherwise, it was assigned a value of 0. The values at each grid point were averaged across all frames of the trajectory. VMD was also utilized for structural superposition[31] and for rendering images.[32]

#### 4.12 Average Residence Time ($\langle\tau\rangle$)

We calculated the average residence time of Co3, $\langle\tau\rangle$, by monitoring two distances − (i) distance between the Co3 ion and the N7 atom of A9 ($d_{\text{Co3-A9N7}}$) and (ii) distance between the Co3 ion and the N7 atom of G40 ($d_{\text{Co3-G40N7}}$) (Fig. 1C, S35, S33). We define the residence time, $\tau$, as the time Co3 takes to go to the bulk ($d_{\text{Co3-A9N7}} > 5.0$ Å and $d_{\text{Co3-G40N7}}$ $> 5.0$ Å) from the crystal binding modes. We performed independent simulations, each for NRA(1,3) and NRA(2,3) to calculate the $\langle\tau\rangle$.

#### 4.13 Isodesmic Reaction Formulation

To compare the thermodynamic stability of $Co^{2+}$ and $Mg^{2+}$ in the bound state, we performed geometry optimization and single point energy calculations for the following complexes, $M_2G_4$ (Fig. S27 A, B) and $M_1G_2$ (Fig. S27 C, D), where M = $Co^{2+}$ and $Mg^{2+}$. To compare the ion-dependent stability of the complexes, we formulated a hypothetical isodesmic ion-exchange reaction:

$$Mg_2G_4 + 2Co_1G_2 \rightarrow Co_2G_4 + 2Mg_1G_2; \quad \Delta G_{iso} = -4.8 \text{ kcal mol}^{-1} \tag{S21}$$

The net reaction shows the replacement of two $Mg^{2+}$ in $Mg_2G_4$ complex (Fig. S27 B) by two $Co^{2+}$ ions from two $Co_1G_2$ complexes (Fig. S27 C) to form one $Co_2G_4$ (Fig. S27 A) and two $Mg_1G_2$ (Fig. S27 D) complexes. We have computed the free energies for individual complexes using Gaussian16[19] and calculated the net free energy change ($\Delta G_{iso}$) for this reaction to contrast the thermodynamic stability of the same IBD segment bound to two different ions (Table. S4).

Table S1: Base−Base non-Watson-Crick H-bond and Base−Base tertiary Stacking Network. The number of H-bonds for a base pair is given in parentheses. All the Watson-Crick and non-Watson-Crick base pairs are shown in the secondary structure of NRA (Fig. 1B).

| Base−Base H-Bond (no. of H-Bonds) | Base−Base Tertiary Stack |
|---|---|
| G34−G37 (2), A12−G40 (2), A56−G81 (2), G34−C15 (1), A9−A84 (1), A23−G26 (1), G47−A50 (1) | G8−A86, G10−A36, G19−A32, A16−G37, G34−A36, G41−G82, G45−G53, A59−G79, G63−A75, A9−G40, A12−G83, G42−A56, A67−A69 |

Table S2: RNA[2] and ions[13,14] parameters used in the CG simulations.

| Type | $R_i$ (Å) | $m_i$ (amu) | $\epsilon_i$ (kcal mol$^{-1}$) | $Z_i$ |
|---|---|---|---|---|
| P | 2.1 | 62.971 | 0.2 | -1 |
| S | 2.9 | 131.108 | 0.2 | 0 |
| A | 2.8 | 134.119 | 0.2 | 0 |
| G | 3.0 | 150.118 | 0.2 | 0 |
| C | 2.7 | 110.094 | 0.2 | 0 |
| U | 2.7 | 111.079 | 0.2 | 0 |
| $Mg^{2+}$ | 1.353 | 24.305 | 0.009 | +2 |
| $Co^{2+}$ | 1.288 | 58.933 | 0.004 | +2 |
| $K^+$ | 1.590 | 39.098 | 0.279 | +1 |
| $Cl^-$ | 2.760 | 35.453 | 0.012 | -1 |

Table S3: Total electronic energies[a] at the stationary points.

| **System** | **Electronic energy (a.u)** | **Multiplicity** |
|---|---|---|
| $Co_2G_4$ | -9498.53414830 | 7 |
| | -9498.52416500 | 5 |

**Table S3 continued from previous page**

| System | Electronic Energy (a. u) | Multiplicity |
|---|---|---|
| | -9498.50102120 | 3 |
| | -9498.50096540 | 1 |
| $Mg_2G_4$ | -7133.25650250 | 1 |
| $Co_1G_2$ | -4805.8447454 | 4 |
| | -4805.8342735 | 2 |
| $Mg_1G_2$ | -3623.2111047 | 1 |

[a]$SMD_{(Water)}$/UB3LYP-D3(BJ)/Def2-TZVPP//UB3LYP-D3(BJ)/Def2-SVP level of theory is used.

Table S4: Single-point electronic energies and free energies for each geometry-optimized component of the isodesmic reaction. The free energies are calculated by adding thermal Gibbs free energy corrections to the electronic energies[a].

| Optimized Structure | Electronic energy (a.u) | Free energy (a.u) |
|---|---|---|
| $Co_2G_4$ | -9498.53414830 | -9497.45380930 |
| $Mg_2G_4$ | -7133.25650250 | -7132.17187350 |
| $Co_1G_2$ | -4805.8447454 | -4805.28709240 |
| $Mg_1G_2$ | -3623.2111047 | -3622.64994270 |

[a]SMD(Water)/UB3LYP-D3(BJ)/Def2-TZVPP//UB3LYP-D3(BJ)/Def2-SVP level of theory is used.

Table S5: Bond energies $E_{\mathrm{BCP}}$ from the proportionality to $V_{\mathrm{BCP}}$ at the bond-critical points obtained from Bader's atom in molecule (AIM) analysis.[23,33]

| Bond Type (Atom index) | $\rho$ | $\nabla^2\rho$ | $V_{\mathrm{BCP}}$ (kcal mol$^{-1}$) | Bond Type (Atom index) | $\rho$ | $\nabla^2\rho$ | $V_{\mathrm{BCP}}$ (kcal mol$^{-1}$) |
|---|---|---|---|---|---|---|---|
| Mg(1) - N7(1) | 0.025 | 0.138 | -8.158 | Co(1) - N7(1) | 0.051 | 0.199 | -20.421 |
| Mg(1) - N7(2) | 0.030 | 0.169 | -10.072 | Co(1) - N7(2) | 0.054 | 0.210 | -21.892 |
| Mg(1) - O2′(1) | 0.029 | 0.196 | -10.683 | Co(1) - O2′(1) | 0.045 | 0.219 | -20.274 |
| Mg(1) - OW(1) | 0.041 | 0.298 | -17.722 | Co(1) - OW(1) | 0.074 | 0.370 | -38.535 |
| Mg(1) - OW(2) | 0.035 | 0.237 | -17.640 | Co(1) - OW(2) | 0.070 | 0.338 | -35.118 |
| Mg(1) - OW(3) | 0.042 | 0.295 | -13.717 | Co(1) - OW(3) | 0.055 | 0.259 | -25.285 |
| Mg(2) - N7(3) | 0.025 | 0.138 | -8.175 | Co(2) - N7(3) | 0.050 | 0.199 | -20.404 |
| Mg(2) - N7(4) | 0.030 | 0.169 | -10.071 | Co(2) - N7(4) | 0.054 | 0.210 | -21.888 |
| Mg(2) - O2′(2) | 0.029 | 0.196 | -10.688 | Co(2) - O2′(2) | 0.045 | 0.219 | -20.288 |
| Mg(2) - OW(4) | 0.041 | 0.298 | -17.701 | Co(2) - OW(4) | 0.074 | 0.370 | -38.549 |
| Mg(2) - OW(5) | 0.035 | 0.237 | -17.625 | Co(2) - OW(5) | 0.070 | 0.338 | -35.144 |
| Mg(2) - OW(6) | 0.042 | 0.295 | -13.714 | Co(2) - OW(6) | 0.055 | 0.259 | -25.280 |

Table S6: Stabilization energies corresponding to specific NBO interactions involving Mg(1) and Co(1) in $Mg_2G_4$ and $Co_2G_4$ complexes (Fig S27), respectively, in terms of second-order-perturbation energies, $E(2)$.[34]

| **Bond Type (Atom index)** | **Mg(1)** | | **Co(1)** | |
|---|---|---|---|---|
| | **orbitals** | $\boldsymbol{E(2)}$ **(kcal mol$^{-1}$)** | **orbitals** | $\boldsymbol{E(2)}$ **(kcal mol$^{-1}$)** |
| N7(1) to M(1) | lp to s<br>lp to s | 6.78<br>1.38 | lp to p<br>lp to s<br>lp to s | 15.82<br>8.98<br>1.15 |
| N7(2) to M(1) | lp to s | 8.69 | lp to p<br>lp to s | 16.90<br>10.73 |
| C(1)-N7(1) to M(1) | $\pi$ to s | 0.64 | $\pi$ to p | 4.13 |
| N7(1)-C(2) to M(1) | $\pi$ to s | 0.65 | $\pi$ to p | 3.55 |
| C(3)-N7(2) to M(1) | $\pi$ to s | 0.90 | $\pi$ to p<br>$\pi$ to s | 4.91<br>1.02 |
| N7(2)-C(4) to M(1) | $\pi$ to s | 0.71 | $\pi$ to p | 3.19 |
| O2'(1)-H(1) to M(1) | $\sigma$ to s | 1.14 | $\sigma$ to p | 4.75 |
| O2'(1) to M(1) | lp to s<br>lp to s | 3.7<br>5.13 | lp to p<br>lp to p<br>lp to p | 14.37<br>2.36<br>1.50 |
| OW(1)-H(2) to M(1) | $\sigma$ to s | 1.26 | $\sigma$ to p<br>$\sigma$ to s | 8.04<br>2.05 |
| OW(1)-H(3) to M(1) | $\sigma$ to s | 1.29 | $\sigma$ to p<br>$\sigma$ to s | 7.21<br>1.66 |
| OW(1) to M(1) | lp to s | 9.87 | lp to p<br>lp to s<br>lp to p | 1.20<br>12.34<br>18.98 |

**Table S6 continued from previous page**

| **Bond Type (Atom index)** | **Mg(1)** | | **Co(1)** | |
|---|---|---|---|---|
| | **orbitals** | ***E*(2) (kcal mol$^{-1}$)** | **orbitals** | ***E*(2) (kcal mol$^{-1}$)** |
| OW(2)-H(4) to M(1) | $\sigma$ to s | 1.46 | $\sigma$ to p | 7.93 |
| | | | $\sigma$ to s | 1.75 |
| OW(2)-H(5) to M(1) | $\sigma$ to s | 1.25 | $\sigma$ to p | 7.30 |
| | | | $\sigma$ to s | 1.57 |
| OW(2) to M(1) | lp to s | 11.09 | lp to p | 15.25 |
| | | | lp to s | 11.06 |
| | | | lp to p | 1.38 |
| OW(3)-H(6) to M(1) | $\sigma$ to s | 1.19 | $\sigma$ to p | 6.52 |
| | | | $\sigma$ to s | 1.77 |
| OW(3)-H(7) to M(1) | $\sigma$ to s | 1.18 | $\sigma$ to p | 5.45 |
| | | | $\sigma$ to s | 1.47 |
| OW(3) to M(1) | lp to s | 9.42 | lp to p | 16.07 |
| | | | lp to s | 10.5 |
| | | | lp to p | 1.70 |

Table S7: Stabilization energies corresponding to specific NBO interactions involving Mg(2) and Co(2) for the $Mg_2G_4$ and $Co_2G_4$ complexes (Fig.S27), respectively, in terms of second-order-perturbation energies, $E(2)$.[34]

| **Bond Type (Atom index)** | **Mg(2)** | | **Co(2)** | |
|---|---|---|---|---|
| | **orbitals** | $\boldsymbol{E(2)}$ **(kcal mol$^{-1}$)** | **orbitals** | $\boldsymbol{E(2)}$ **(kcal mol$^{-1}$)** |
| N7(3) to M(2) | lp to s<br>lp to s | 6.79<br>1.38 | lp to p<br>lp to s<br>lp to s | 15.81<br>8.97<br>1.15 |
| N7(4) to M(2) | lp to s | 8.69 | lp to p<br>lp to s | 16.90<br>10.73 |
| C(5)-N7(3) to M(2) | $\pi$ to s | 0.64 | $\pi$ to p | 4.13 |
| N7(3)-C(6) to M(2) | $\pi$ to s | 0.65 | $\pi$ to p | 3.55 |
| C(7)-N7(4) to M(2) | $\pi$ to s | 0.90 | $\pi$ to p<br>$\pi$ to s | 4.91<br>1.01 |
| N7(4)-C(8) to M(2) | $\pi$ to s | 0.71 | $\pi$ to p | 3.19 |
| O2'(1)-H(8) to M(2) | $\sigma$ to s | 1.14 | $\sigma$ to p | 4.78 |
| O2'(2) to M(2) | lp to s<br>lp to s | 3.7<br>5.13 | lp to p<br>lp to p<br>lp to p | 14.43<br>2.35<br>1.45 |
| OW(4)-H(9) to M(2) | $\sigma$ to s | 1.5 | $\sigma$ to p<br>$\sigma$ to s | 8.07<br>2.05 |
| OW(4)-H(10) to M(2) | $\sigma$ to s | 1.27 | $\sigma$ to p<br>$\sigma$ to s | 7.23<br>1.66 |
| OW(4) to M(2) | lp to s | 9.07 | lp to p<br>lp to s<br>lp to p | 19.03<br>12.34<br>1.21 |

**Table S7 continued from previous page**

| **Bond Type (Atom index)** | **Mg(2)** | | **Co(2)** | |
|---|---|---|---|---|
| | **orbitals** | $\boldsymbol{E(2)}$ **(kcal mol$^{-1}$)** | **orbitals** | $\boldsymbol{E(2)}$ **(kcal mol$^{-1}$)** |
| OW(5)-H(11) to M(2) | $\sigma$ to s | 1.6 | $\sigma$ to p<br>$\sigma$ to s | 7.94<br>1.76 |
| OW(5)-H(12) to M(2) | $\sigma$ to s | 1.42 | $\sigma$ to p<br>$\sigma$ to s | 7.32<br>1.57 |
| OW(5) to M(2) | lp to s | 10.77 | lp to p<br>lp to s<br>lp to p | 15.31<br>11.07<br>1.36 |
| OW(6)-H(13) to M(2) | $\sigma$ to s | 1.24 | $\sigma$ to p<br>$\sigma$ to s | 5.46<br>1.47 |
| OW(6)-H(14) to M(2) | $\sigma$ to s | 1.31 | $\sigma$ to p<br>$\sigma$ to s | 6.53<br>1.77 |
| OW(6) to M(2) | lp to s | 9.24 | lp to p<br>lp to s<br>lp to p | 16.12<br>10.50<br>1.70 |

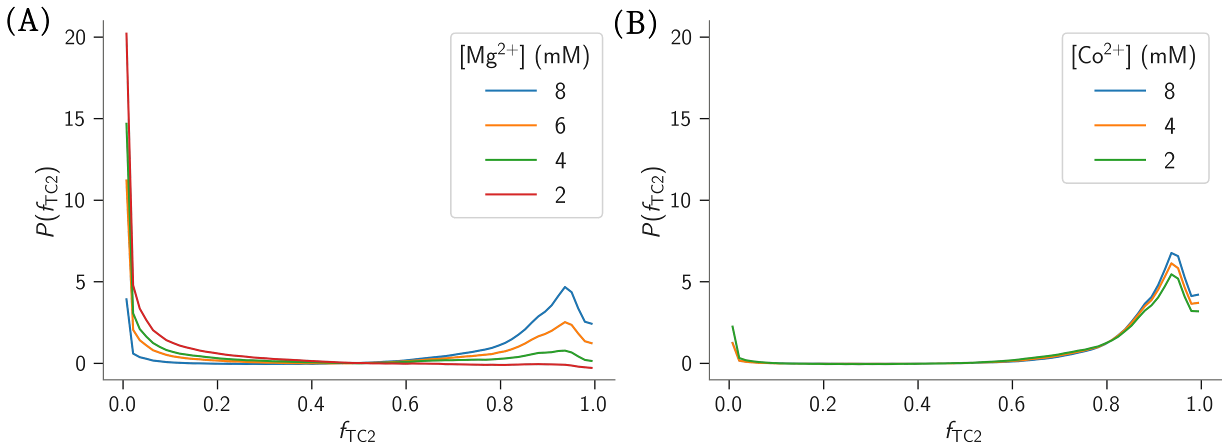


Figure S1: Probability distribution of fraction of TC2 tertiary contacts, $P(f_{TC2})$, for different (A) [Mg$^{2+}$] at a fixed background of [K$^{+}$] = 150 mM. and (B) [Co$^{2+}$] at a fixed background of [K$^{+}$] = 150 mM and [Mg$^{+}$] = 8 mM. TC2 contacts are present between two groups – group1: G40, G41, G42 and group2: G81, G82, G83

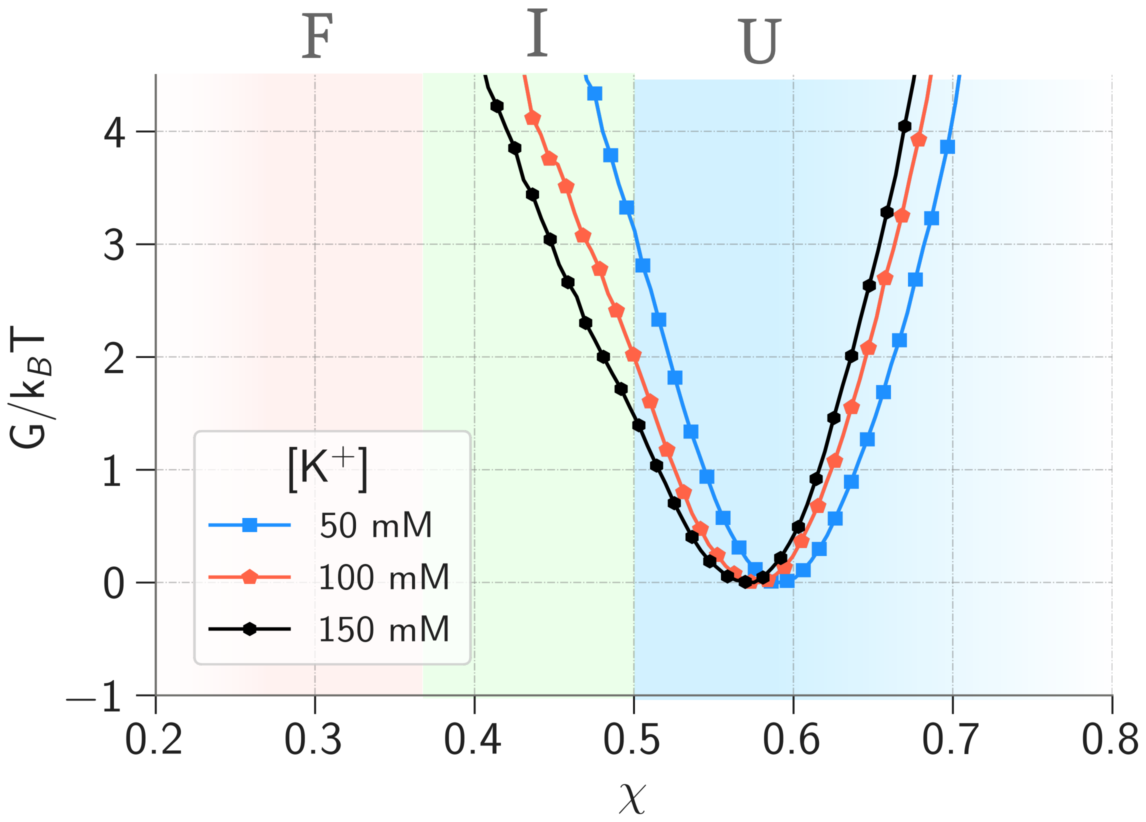


Figure S2: Free energy surface projected onto the structural overlap factor ($\chi$) for different [$K^+$] at 300 K. States F, I, and U are highlighted by shaded regions in pink, lime green, and sky blue, respectively.

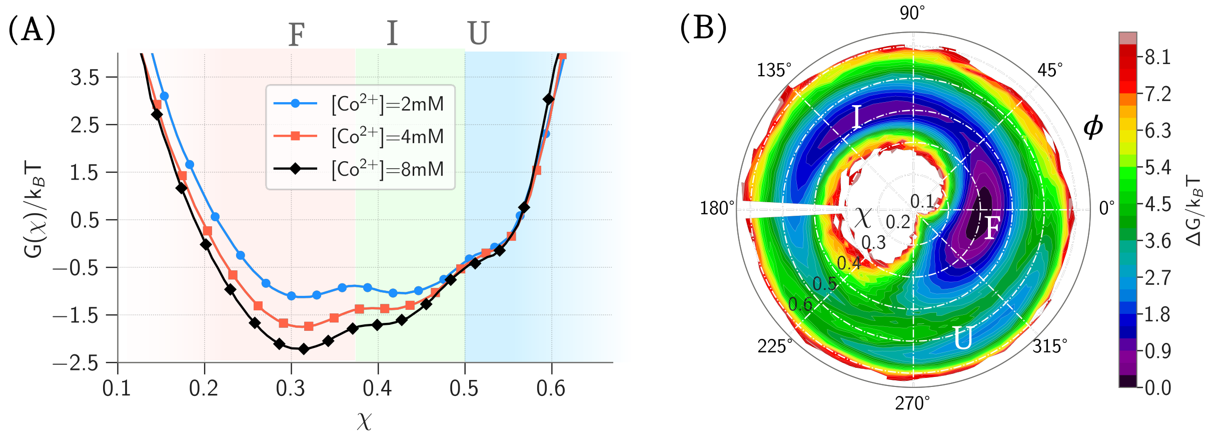


Figure S3: **(A)** Free energy surface projected onto the structural overlap factor ($\chi$) for different [$Co^{2+}$] at 300 K (with a fixed background of [$Mg^{2+}$] = 8 mM and [$K^{+}$] = 150 mM). States F, I, and U are highlighted by shaded regions in pink, lime green, and sky blue, respectively. **(B)** 2D polar FES projected onto $\chi$ and $\phi$, where $\chi$ is along the radius and $\phi$ is along the azimuth. The FES is shown for $T$ = 300 K, [$K^{+}$] = 150 mM, [$Mg^{2+}$] = 8 mM and [$Co^{2+}$] = 8 mM.

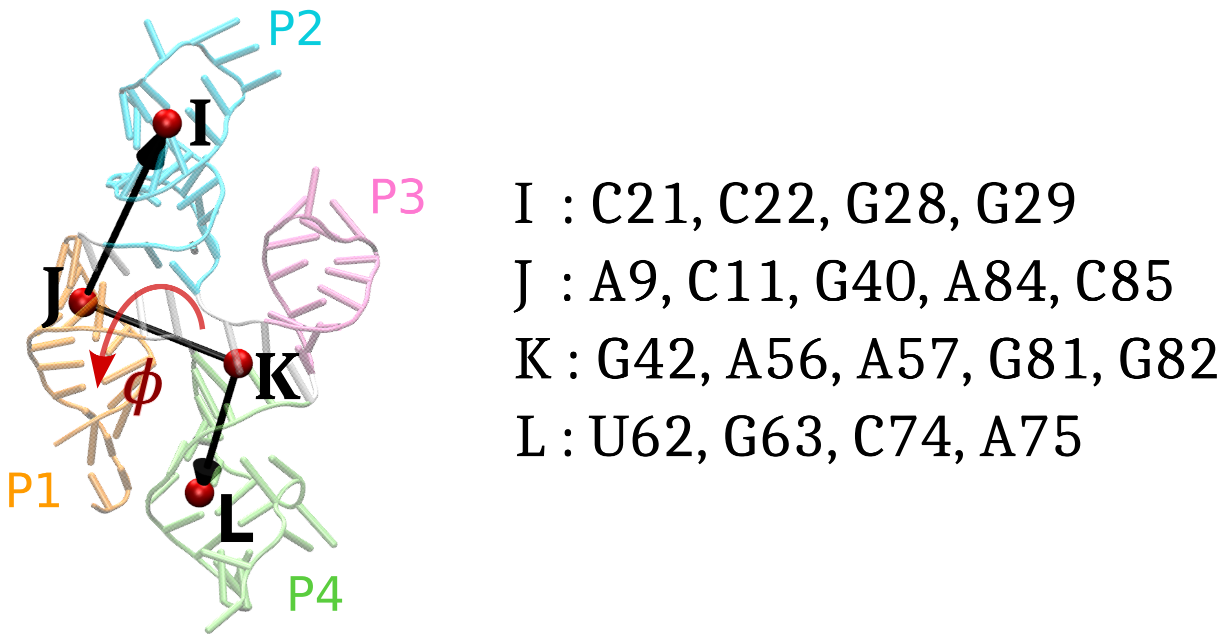


Figure S4: Dihedral angle ($\phi$) between P2 and P4 helix. $\phi$ is calculated using the center of mass (COM) of backbone beads in the sets of nucleotides I, J, K, L shown on the right panel.

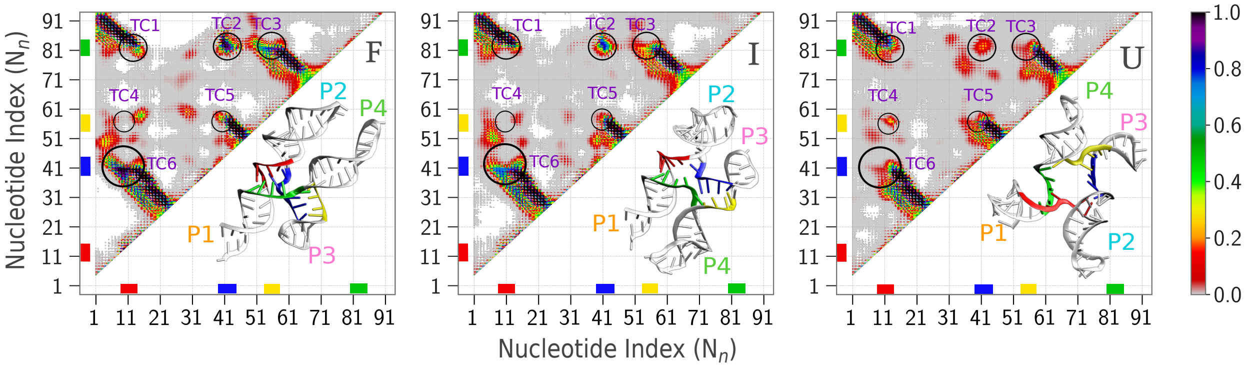


Figure S5: Contact map for three subensembles (F, I, and U states) with the representative structures. The 4WJ nucleotides on the representative structures are shown: A9-G14 (red), C39-C43 (blue), G54-A57 (yellow), and G81-A84 (green). The same color blocks on the $x$ and $y$ axes are a guide to the eye. The tertiary contacts in the 4WJ (TC1 to TC6) are highlighted using black circles.

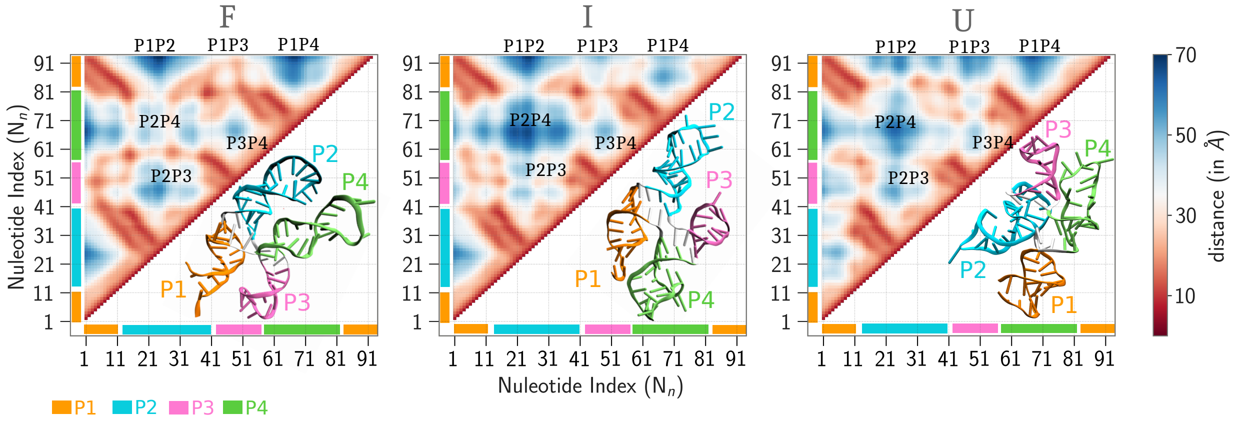


Figure S6: Pair-distance matrices for states F, I, and U. The color bars on the axes are given as a guide to the eye for identifying the helices. For the comparison of distances between the helices in different states, we used the distances between the helix head-loops for P2 (A25), P3 (A48), and P4(C68), 5′-end of NRA for P1(G1).

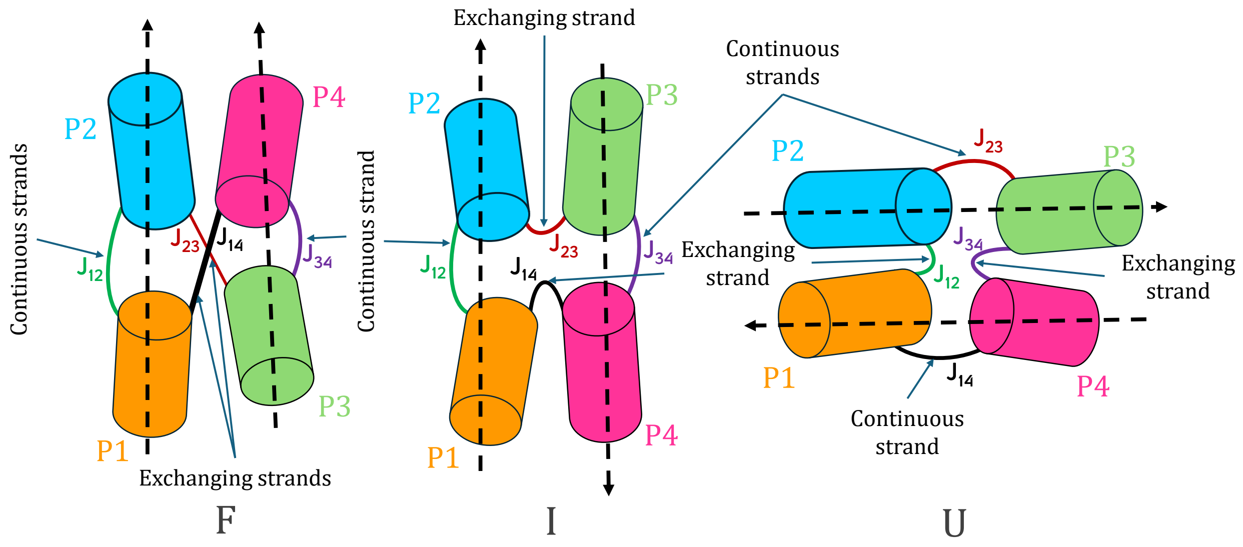


Figure S7: Schematic for states F, I, and U. All four junctions – $J_{12}$, $J_{23}$, $J_{34}$, and $J_{14}$ are shown in green, red, violet, and black solid lines. The axes of the coaxial helices are shown in black dashed arrows. The arrow direction is assigned depending on the direction of the continuous strands ($5' \rightarrow 3'$).

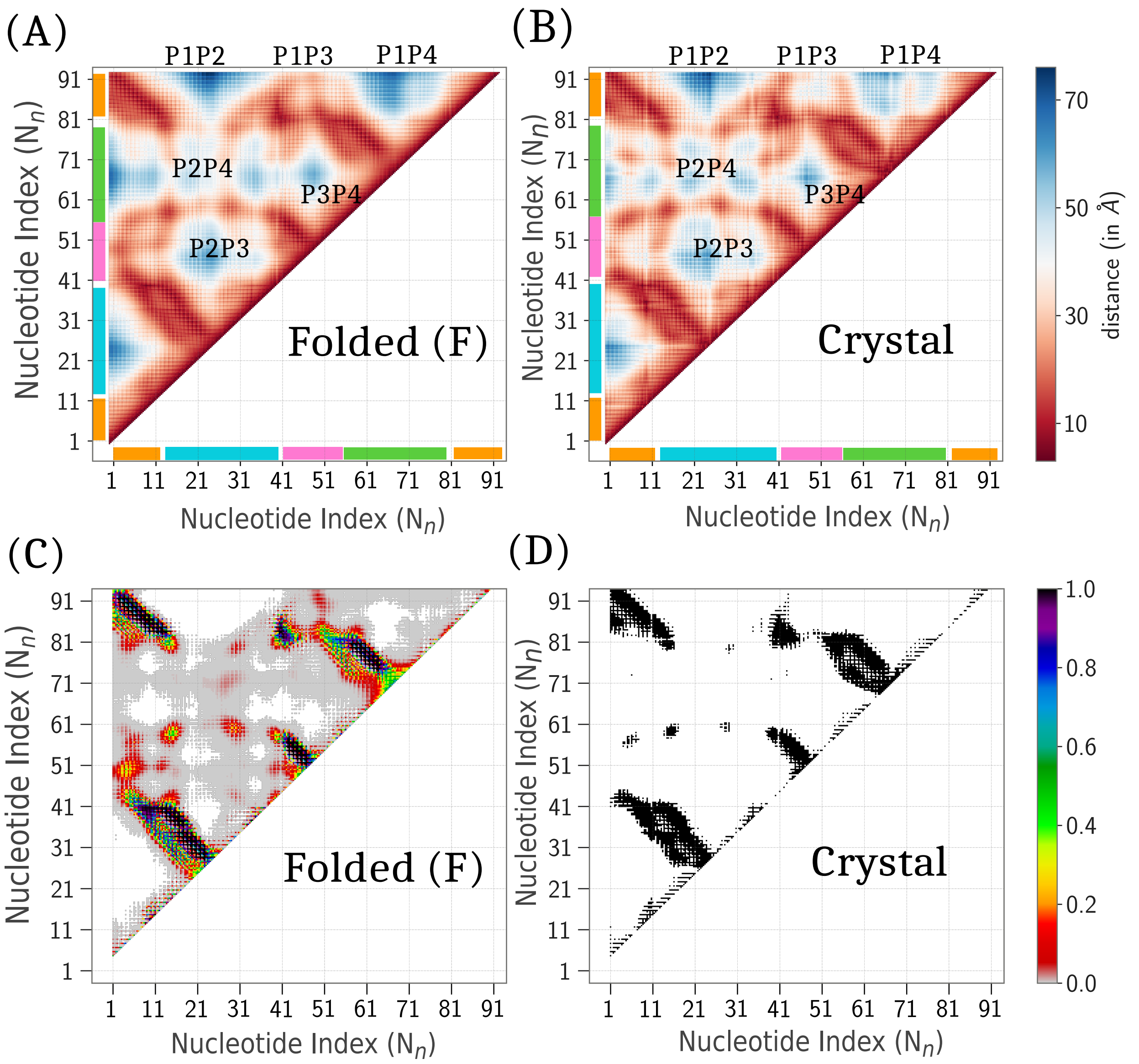


Figure S8: Pair distance matrices for (A) F state (B) crystal structure (PDB ID: 4RUM). The color bars on the axes are a guide to the eye for identifying the helices. Contact maps for (C) F state and (D) crystal structure (PDB ID: 4RUM).

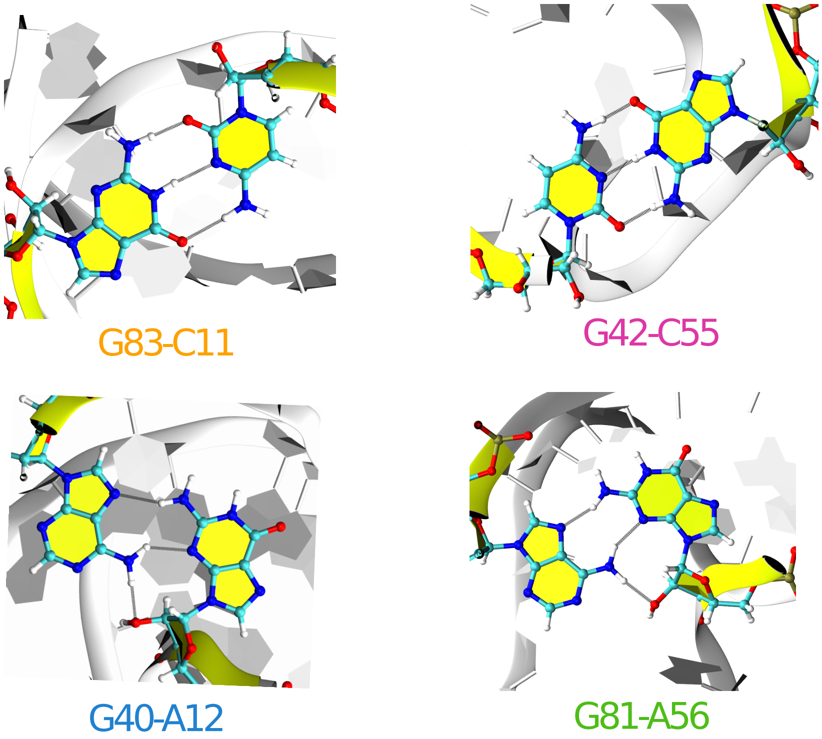


Figure S9: Terminal hydrogen-bonded base pairs from the crystal structure (PDB: 4RUM[3]). The terminal base pairs G83·C11, G42·C55, G40·A12, and G81·A56 are from helices P1, P3, P2 and P4, respectively.

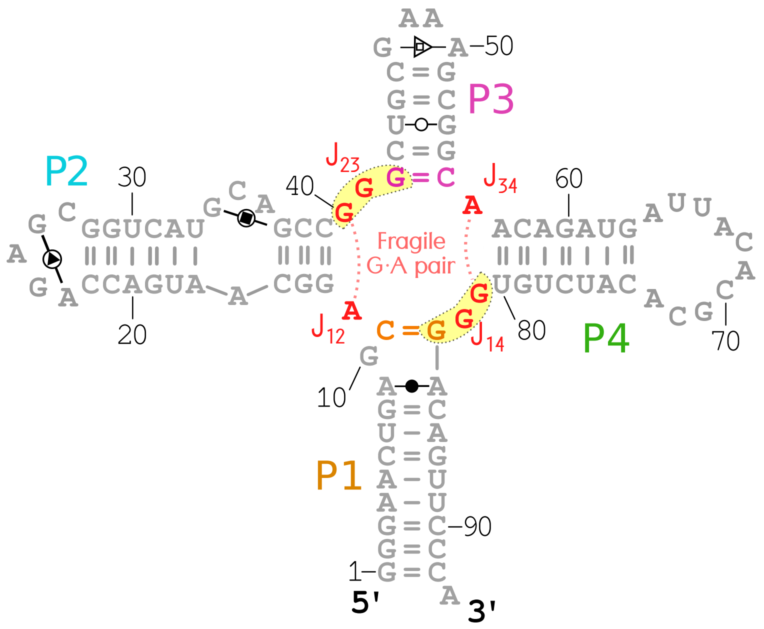


Figure S10: The terminal base pairs of the helices P1 (G83·C11: orange), P2 (G40·A12: cyan), P3 (G42·C55: mauve), and P4 (G81·A56: lime) in NRA secondary structure corresponding to state U. The junction loop ($J_{ij}$) nucleotides (red), the nucleotides corresponding to TC2 (yellow shade), and non-canonical H-bonds(black) are highlighted.

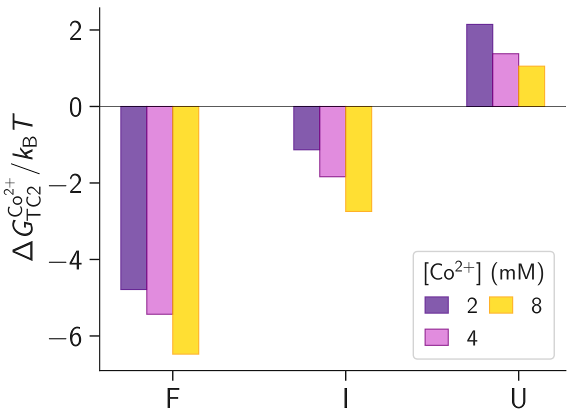


Figure S11: $\Delta G_{\mathrm{TC2}}^{\mathrm{Co}^{2+}}$ for different [$Co^{2+}$] at 300 K with a fixed background of [$Mg^{2+}$] = 8 mM and [$K^+$] = 150 mM for states F, I, and U.

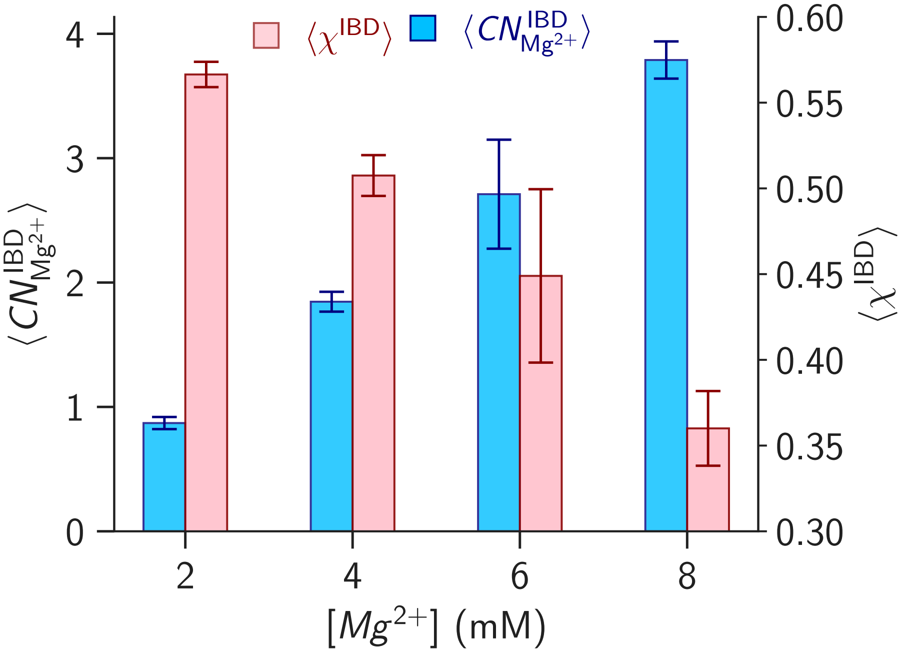


Figure S12: Average coordination number of $Mg^{2+}$ ion with the IBD nucleotides ($\langle CN^{IBD}_{Mg^{2+}} \rangle$) (blue) (left axis) as a function of [$Mg^{2+}$]. Structural overlap function ($\langle \chi^{IBD} \rangle$) (pink) (right axis) of IBD plotted as a function of [$Mg^{2+}$].

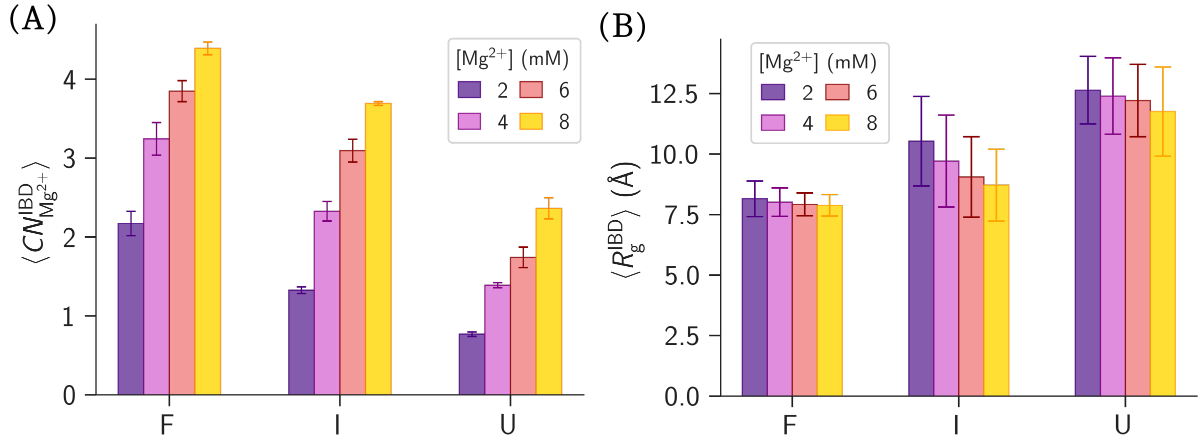


Figure S13: (A) Average coordination number of $Mg^{2+}$ ion with the IBD nucleotides ($\langle CN^{IBD}_{Mg^{2+}} \rangle$) for three states (U, I, and F) at different [$Mg^{2+}$]. (B) The average radius of gyration of IBD ($\langle R^{IBD}_g \rangle$) for three states (U, I, and F) at different [$Mg^{2+}$].

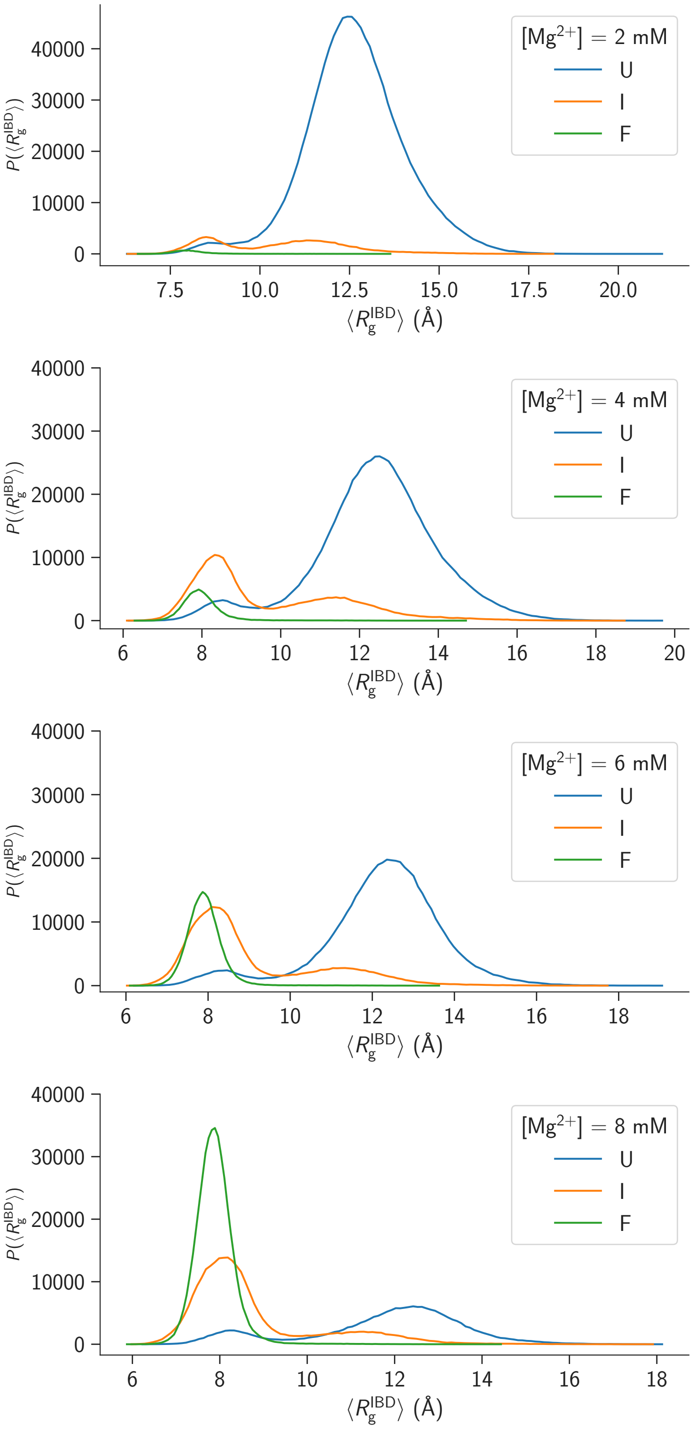


Figure S14: Probability distribution of average radius of gyration of IBD ($\langle R_g^{IBD} \rangle$) for three states (U, I and F) for $[Mg^{2+}]$ = 2, 4, 6 and 8 mM.

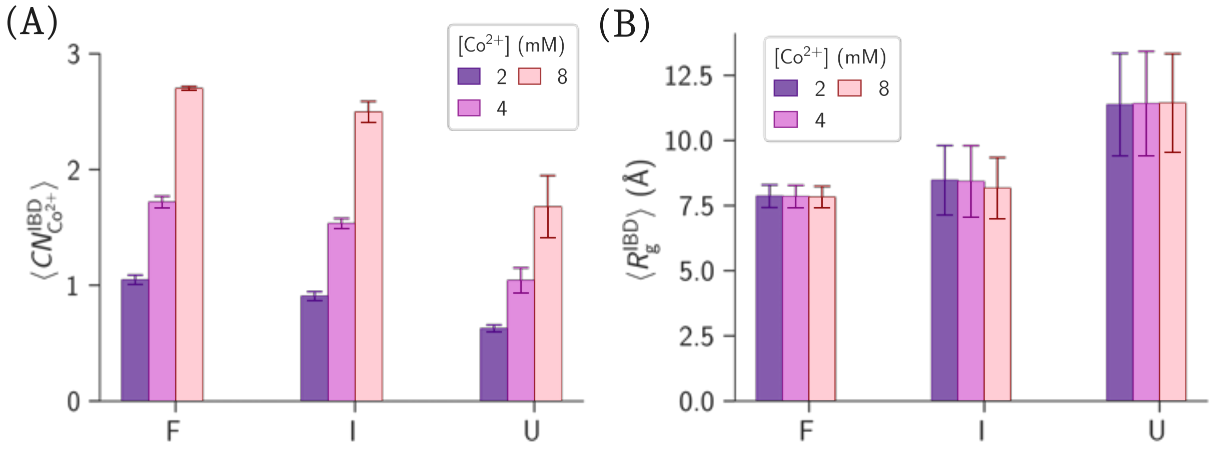


Figure S15: (A) Average coordination number of $Co^{2+}$ ion with the IBD nucleotides ($\langle CN^{\mathrm{IBD}}_{\mathrm{Mg}^{2+}} \rangle$) for three states (U, I, and F) at different [$Co^{2+}$]. (B) Average radius of gyration of IBD ($\langle R^{\mathrm{IBD}}_{\mathrm{g}} \rangle$) for three states (U, I, and F) at different [$Co^{2+}$]. The background of concentration of $Mg^{2+}$ is fixed at 8 mM.

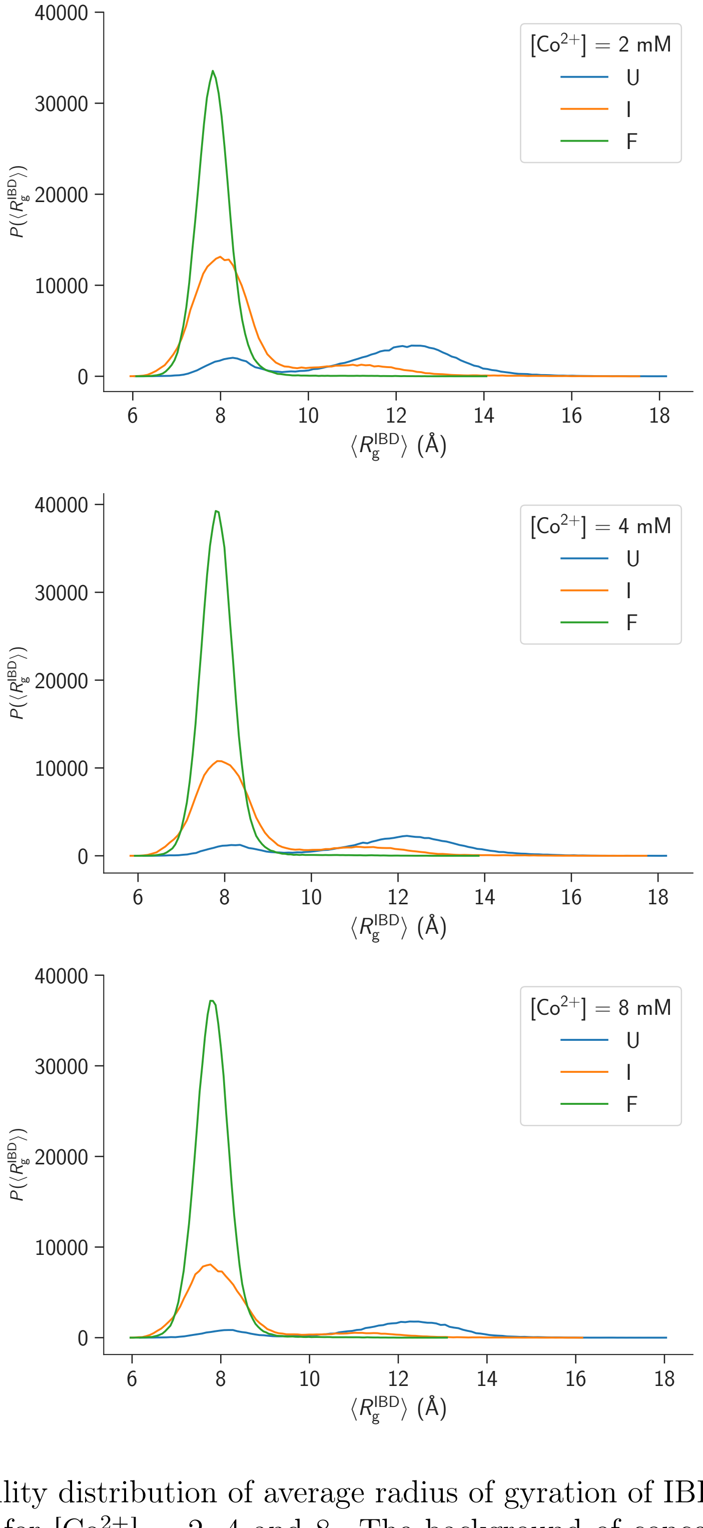


Figure S16: Probability distribution of average radius of gyration of IBD ($\langle R_g^{IBD} \rangle$) for three states (U, I and F) for $[Co^{2+}] = 2$, 4 and 8. The background of concentration of $Mg^{2+}$ is fixed at 8 mM.

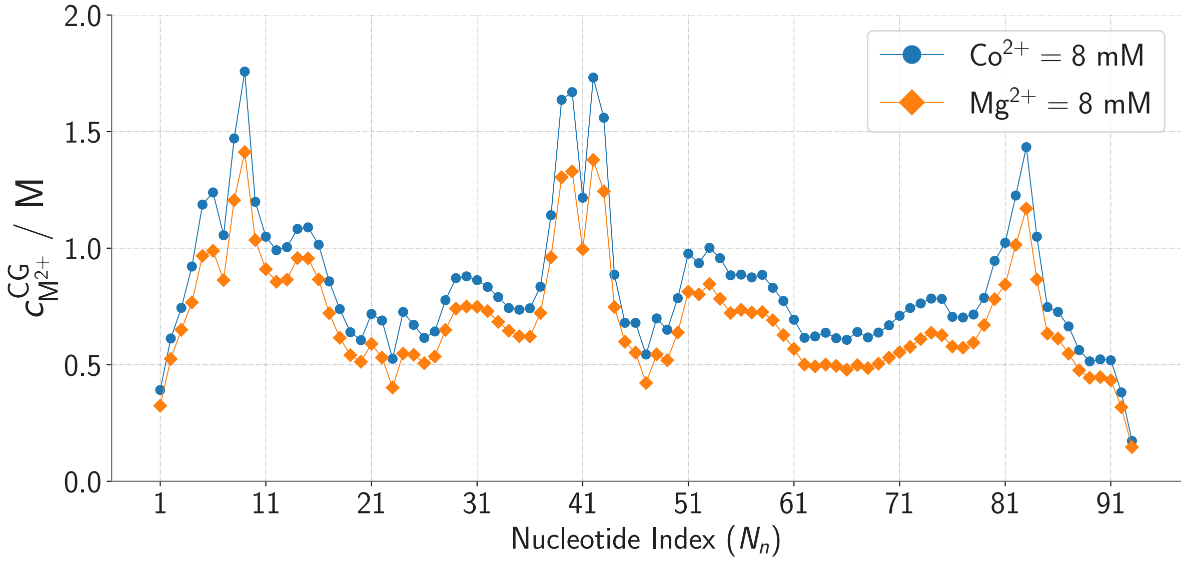


Figure S17: Local ion concentration, $c^{CG}_{M^{2+}}$, of $Co^{2+}$ and $Mg^{2+}$ around the NRA nucleotides in the F state for $[Co^{2+}] = [Mg^{2+}] = 8$ mM.

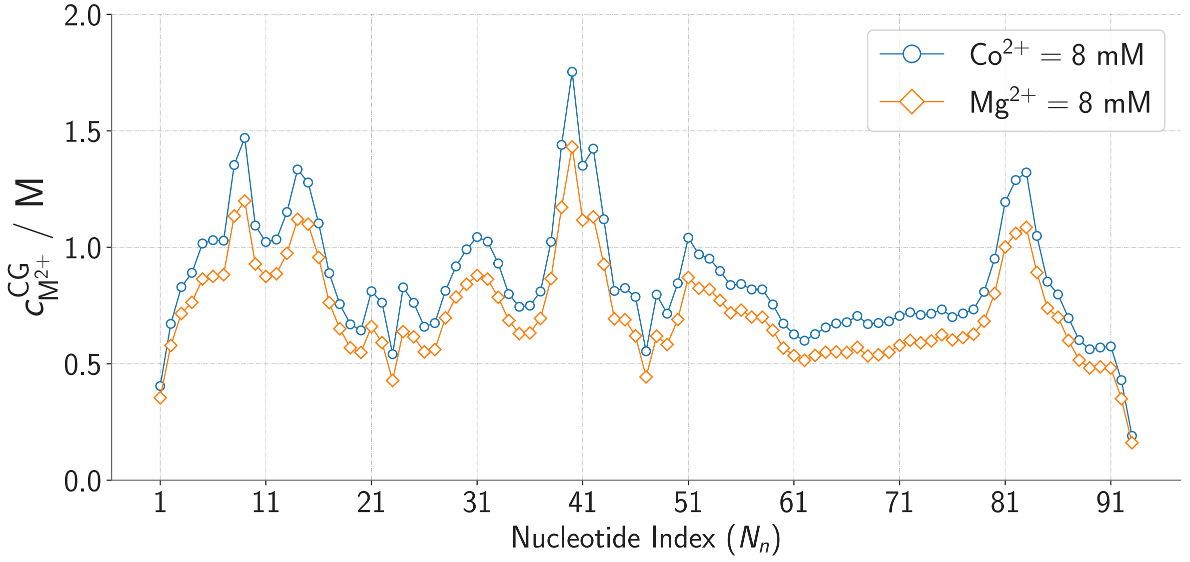


Figure S18: Local ion concentration, $c^{CG}_{M^{2+}}$, of $Co^{2+}$ and $Mg^{2+}$ around the NRA nucleotides in the I state for $[Co^{2+}] = [Mg^{2+}] = 8$ mM.

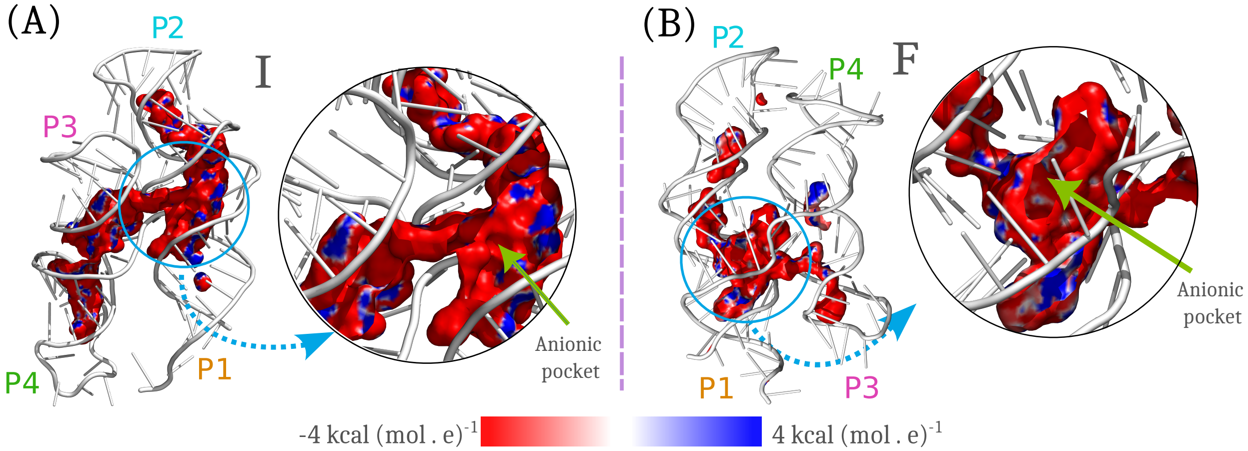


Figure S19: An anionic cavity present at the 4WJ (solid sky blue circle) in (A) state I (anionic pocket is wide open) and (B) state F (anionic pocket is compact). The inset shows the zoomed anionic cavity. The electrostatic potential surfaces are calculated by solving the Poisson-Boltzmann equation[35] using the PBEQ Solver server[36] present in CHARMM-GUI.[37] The parameters used for the calculation: (i) dielectric constant for the reference environment, $\varepsilon_R = 1.0$, (ii) dielectric constant for the RNA interior, $\varepsilon_P = 1.0$, (iii) solvent dielectric constant, $\varepsilon_W = 80$, and (iv) salt concentration, [salt] $= 0.15$ M. The electrostatic potential surface is projected on the cavity and visualized using the surface cavity module in PYMOL.[38]

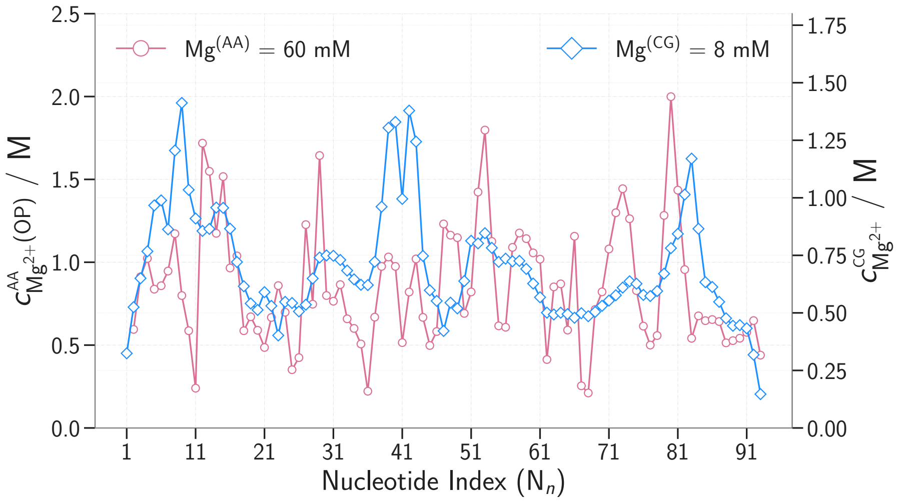


Figure S20: Local $Mg^{2+}$ ion concentration around the phosphate backbone ($c_{Mg^{2+}}$) computed from all-atom (AA) and coarse-grained (CG) simulations. The y-axis on the left and right shows $c_{Mg^{2+}}$ from all-atom and coarse-grained (CG) simulations, respectively. The concentration difference is due to the different box sizes in AA ($\sim$ 100 Å) and CG ($\sim$ 200 Å) simulations, but the number of ions in the box is similar in both cases (37 and 39, respectively).

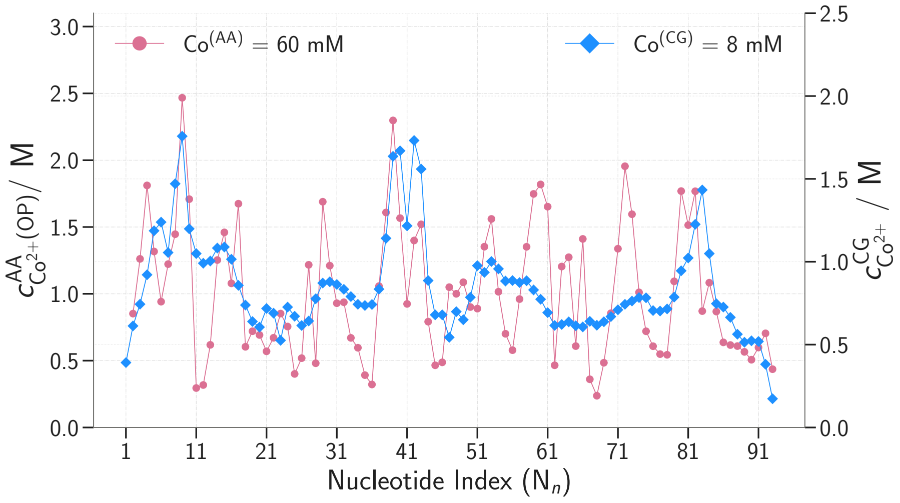


Figure S21: Local $Co^{2+}$ ion concentration around the phosphate backbone ($c_{Co^{2+}}$) from all-atom (AA) and coarse-grained (CG) simulations. The y-axis on the left and right shows $c_{Co^{2+}}$ from AA and CG simulations, respectively. The concentration difference is due to the different box sizes in AA ($\sim$ 100 Å) and CG ($\sim$ 200 Å) simulations, but the number of ions in the box is similar in both cases (37 and 39, respectively).

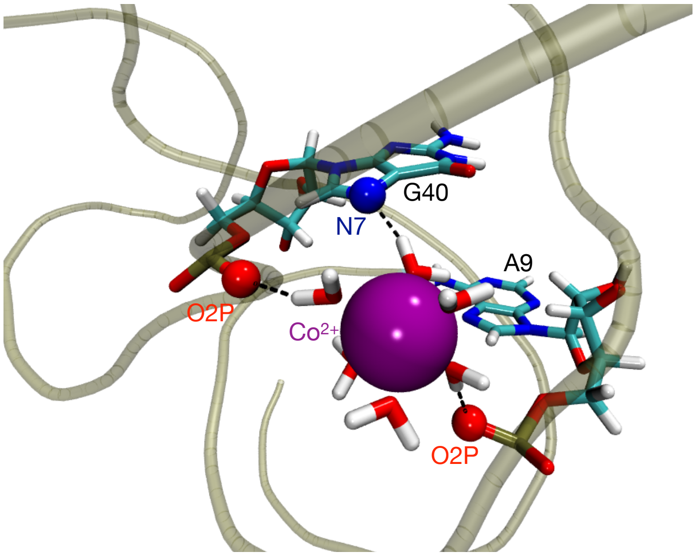


Figure S22: Representative snapshot of outer shell (OS) contacts between $Co^{2+}$ ion and N7 of G40 and phosphate oxygens (O2P) of A9 and G40 in the IBD.

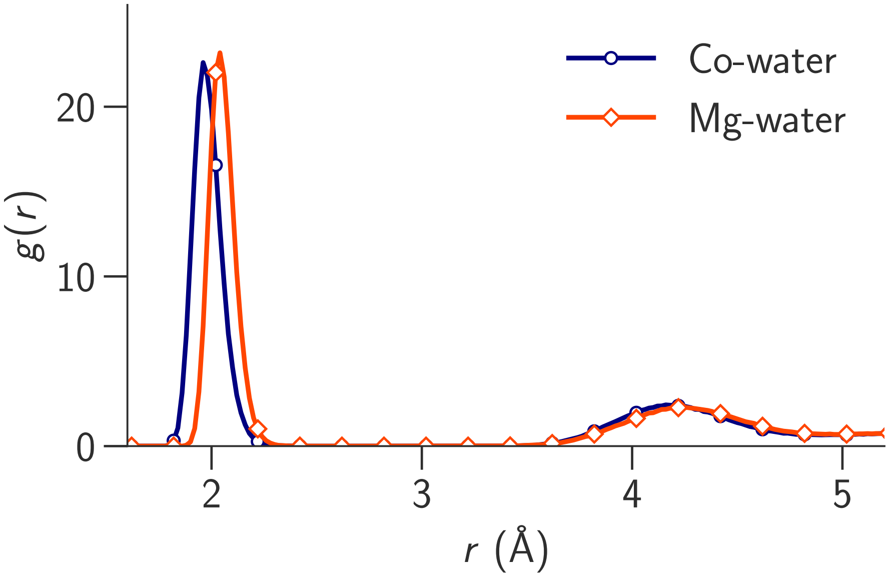


Figure S23: Radial distribution functions ($g(r)$) of $M^{2+}$ ions (M = Co, Mg) with the electronegative oxygen atoms of water molecules in water-$M^{2+}$ system.

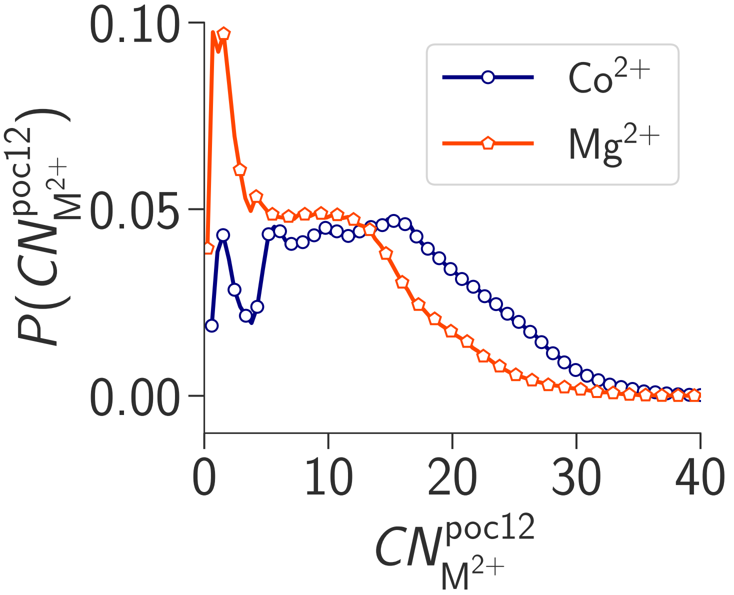


Figure S24: Probability distribution of the coordination number of $Co^{2+}$ (blue) and $Mg^{2+}$ (orange) ions ($P(CN^{\mathrm{poc12}}_{\mathrm{M}^{2+}})$) with the four conserved nucleotides (G41, G42, G82, G83) in poc12 pocket.

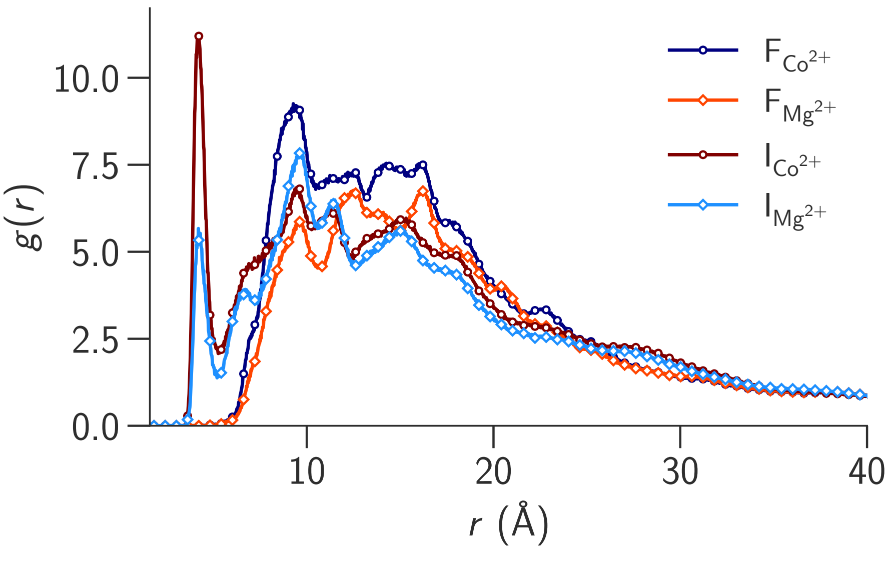


Figure S25: Radial distribution functions ($g(r)$) between $M^{2+}$ ions (M = Co, Mg) and N7 atoms of poc12 nucleotides (G41, G42, G82, G83) in the F-NRA and I-NRA systems.

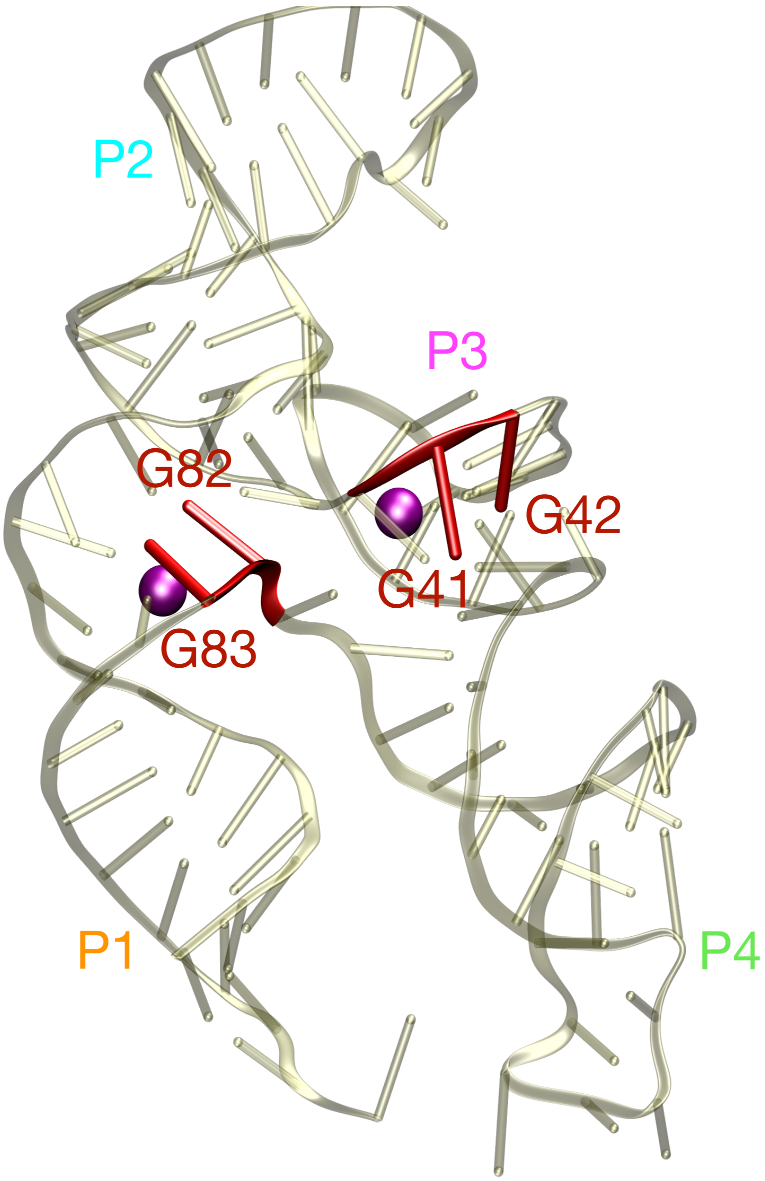


Figure S26: Representative conformation of the I state showing the interaction between poc12 nucleotides (red) and the two OS bound $Co^{2+}$ ions in purple.

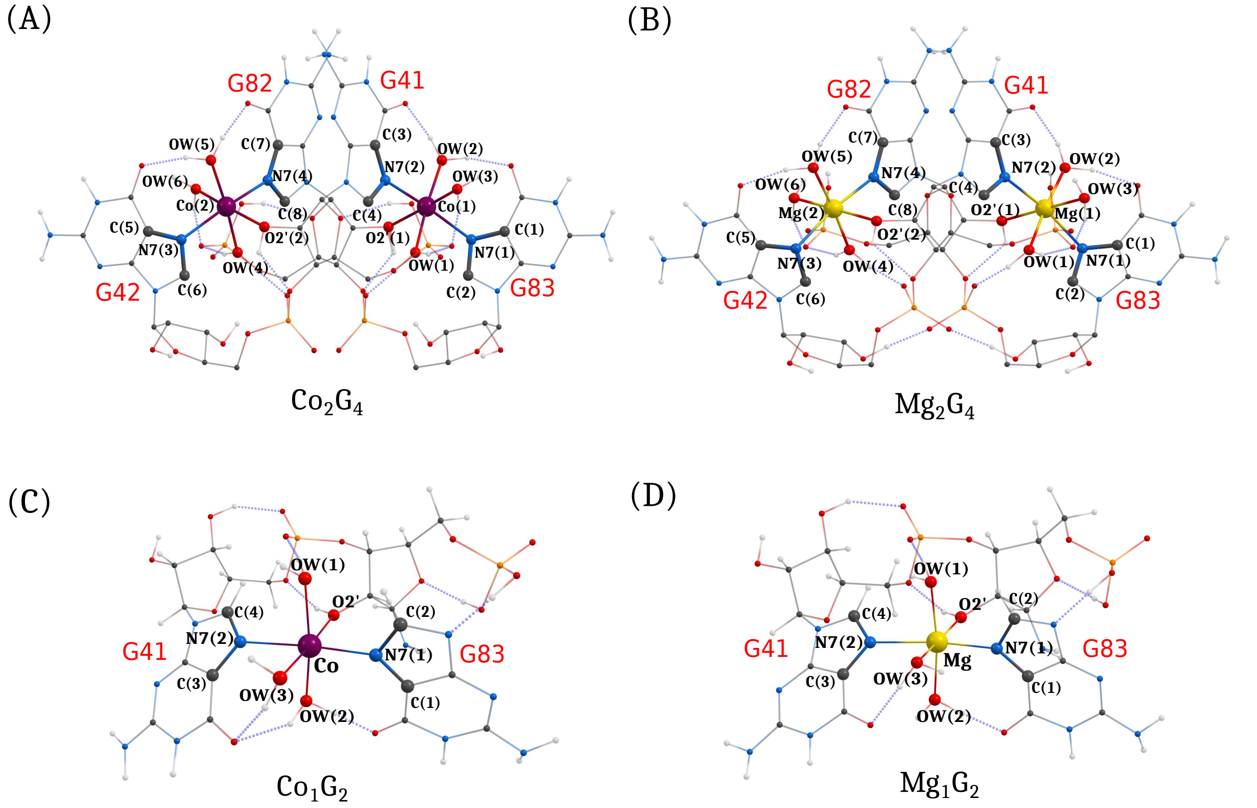


Figure S27: Geometry optimized structures of (A) $Co_2G_4$, (B) $Mg_2G_4$, (C) $Co_1G_2$, and (D) $Mg_1G_2$. In the structures, hydrogen (ivory), carbon (grey), nitrogen (blue), and oxygen (red) atoms around the $Co^{2+}$ (purple) and $Mg^{2+}$ (yellow) ions are highlighted and the rest of the atoms (including phosphorus in orange) are kept small for clarity.

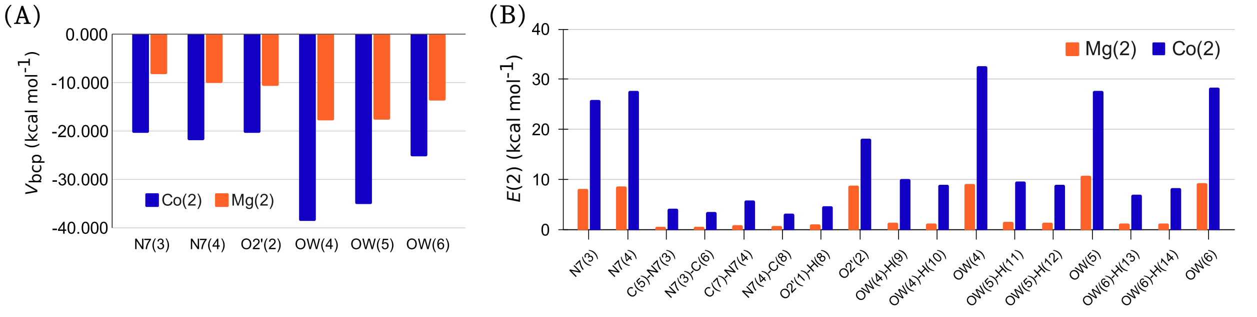


Figure S28: (A) $V_{\mathrm{bcp}}$ (in kcal mol$^{-1}$) of Mg(2) and Co(2) in $Mg_2G_4$ and $Co_2G_4$ complexes (Table S5). (B) Cumulative interaction energies, $E(2)$ (in kcal mol$^{-1}$), between every NBOs interacting with Mg(2) and Co(2) in $Mg_2G_4$ and $Co_2G_4$ complexes. Individual components are in Table S7.

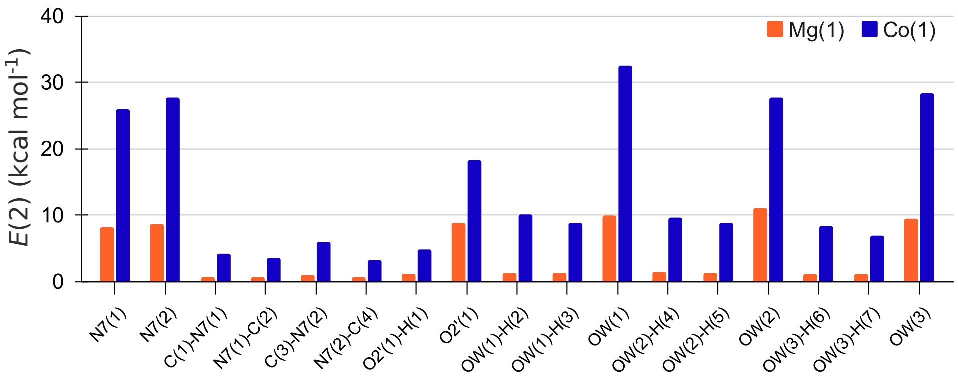


Figure S29: For all six co-ordinations (A) $V_{\mathrm{bcp}}$ (in kcal mol$^{-1}$) of Mg(1) and Co(1) in $Mg_2G_4$ and $Co_2G_4$ complexes (Table S5). (B) Cumulative interaction energies, $E(2)$ (in kcal mol$^{-1}$), between every NBOs interacting with Mg(2) and Co(2) in $Mg_2G_4$ and $Co_2G_4$ complexes. Individual components are in Table S6.

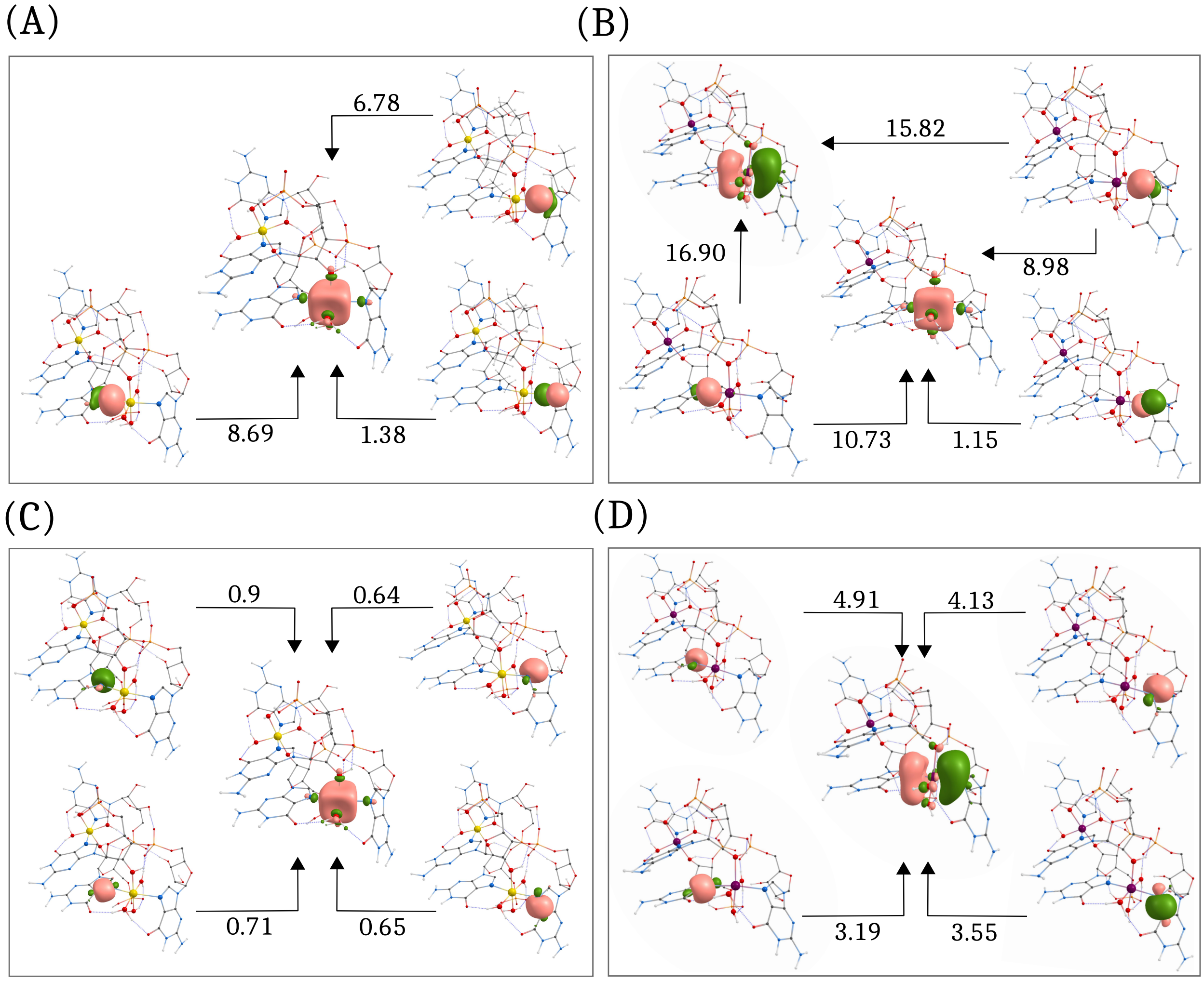


Figure S30: NBO orbital interactions in $M_2G_4$ complexes corresponding to the orbitals associated with N7 atoms of the conserved G residues with $M^{2+}$, M = Mg and Co. The interaction energies, $E(2)$ (in kcal mol$^{-1}$), are shown on top of the arrows. We are showing orbitals only for Mg(1) and Co(1) (Table. S6) for clarity. Mg(2) and Co(2) orbitals look almost similar to Mg(1) and Co(1), respectively. (A) $lp \rightarrow s$ interaction from two N7 atoms to $Mg^{2+}$ ion, (B) $lp \rightarrow s$ and $lp \rightarrow p$ interactions from two N7 atoms to $Co^{2+}$ ion. (C) $\pi \rightarrow s$ interaction from two C−N7 and two N7−C atoms to $Mg^{2+}$ ion, and (D) $\pi \rightarrow p$ interaction from two C−N7 and two N7−C atoms to $Co^{2+}$ ion.

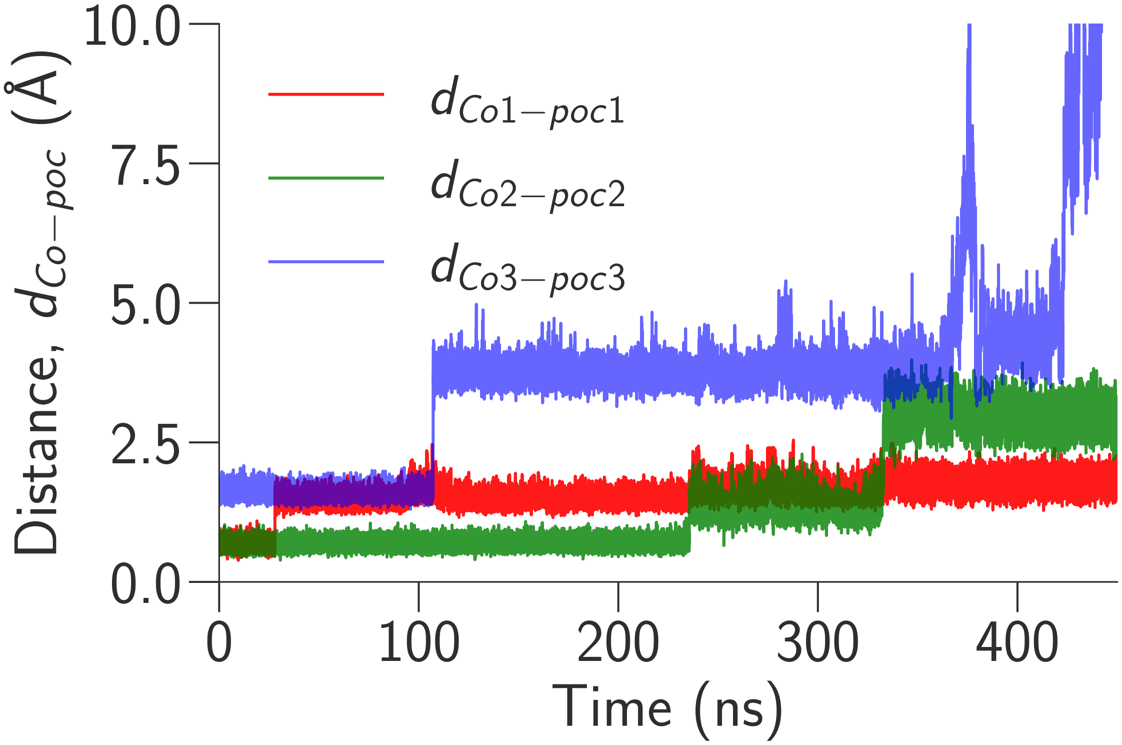


Figure S31: Data from the simulation of NRA(1,2,3) system. Distance between Co1 and the center of mass (COM) of N7 atom of G42, N7 atom of G82, O′ atom of G41 as a function of simulation time (red). Distance between Co2 and COM of N7 atom of G41, O′ atom of G82, and N7 atom of G83 as a function of simulation time (green). Distance between Co3 and COM of N7 atoms of A9, G40, as a function of simulation time (blue).

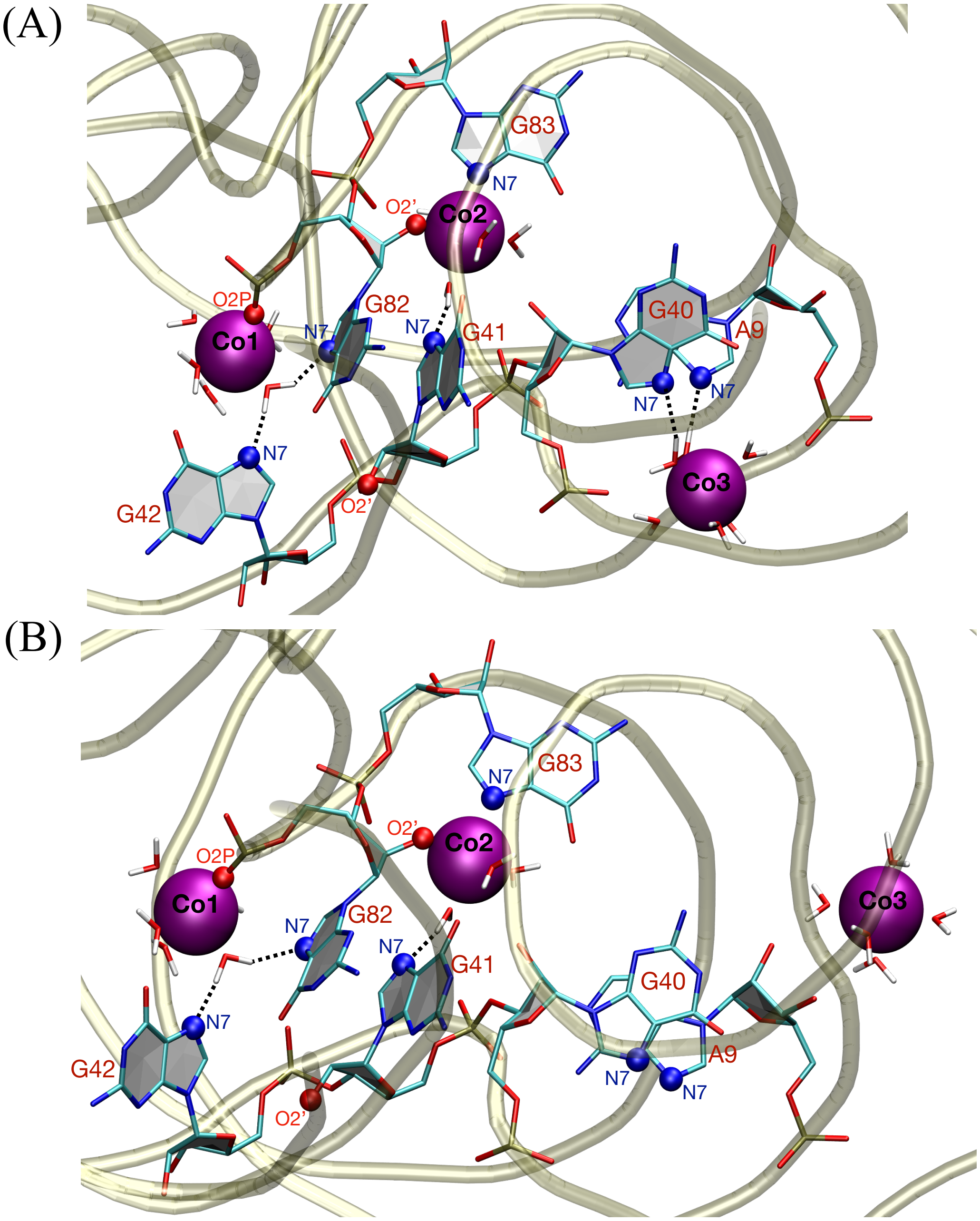


Figure S32: (A) Outer shell (OS) interactions of Co3 ion with the N7 atoms of A9 and G40. Co1 and Co2 ions are also forming OS interactions with the N7 atoms in the NRA(1,2,3). (B) The Co3 ion is displaced from its crystal position into the bulk. Co1 and Co2 ions are present in their crystal positions in NRA(1,2,3) system.

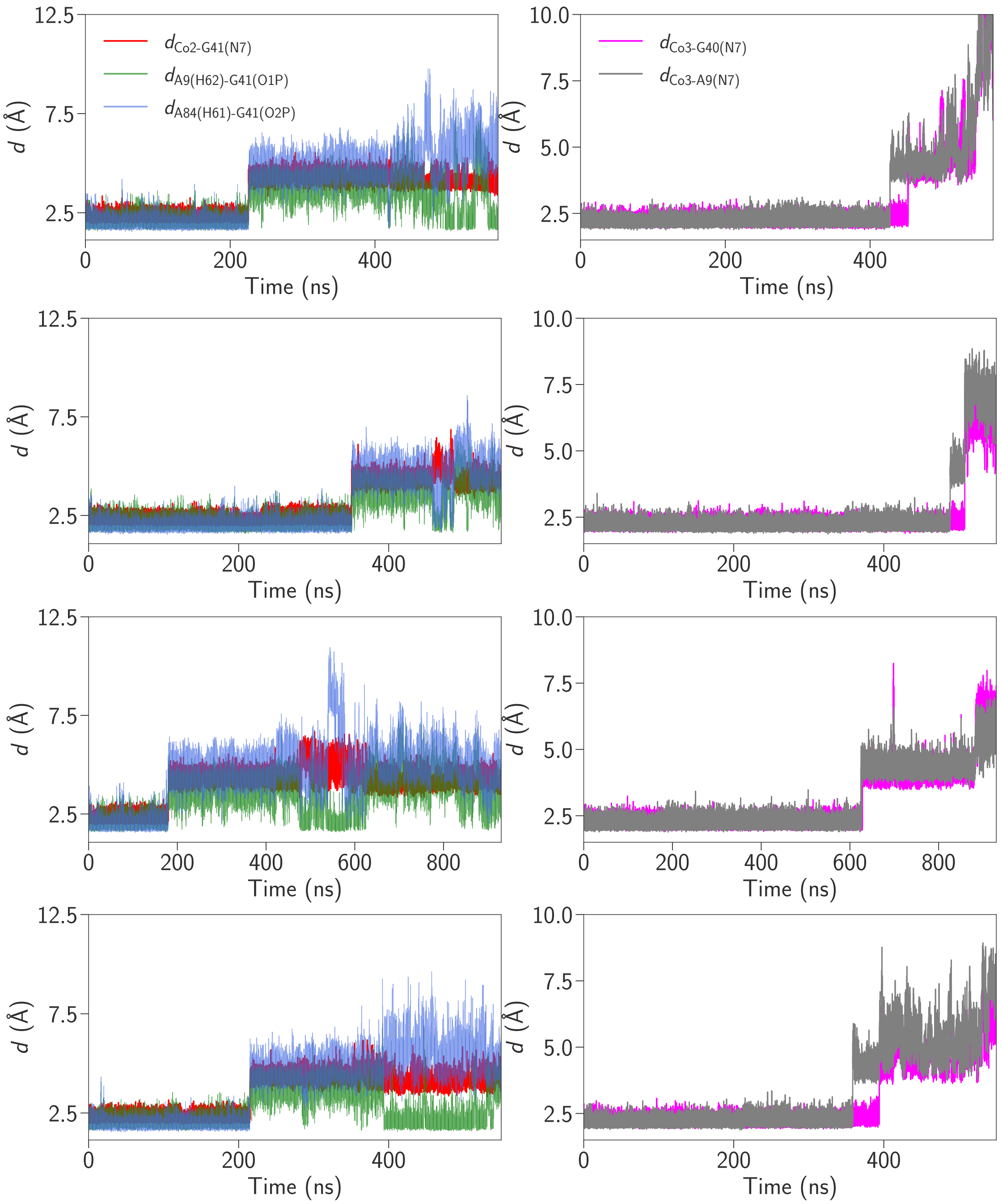


Figure S33: Two independent simulation trajectories with NRA(2,3) depicting the departure of Co3 from poc3 to bulk. (Left Column) Distance between Co2 and N7 atoms of G41, two base-phosphate H-bond linkages – A9-G41 ($d_{\text{A9(H62)-G41(O1P)}}$) and A84-G41 $d_{\text{A84(H61)-G41(O2P)}}$) – as a function of simulation time. (Right Column) Distance between Co3 and G40(N7) ($d_{\text{Co3-G40(N7)}}$), and distance between Co3 and A9(N7) ($d_{\text{Co3-A9(N7)}}$) as a function of simulation time.

(A) (B)

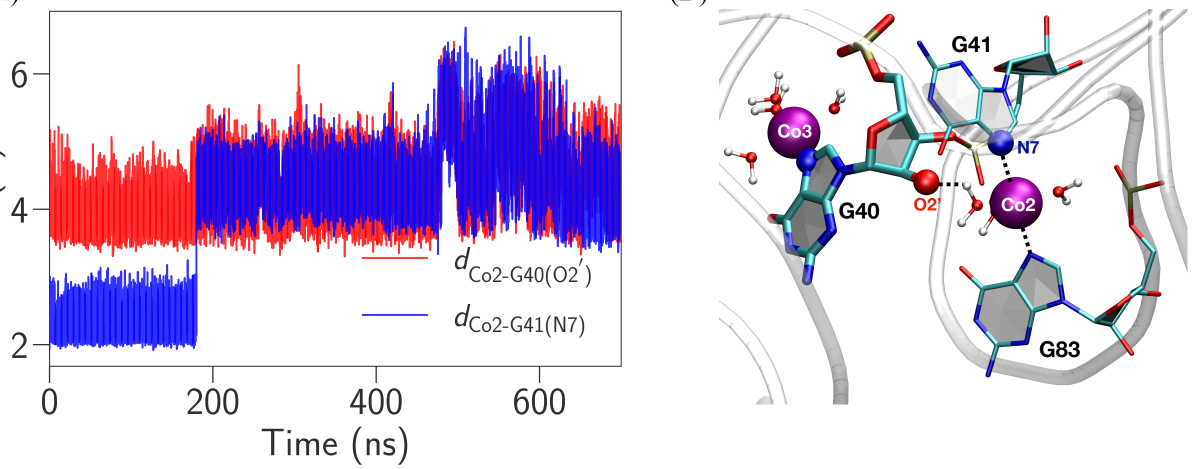


Figure S34: For NRA(2,3), (A) Distance between Co2 and G40(O2′) (red), and distance between Co2 and G41(N7) (blue) as a function of simulation time. (B) Representative snapshots of Co2 and Co3 ions forming IS and OS interactions with RNA atoms.

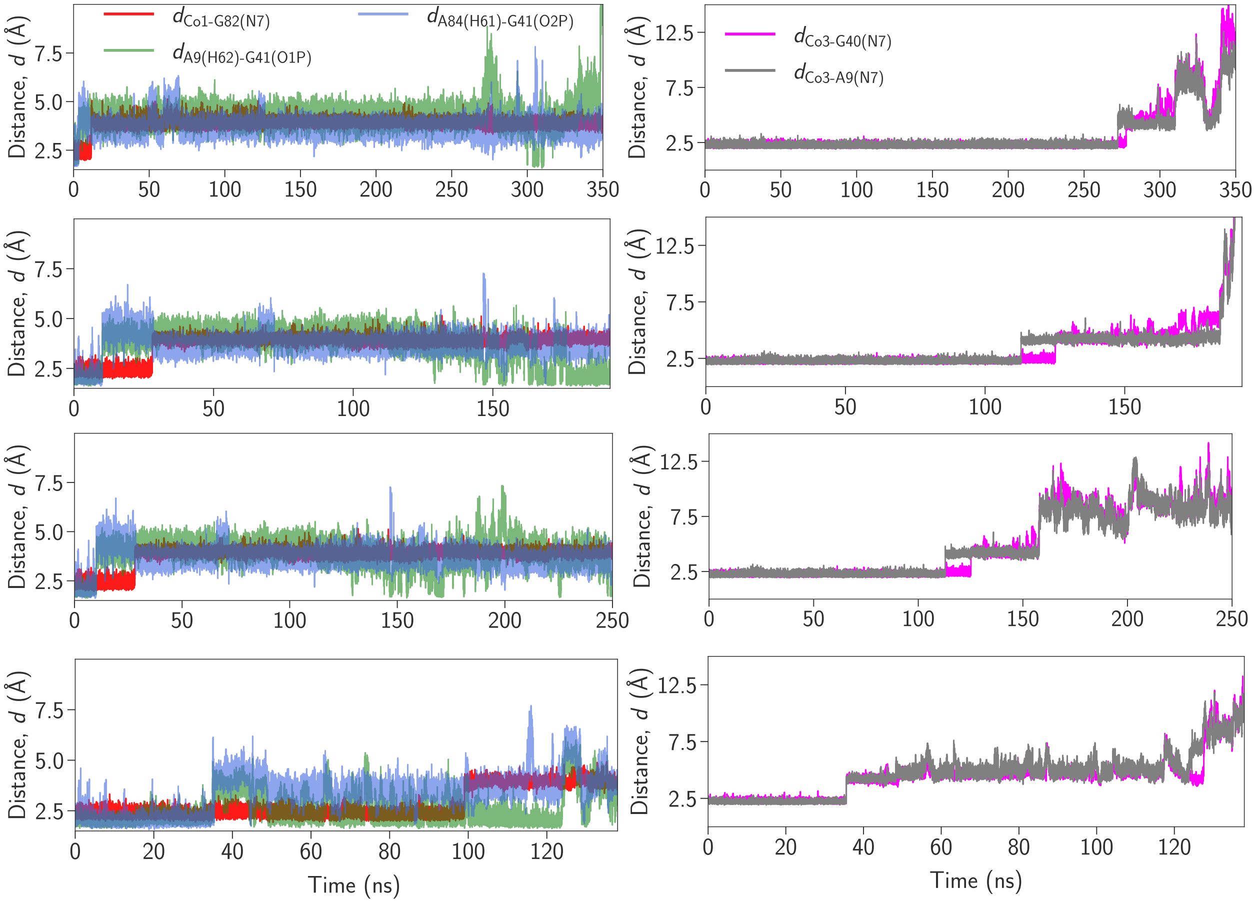


Figure S35: Four independent trajectories for NRA(1,3) depicting the departure of Co3 from poc3 to bulk. (Left Column) Distance between Co1 and G82(N7) ($d_{\text{Co1-G82(N7)}}$) and distance of two base-phosphate H-bond linkages – A9-G41 ($d_{\text{A9(H62)-G41(O1P)}}$) and A84-G41 ($d_{\text{A84(H61)-G41(O2P)}}$) – as a function of simulation time. (Right Column) Distance between Co3 and G40(N7) ($d_{\text{Co3-G40(N7)}}$), and distance between Co3 and A9(N7) ($d_{\text{Co3-A9(N7)}}$) as a function of simulation time to track the Co3 departure from poc3 to bulk.

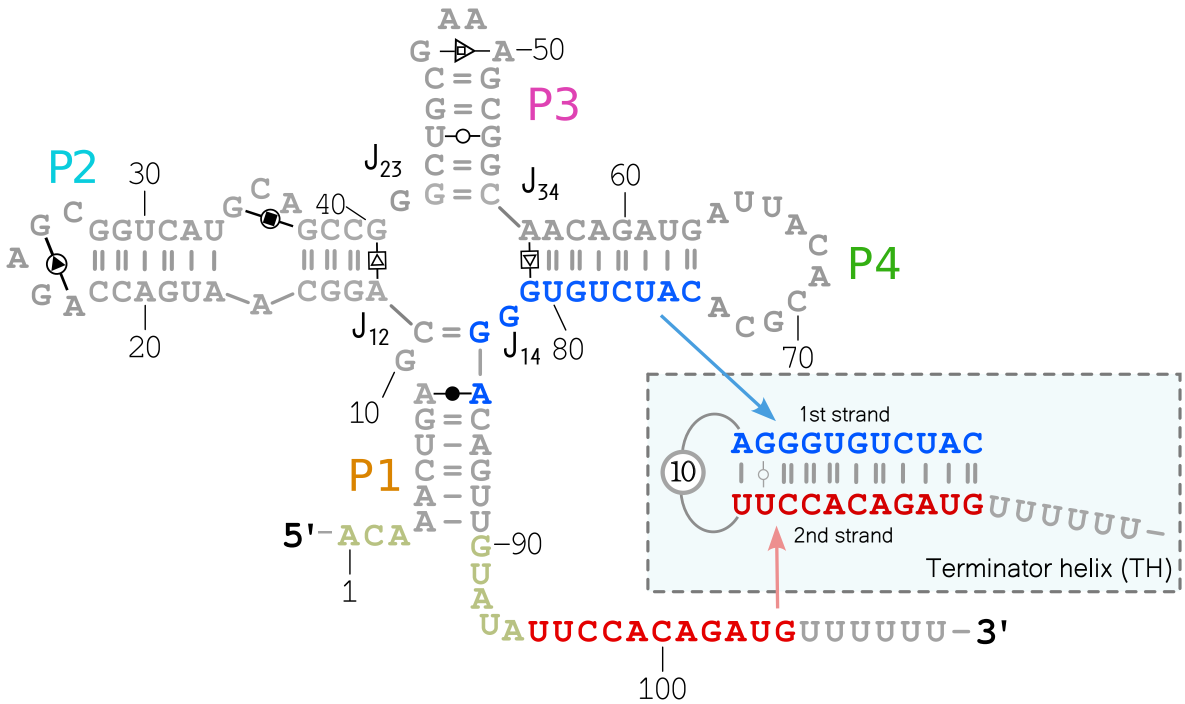


Figure S36: Sequence and predicted secondary structure of the *E. bacterium* NiCo riboswitch.[3] The full secondary structure includes the 1st (blue) and 2nd strand (blue) of the terminator helix. In the absence of transition metal ions, the 1st strand can break apart from the NRA and form the terminator helix (TH) (shown in the inset).

### 5 Movie S1

Transition from U→I→F. For the U→I transition, NRA undergoes a change in coaxial stacking arrangement. For the I→F transition, NRA undergoes a torsional twist at the 4WJ.

### 6 DFT Geometry Optimized Coordinates

## 6.1 1. $Co_2G_4$

```
Number of imaginary frequencies : 0

Electronic energy : HF=-9491.1676434

Zero-point correction=                          1.207294 (Hartree/Particle)

Thermal correction to Energy=                   1.302490

Thermal correction to Enthalpy=                 1.303434

Thermal correction to Gibbs Free Energy=        1.080339

Sum of electronic and zero-point Energies=         -9489.960349

Sum of electronic and thermal Energies=            -9489.865153

Sum of electronic and thermal Enthalpies=          -9489.864209

Sum of electronic and thermal Free Energies=       -9490.087304

............................................................

 Cartesian Coordinates

............................................................

   15         -3.989318   -0.265214   -3.383291

    8         -4.486371   -0.700027   -4.714131

    8         -4.855395   -0.300199   -2.112374

    8         -2.647441   -1.195897   -2.949883

    6         -1.693818   -1.502931   -3.964989

    1         -2.145776   -1.344763   -4.956784

    1         -1.416784   -2.566061   -3.871354

    6         -0.403449   -0.709780   -3.805528

    1          0.270113   -0.968138   -4.640687

    8         -0.646427    0.717900   -3.825253

    6          0.094200    1.404771   -2.837604
```

| | | | |
|---|---|---|---|
| 1 | 0.700799 | 2.205460 | -3.283552 |
| 7 | -0.813262 | 2.040986 | -1.882760 |
| 6 | -1.627442 | 1.335808 | -1.038904 |
| 1 | -1.739313 | 0.259997 | -1.106266 |
| 7 | -2.255813 | 2.110729 | -0.182516 |
| 6 | -1.873833 | 3.398755 | -0.499953 |
| 6 | -2.230222 | 4.658963 | 0.075477 |
| 8 | -2.985030 | 4.923929 | 1.007730 |
| 7 | -1.540259 | 5.711628 | -0.573777 |
| 1 | -1.625012 | 6.605982 | -0.096320 |
| 6 | -0.625936 | 5.571168 | -1.590822 |
| 7 | 0.035835 | 6.711466 | -1.983754 |
| 1 | -0.507240 | 7.567813 | -2.041546 |
| 1 | 0.646951 | 6.554872 | -2.780724 |
| 7 | -0.320852 | 4.416955 | -2.124538 |
| 6 | -0.956594 | 3.369642 | -1.560226 |
| 6 | 0.319112 | -0.982640 | -2.485690 |
| 1 | -0.400026 | -1.292427 | -1.710156 |
| 6 | 0.956989 | 0.366236 | -2.105564 |
| 1 | 0.911774 | 0.517645 | -1.027846 |
| 8 | 2.310166 | 0.404325 | -2.526082 |
| 1 | 2.508166 | -0.536152 | -2.748535 |
| 8 | 1.334113 | -1.948036 | -2.641364 |
| 15 | 1.422241 | -3.026098 | -1.327167 |
| 8 | 0.433499 | -4.116055 | -1.490009 |
| 8 | 1.501794 | -2.108918 | -0.099183 |
| 8 | 2.963477 | -3.504689 | -1.677879 |

| | | | |
|---|---|---|---|
| 6 | 3.398801 | -4.744363 | -1.115800 |
| 1 | 3.595213 | -5.450876 | -1.937749 |
| 1 | 2.614814 | -5.176805 | -0.473945 |
| 6 | 4.655531 | -4.554976 | -0.296701 |
| 1 | 4.907917 | -5.514941 | 0.187338 |
| 8 | 5.742918 | -4.160100 | -1.159950 |
| 6 | 6.591472 | -3.261586 | -0.513762 |
| 1 | 7.641397 | -3.585748 | -0.552786 |
| 7 | 6.520984 | -1.942722 | -1.190643 |
| 6 | 5.386886 | -1.373031 | -1.708651 |
| 1 | 4.503246 | -1.979376 | -1.906563 |
| 7 | 5.502940 | -0.063687 | -1.857639 |
| 6 | 6.790413 | 0.237449 | -1.443591 |
| 6 | 7.453613 | 1.484041 | -1.256809 |
| 8 | 7.039761 | 2.638719 | -1.423765 |
| 7 | 8.756674 | 1.294754 | -0.747770 |
| 1 | 9.282392 | 2.156848 | -0.631323 |
| 6 | 9.311145 | 0.087181 | -0.379366 |
| 7 | 10.591550 | 0.101366 | 0.083759 |
| 1 | 10.981492 | 0.946143 | 0.481719 |
| 1 | 10.912892 | -0.775243 | 0.478860 |
| 7 | 8.666724 | -1.048303 | -0.490365 |
| 6 | 7.433580 | -0.931025 | -1.015882 |
| 6 | 4.602882 | -3.455073 | 0.767447 |
| 1 | 4.066776 | -2.599471 | 0.353311 |
| 6 | 6.094788 | -3.118488 | 0.933096 |
| 1 | 6.244900 | -2.101552 | 1.333056 |

| | | | |
|---|---|---|---|
| 8 | 6.711841 | -4.099381 | 1.727046 |
| 1 | 6.056165 | -4.311899 | 2.414283 |
| 8 | 4.485429 | -0.695159 | 4.713644 |
| 8 | 4.853509 | -0.299486 | 2.111110 |
| 8 | 2.645741 | -1.193415 | 2.950780 |
| 6 | 1.692408 | -1.498899 | 3.966597 |
| 1 | 2.144551 | -1.339098 | 4.958046 |
| 1 | 1.415382 | -2.562185 | 3.874740 |
| 6 | 0.402062 | -0.705906 | 3.806027 |
| 1 | -0.271648 | -0.963094 | 4.641418 |
| 8 | 0.645241 | 0.721782 | 3.824023 |
| 6 | -0.094676 | 1.407476 | 2.834945 |
| 1 | -0.701209 | 2.208950 | 3.279558 |
| 7 | 0.813368 | 2.042109 | 1.879723 |
| 6 | 1.625833 | 1.335661 | 1.035318 |
| 1 | 1.734865 | 0.259503 | 1.101627 |
| 7 | 2.256473 | 2.109762 | 0.179889 |
| 6 | 1.878039 | 3.398472 | 0.498701 |
| 6 | 2.237741 | 4.658282 | -0.075467 |
| 8 | 2.993331 | 4.922208 | -1.007393 |
| 7 | 1.550245 | 5.712082 | 0.574548 |
| 1 | 1.637267 | 6.606556 | 0.097777 |
| 6 | 0.635052 | 5.572950 | 1.590964 |
| 7 | -0.024892 | 6.714361 | 1.983935 |
| 1 | 0.519873 | 7.569589 | 2.042679 |
| 1 | -0.636520 | 6.558455 | 2.780667 |
| 7 | 0.327201 | 4.419087 | 2.123792 |

| | | | |
|---|---|---|---|
| 6 | 0.960547 | 3.370717 | 1.558765 |
| 6 | -0.320260 | -0.980404 | 2.486381 |
| 1 | 0.398963 | -1.291698 | 1.711541 |
| 6 | -0.957499 | 0.368183 | 2.104070 |
| 1 | -0.912160 | 0.517907 | 1.026077 |
| 8 | -2.310693 | 0.407418 | 2.524435 |
| 1 | -2.509099 | -0.532685 | 2.748218 |
| 8 | -1.335708 | -1.945208 | 2.643209 |
| 15 | -1.424767 | -3.024219 | 1.329732 |
| 8 | -0.436734 | -4.114715 | 1.493162 |
| 8 | -1.503887 | -2.107827 | 0.101116 |
| 8 | -2.966203 | -3.501591 | 1.680996 |
| 6 | -3.402172 | -4.741841 | 1.120760 |
| 1 | -3.598977 | -5.447048 | 1.943738 |
| 1 | -2.618413 | -5.175602 | 0.479528 |
| 6 | -4.658786 | -4.552940 | 0.301386 |
| 1 | -4.911596 | -5.513427 | -0.181406 |
| 8 | -5.745942 | -4.156305 | 1.163973 |
| 6 | -6.593913 | -3.258042 | 0.516593 |
| 1 | -7.644022 | -3.581591 | 0.555900 |
| 7 | -6.522637 | -1.938540 | 1.191957 |
| 6 | -5.388222 | -1.369041 | 1.709552 |
| 1 | -4.504941 | -1.975699 | 1.908135 |
| 7 | -5.503584 | -0.059535 | 1.857497 |
| 6 | -6.790754 | 0.242042 | 1.442814 |
| 6 | -7.453122 | 1.488935 | 1.254868 |
| 8 | -7.038700 | 2.643456 | 1.421235 |

| | | | |
|---|---|---|---|
| 7 | -8.756110 | 1.300002 | 0.745458 |
| 1 | -9.281399 | 2.162273 | 0.628384 |
| 6 | -9.311289 | 0.092424 | 0.378131 |
| 7 | -10.591607 | 0.107036 | -0.085330 |
| 1 | -10.980425 | 0.951576 | -0.484927 |
| 1 | -10.913236 | -0.769759 | -0.479810 |
| 7 | -8.667679 | -1.043379 | 0.490418 |
| 6 | -7.434572 | -0.926409 | 1.016102 |
| 6 | -4.605362 | -3.454533 | -0.764302 |
| 1 | -4.068490 | -2.598814 | -0.351399 |
| 6 | -6.096995 | -3.117055 | -0.930416 |
| 1 | -6.246532 | -2.100528 | -1.331603 |
| 8 | -6.714705 | -4.098443 | -1.723252 |
| 1 | -6.058988 | -4.312495 | -2.409982 |
| 27 | 3.805129 | 1.144941 | -1.073305 |
| 27 | -3.805128 | 1.147331 | 1.071076 |
| 8 | 3.404000 | -0.533611 | -0.028983 |
| 1 | 3.928495 | -0.585282 | 0.819002 |
| 1 | 2.614894 | -1.206680 | -0.033356 |
| 8 | 4.371164 | 2.841370 | -2.072447 |
| 1 | 5.343898 | 2.935149 | -1.944488 |
| 1 | 3.933324 | 3.662974 | -1.743484 |
| 8 | 5.023395 | 1.787303 | 0.569151 |
| 1 | 5.112226 | 1.075777 | 1.273359 |
| 1 | 5.869680 | 2.223921 | 0.418390 |
| 8 | -3.405688 | -0.532326 | 0.028088 |
| 1 | -3.931124 | -0.585104 | -0.819199 |

| | | | |
|---|---|---|---|
| 1 | -2.616747 | -1.205718 | 0.033351 |
| 8 | -4.369831 | 2.844988 | 2.068377 |
| 1 | -5.342325 | 2.940080 | 1.939681 |
| 1 | -3.930147 | 3.666160 | 1.740964 |
| 8 | -5.023273 | 1.788205 | -0.572082 |
| 1 | -5.869340 | 2.225758 | -0.422940 |
| 1 | -5.112203 | 1.075967 | -1.275560 |
| 15 | 3.987936 | -0.262342 | 3.382317 |
| 8 | 3.349933 | 1.249471 | 3.381406 |
| 1 | 2.513958 | 1.265916 | 3.881779 |
| 8 | 4.017728 | -3.880697 | 1.969999 |
| 1 | 3.478988 | -3.146583 | 2.310552 |
| 8 | -4.020514 | -3.882417 | -1.966263 |
| 1 | -3.481242 | -3.149197 | -2.307921 |
| 8 | -3.351109 | 1.246504 | -3.384370 |
| 1 | -2.515118 | 1.262376 | -3.884705 |

## 6.2 2. $Mg_2G_4$

Number of imaginary frequencies : 0

Electronic energy : HF=-7126.1653067

Zero-point correction= 1.209953 (Hartree/Particle)

Thermal correction to Energy= 1.304648

Thermal correction to Enthalpy= 1.305592

Thermal correction to Gibbs Free Energy= 1.084629

Sum of electronic and zero-point Energies= -7124.955354

Sum of electronic and thermal Energies= -7124.860659

Sum of electronic and thermal Enthalpies= -7124.859715

Sum of electronic and thermal Free Energies= -7125.080678

..............................................................

Cartesian Coordinates

..............................................................

| | | | |
|---|---|---|---|
| 15 | -4.179785 | 0.292768 | -3.381078 |
| 8 | -4.734161 | 0.252717 | -4.760965 |
| 8 | -4.807408 | -0.367292 | -2.149233 |
| 8 | -2.622407 | -0.245719 | -3.336840 |
| 6 | -1.815507 | -0.305090 | -4.501550 |
| 1 | -2.252347 | 0.291805 | -5.316193 |
| 1 | -1.742963 | -1.351821 | -4.842675 |
| 6 | -0.421741 | 0.181542 | -4.153265 |
| 1 | 0.264697 | -0.026832 | -4.991982 |
| 8 | -0.435085 | 1.597336 | -3.896201 |
| 6 | 0.191226 | 1.931056 | -2.690695 |
| 1 | 0.906130 | 2.751957 | -2.843176 |
| 7 | -0.783898 | 2.417494 | -1.704737 |
| 6 | -1.611346 | 1.619909 | -0.965806 |
| 1 | -1.663221 | 0.545532 | -1.084384 |
| 7 | -2.354672 | 2.302275 | -0.119980 |
| 6 | -2.029927 | 3.627450 | -0.337929 |
| 6 | -2.519492 | 4.826666 | 0.271624 |
| 8 | -3.355300 | 4.977361 | 1.155786 |
| 7 | -1.886583 | 5.961190 | -0.299625 |
| 1 | -2.118313 | 6.838214 | 0.157873 |
| 6 | -0.915340 | 5.938521 | -1.272363 |
| 7 | -0.362285 | 7.140478 | -1.636672 |

| | | | |
|---|---|---|---|
| 1 | -0.935400 | 7.972860 | -1.554215 |
| 1 | 0.182173 | 7.085018 | -2.492357 |
| 7 | -0.471696 | 4.833492 | -1.816725 |
| 6 | -1.049373 | 3.715083 | -1.332829 |
| 6 | 0.110304 | -0.467069 | -2.872128 |
| 1 | -0.743526 | -0.772525 | -2.252465 |
| 6 | 0.874113 | 0.660376 | -2.159914 |
| 1 | 0.784737 | 0.558868 | -1.077565 |
| 8 | 2.249121 | 0.631591 | -2.512988 |
| 1 | 2.395661 | -0.219814 | -2.973753 |
| 8 | 0.939563 | -1.575827 | -3.134081 |
| 15 | 0.898583 | -2.678772 | -1.854777 |
| 8 | -0.352318 | -3.496360 | -1.904704 |
| 8 | 1.265920 | -1.854968 | -0.633554 |
| 8 | 2.204541 | -3.522152 | -2.359694 |
| 6 | 2.438978 | -4.849611 | -1.879819 |
| 1 | 2.677094 | -5.473402 | -2.754460 |
| 1 | 1.531576 | -5.250665 | -1.400763 |
| 6 | 3.581528 | -4.892535 | -0.885592 |
| 1 | 3.692073 | -5.933977 | -0.533429 |
| 8 | 4.796095 | -4.471335 | -1.522404 |
| 6 | 5.592998 | -3.725922 | -0.645722 |
| 1 | 6.605009 | -4.146736 | -0.552151 |
| 7 | 5.749898 | -2.368705 | -1.208042 |
| 6 | 4.801940 | -1.680323 | -1.912748 |
| 1 | 3.874460 | -2.166230 | -2.212226 |
| 7 | 5.145684 | -0.420611 | -2.131198 |

| | | | |
|---|---|---|---|
| 6 | 6.407205 | -0.293690 | -1.573656 |
| 6 | 7.274811 | 0.841003 | -1.485609 |
| 8 | 7.138841 | 1.978229 | -1.934071 |
| 7 | 8.440896 | 0.509231 | -0.760544 |
| 1 | 9.106171 | 1.272762 | -0.677682 |
| 6 | 8.718147 | -0.713560 | -0.186594 |
| 7 | 9.910681 | -0.837155 | 0.465994 |
| 1 | 10.341233 | -0.015204 | 0.871632 |
| 1 | 10.003577 | -1.692352 | 1.003296 |
| 7 | 7.915852 | -1.745087 | -0.279925 |
| 6 | 6.787531 | -1.495425 | -0.971909 |
| 6 | 3.422500 | -3.992793 | 0.351810 |
| 1 | 2.922575 | -3.061730 | 0.071033 |
| 6 | 4.893359 | -3.718564 | 0.726649 |
| 1 | 5.035621 | -2.757871 | 1.254087 |
| 8 | 5.380441 | -4.804376 | 1.474971 |
| 1 | 4.613063 | -5.101311 | 1.997350 |
| 8 | 4.736806 | 0.246368 | 4.759777 |
| 8 | 4.807099 | -0.370397 | 2.147248 |
| 8 | 2.623203 | -0.247898 | 3.336616 |
| 6 | 1.816716 | -0.305382 | 4.501803 |
| 1 | 2.253083 | 0.293841 | 5.314986 |
| 1 | 1.745600 | -1.351385 | 4.845398 |
| 6 | 0.422311 | 0.179332 | 4.153474 |
| 1 | -0.263683 | -0.030293 | 4.992238 |
| 8 | 0.433693 | 1.595305 | 3.897317 |
| 6 | -0.191963 | 1.928956 | 2.691542 |

| | | | |
|---|---|---|---|
| 1 | -0.907380 | 2.749439 | 2.843824 |
| 7 | 0.783437 | 2.416308 | 1.706247 |
| 6 | 1.611642 | 1.619406 | 0.967455 |
| 1 | 1.663979 | 0.545015 | 1.085629 |
| 7 | 2.354825 | 2.302393 | 0.122023 |
| 6 | 2.029190 | 3.627335 | 0.340107 |
| 6 | 2.518336 | 4.827021 | -0.268877 |
| 8 | 3.354319 | 4.978405 | -1.152777 |
| 7 | 1.884566 | 5.961007 | 0.302408 |
| 1 | 2.115484 | 6.838237 | -0.155110 |
| 6 | 0.913150 | 5.937534 | 1.274969 |
| 7 | 0.359513 | 7.139153 | 1.639562 |
| 1 | 0.932787 | 7.971581 | 1.558554 |
| 1 | -0.185645 | 7.082875 | 2.494756 |
| 7 | 0.469884 | 4.832110 | 1.818828 |
| 6 | 1.048188 | 3.714143 | 1.334659 |
| 6 | -0.109440 | -0.469365 | 2.872227 |
| 1 | 0.744323 | -0.774657 | 2.252401 |
| 6 | -0.873790 | 0.658003 | 2.160155 |
| 1 | -0.784227 | 0.556773 | 1.077798 |
| 8 | -2.248818 | 0.628296 | 2.513186 |
| 1 | -2.394769 | -0.223522 | 2.973429 |
| 8 | -0.938519 | -1.578332 | 3.134148 |
| 15 | -0.899129 | -2.680586 | 1.854240 |
| 8 | 0.351241 | -3.499080 | 1.902535 |
| 8 | -1.267085 | -1.855901 | 0.633858 |
| 8 | -2.205202 | -3.523610 | 2.359652 |

| | | | |
|---|---|---|---|
| 6 | -2.440006 | -4.851160 | 1.880161 |
| 1 | -2.679354 | -5.474374 | 2.754878 |
| 1 | -1.532338 | -5.252873 | 1.402197 |
| 6 | -3.581730 | -4.894021 | 0.884952 |
| 1 | -3.691907 | -5.935450 | 0.532621 |
| 8 | -4.796867 | -4.473003 | 1.520748 |
| 6 | -5.592943 | -3.727249 | 0.643655 |
| 1 | -6.604960 | -4.147853 | 0.549239 |
| 7 | -5.750018 | -2.370149 | 1.206160 |
| 6 | -4.802223 | -1.681955 | 1.911231 |
| 1 | -3.874776 | -2.167878 | 2.210783 |
| 7 | -5.145917 | -0.422257 | 2.129750 |
| 6 | -6.407284 | -0.295158 | 1.571934 |
| 6 | -7.275003 | 0.839483 | 1.484305 |
| 8 | -7.139272 | 1.976417 | 1.933568 |
| 7 | -8.440839 | 0.507996 | 0.758747 |
| 1 | -9.106233 | 1.271461 | 0.676231 |
| 6 | -8.718023 | -0.714684 | 0.184465 |
| 7 | -9.910354 | -0.838027 | -0.468483 |
| 1 | -10.341103 | -0.015896 | -0.873521 |
| 1 | -10.003250 | -1.692990 | -1.006144 |
| 7 | -7.915788 | -1.746259 | 0.277725 |
| 6 | -6.787552 | -1.496765 | 0.969916 |
| 6 | -3.421672 | -3.994158 | -0.352244 |
| 1 | -2.921910 | -3.063180 | -0.070934 |
| 6 | -4.892205 | -3.719642 | -0.728220 |
| 1 | -5.034046 | -2.758726 | -1.255407 |

| | | | |
|---|---|---|---|
| 8 | -5.378731 | -4.805099 | -1.477378 |
| 1 | -4.610682 | -5.102392 | -1.998599 |
| 8 | 3.229172 | -0.480172 | 0.059553 |
| 1 | 3.700191 | -0.668771 | 0.912399 |
| 1 | 2.475948 | -1.120100 | -0.158787 |
| 8 | 4.522197 | 2.659735 | -2.183970 |
| 1 | 5.509285 | 2.579875 | -2.209902 |
| 1 | 4.271078 | 3.559790 | -1.886413 |
| 8 | 5.219602 | 1.501164 | 0.425815 |
| 1 | 5.256514 | 0.711398 | 1.051407 |
| 1 | 4.885083 | 2.172432 | 1.045249 |
| 8 | -3.229548 | -0.480610 | -0.060735 |
| 1 | -3.700480 | -0.668323 | -0.913751 |
| 1 | -2.476720 | -1.120910 | 0.157599 |
| 8 | -4.522409 | 2.657754 | 2.185921 |
| 1 | -5.509516 | 2.577649 | 2.210949 |
| 1 | -4.271463 | 3.557944 | 1.888664 |
| 8 | -5.218611 | 1.502123 | -0.424886 |
| 1 | -4.883201 | 2.174076 | -1.043088 |
| 1 | -5.256044 | 0.713415 | -1.051751 |
| 15 | 4.181090 | 0.288760 | 3.380491 |
| 8 | 3.994573 | 1.892020 | 2.918275 |
| 1 | 4.018866 | 2.429617 | 3.722862 |
| 8 | 2.741083 | -4.639682 | 1.386258 |
| 1 | 1.891724 | -4.176814 | 1.586965 |
| 8 | -2.739267 | -4.641181 | -1.385915 |
| 1 | -1.891035 | -4.176743 | -1.587793 |

| | | | |
|---|---|---|---|
| 8 | -3.991979 | 1.894957 | -2.916020 |
| 1 | -4.008773 | 2.433851 | -3.719921 |
| 12 | 3.804265 | 1.099612 | -1.086318 |
| 12 | -3.803942 | 1.098233 | 1.087339 |

## 6.3 3. $Co_1G_2$

Number of imaginary frequencies : 0

Electronic energy : HF=-4802.0818241

Zero-point correction= 0.639554 (Hartree/Particle)

Thermal correction to Energy= 0.689578

Thermal correction to Enthalpy= 0.690522

Thermal correction to Gibbs Free Energy= 0.557653

Sum of electronic and zero-point Energies= -4801.442270

Sum of electronic and thermal Energies= -4801.392246

Sum of electronic and thermal Enthalpies= -4801.391302

Sum of electronic and thermal Free Energies= -4801.524171

............................................................

Cartesian Coordinates

............................................................

| | | | |
|---|---|---|---|
| 15 | -5.559987 | -1.614319 | 0.644281 |
| 8 | -6.598959 | -2.628936 | 0.354709 |
| 8 | -5.862148 | -0.574321 | 1.773822 |
| 8 | -4.129317 | -2.304856 | 1.111112 |
| 6 | -3.744508 | -3.508380 | 0.461127 |
| 1 | -4.616799 | -3.966957 | -0.031545 |
| 1 | -3.374198 | -4.212909 | 1.224038 |
| 6 | -2.634807 | -3.265968 | -0.546662 |

| | | | |
|---|---|---|---|
| 1 | -2.339367 | -4.236674 | -0.979428 |
| 8 | -3.084998 | -2.388764 | -1.595716 |
| 6 | -2.046382 | -1.413660 | -1.910037 |
| 1 | -1.328594 | -1.886793 | -2.605248 |
| 7 | -2.552462 | -0.228306 | -2.482654 |
| 6 | -1.364622 | -2.624549 | 0.046053 |
| 1 | -1.363423 | -2.617050 | 1.144205 |
| 6 | -1.412596 | -1.206663 | -0.534581 |
| 1 | -2.140658 | -0.648268 | 0.063100 |
| 8 | -0.217613 | -0.453503 | -0.524712 |
| 1 | 0.518793 | -0.927704 | -0.969772 |
| 8 | -0.256019 | -3.386562 | -0.435682 |
| 15 | 1.259713 | -3.201854 | 0.124326 |
| 8 | 2.018552 | -4.489333 | 0.026821 |
| 8 | 1.234383 | -2.381244 | 1.407617 |
| 8 | 1.853621 | -2.144498 | -1.047685 |
| 6 | 2.857472 | -2.607833 | -1.953167 |
| 1 | 2.760779 | -2.017426 | -2.875781 |
| 1 | 2.689554 | -3.670487 | -2.179678 |
| 6 | 4.261881 | -2.438927 | -1.394655 |
| 1 | 4.954016 | -3.061197 | -1.988032 |
| 8 | 4.675977 | -1.068065 | -1.493488 |
| 6 | 5.275185 | -0.637754 | -0.303760 |
| 1 | 6.196121 | -0.078161 | -0.515265 |
| 7 | 4.366106 | 0.331549 | 0.342914 |
| 6 | 3.069171 | 0.104522 | 0.758865 |
| 1 | 2.648413 | -0.884442 | 0.935562 |

| | | | |
|---|---|---|---|
| 7 | 2.383882 | 1.217191 | 0.898317 |
| 6 | 3.256878 | 2.241393 | 0.558287 |
| 6 | 3.088306 | 3.653031 | 0.470806 |
| 8 | 2.090551 | 4.352263 | 0.706252 |
| 7 | 4.265953 | 4.285354 | 0.031660 |
| 1 | 4.189269 | 5.297557 | -0.029036 |
| 6 | 5.442430 | 3.651397 | -0.305198 |
| 7 | 6.482554 | 4.427450 | -0.696556 |
| 1 | 6.354874 | 5.392225 | -0.969700 |
| 1 | 7.305438 | 3.942252 | -1.032903 |
| 7 | 5.587082 | 2.348666 | -0.231341 |
| 6 | 4.495542 | 1.694192 | 0.197790 |
| 6 | 4.436798 | -2.817072 | 0.091220 |
| 1 | 3.548516 | -2.533865 | 0.668504 |
| 6 | 5.570426 | -1.891462 | 0.529237 |
| 1 | 5.546806 | -1.676350 | 1.614299 |
| 8 | 6.812692 | -2.403882 | 0.124077 |
| 1 | 6.736282 | -3.369616 | 0.199779 |
| 7 | -4.010581 | 1.253738 | 1.725850 |
| 6 | -2.690837 | 1.011553 | 1.828425 |
| 1 | -2.260097 | 0.386508 | 2.609413 |
| 7 | -1.931648 | 1.559898 | 0.864651 |
| 6 | -2.845190 | 2.215763 | 0.052562 |
| 6 | -2.724365 | 2.926646 | -1.173710 |
| 8 | -1.724590 | 3.222688 | -1.855379 |
| 7 | -3.977998 | 3.330666 | -1.657623 |
| 1 | -3.932499 | 3.882903 | -2.508795 |

| | | | |
|---|---|---|---|
| 6 | -5.192667 | 3.096894 | -1.041997 |
| 7 | -6.302129 | 3.629623 | -1.657462 |
| 1 | -6.308598 | 3.656467 | -2.671525 |
| 1 | -7.167965 | 3.289763 | -1.250245 |
| 7 | -5.303703 | 2.452742 | 0.080032 |
| 6 | -4.133130 | 2.012696 | 0.607847 |
| 27 | 0.132608 | 1.280504 | 0.815174 |
| 8 | 0.134177 | -0.262757 | 2.257042 |
| 1 | 0.556546 | -0.149388 | 3.117842 |
| 1 | 0.501466 | -1.146577 | 1.901079 |
| 8 | 0.447247 | 2.559816 | -0.759399 |
| 1 | 0.799636 | 3.417488 | -0.469569 |
| 1 | -0.398824 | 2.773258 | -1.286156 |
| 8 | 0.456350 | 2.874484 | 2.269438 |
| 1 | 1.108905 | 2.596951 | 2.927338 |
| 1 | 0.930927 | 3.578560 | 1.765005 |
| 8 | 4.731470 | -4.161072 | 0.301809 |
| 1 | 3.852976 | -4.607602 | 0.288787 |
| 8 | -5.080197 | -0.737125 | -0.645592 |
| 1 | -4.532498 | -1.331210 | -1.207087 |
| 1 | -5.174396 | 0.181110 | 1.884819 |
| 1 | -3.217356 | 0.243337 | -1.870160 |
| 1 | -2.969099 | -0.376294 | -3.398219 |

## 6.4 4. $Mg_1G_2$

Number of imaginary frequencies : 0

Electronic energy : HF=-3619.5748274

```
Zero-point correction=                           0.640804 (Hartree/Particle)
Thermal correction to Energy=                    0.690390
Thermal correction to Enthalpy=                  0.691335
Thermal correction to Gibbs Free Energy=         0.561162
Sum of electronic and zero-point Energies=          -3618.934023
Sum of electronic and thermal Energies=             -3618.884437
Sum of electronic and thermal Enthalpies=           -3618.883493
Sum of electronic and thermal Free Energies=        -3619.013665
```

............................................................

Cartesian Coordinates

............................................................

| | | | |
|---|---|---|---|
| 15 | -5.562920 | -1.599717 | 0.699896 |
| 8 | -6.583169 | -2.637955 | 0.429274 |
| 8 | -5.879109 | -0.552540 | 1.819412 |
| 8 | -4.114766 | -2.254734 | 1.163815 |
| 6 | -3.715044 | -3.461876 | 0.529139 |
| 1 | -4.583610 | -3.941442 | 0.050202 |
| 1 | -3.327937 | -4.149252 | 1.299235 |
| 6 | -2.617279 | -3.216862 | -0.491089 |
| 1 | -2.320737 | -4.187362 | -0.923141 |
| 8 | -3.084600 | -2.346658 | -1.539216 |
| 6 | -2.062667 | -1.359910 | -1.864217 |
| 1 | -1.349055 | -1.819950 | -2.572181 |
| 7 | -2.589497 | -0.176274 | -2.423014 |
| 6 | -1.342647 | -2.566888 | 0.083480 |
| 1 | -1.327005 | -2.555643 | 1.181446 |
| 6 | -1.408286 | -1.150764 | -0.499510 |

| | | | |
|---|---|---|---|
| 1 | -2.122727 | -0.588257 | 0.111810 |
| 8 | -0.208553 | -0.397155 | -0.520412 |
| 1 | 0.522788 | -0.878669 | -0.969988 |
| 8 | -0.237269 | -3.319883 | -0.412672 |
| 15 | 1.282948 | -3.138399 | 0.145757 |
| 8 | 2.037982 | -4.427029 | 0.037971 |
| 8 | 1.259797 | -2.318424 | 1.427526 |
| 8 | 1.867539 | -2.080683 | -1.032975 |
| 6 | 2.867555 | -2.541102 | -1.945780 |
| 1 | 2.765481 | -1.946073 | -2.864780 |
| 1 | 2.695378 | -3.602289 | -2.175982 |
| 6 | 4.278252 | -2.379934 | -1.399329 |
| 1 | 4.958752 | -3.004488 | -2.003842 |
| 8 | 4.701801 | -1.012246 | -1.496981 |
| 6 | 5.319745 | -0.591433 | -0.311376 |
| 1 | 6.241355 | -0.035509 | -0.530445 |
| 7 | 4.420725 | 0.373697 | 0.351997 |
| 6 | 3.110224 | 0.139856 | 0.720901 |
| 1 | 2.688203 | -0.851326 | 0.866592 |
| 7 | 2.418295 | 1.246391 | 0.861932 |
| 6 | 3.306653 | 2.274034 | 0.587960 |
| 6 | 3.134141 | 3.684259 | 0.558693 |
| 8 | 2.117128 | 4.357528 | 0.787466 |
| 7 | 4.323549 | 4.339706 | 0.189338 |
| 1 | 4.237983 | 5.351385 | 0.134726 |
| 6 | 5.513247 | 3.719671 | -0.132506 |
| 7 | 6.548534 | 4.509587 | -0.505471 |

| | | | |
|---|---|---|---|
| 1 | 6.532034 | 5.513461 | -0.395625 |
| 1 | 7.435583 | 4.053123 | -0.676432 |
| 7 | 5.662070 | 2.415071 | -0.105253 |
| 6 | 4.556370 | 1.740129 | 0.251827 |
| 6 | 4.471261 | -2.769373 | 0.081643 |
| 1 | 3.593778 | -2.487645 | 0.676132 |
| 6 | 5.618308 | -1.853789 | 0.508200 |
| 1 | 5.612972 | -1.645549 | 1.594645 |
| 8 | 6.850102 | -2.373251 | 0.079822 |
| 1 | 6.768747 | -3.338636 | 0.153867 |
| 7 | -4.053066 | 1.303529 | 1.747203 |
| 6 | -2.729159 | 1.106612 | 1.875460 |
| 1 | -2.295149 | 0.495775 | 2.666845 |
| 7 | -1.964720 | 1.673383 | 0.920945 |
| 6 | -2.890360 | 2.287289 | 0.087919 |
| 6 | -2.761469 | 3.010835 | -1.130957 |
| 8 | -1.752833 | 3.346646 | -1.776950 |
| 7 | -4.016913 | 3.375994 | -1.642789 |
| 1 | -3.970843 | 3.932521 | -2.491279 |
| 6 | -5.236643 | 3.105708 | -1.050953 |
| 7 | -6.348298 | 3.605335 | -1.687272 |
| 1 | -6.333783 | 3.649362 | -2.700391 |
| 1 | -7.213326 | 3.240749 | -1.300542 |
| 7 | -5.352178 | 2.457816 | 0.069345 |
| 6 | -4.180102 | 2.049998 | 0.619171 |
| 8 | 0.148691 | -0.176758 | 2.264993 |
| 1 | 0.567830 | 0.057013 | 3.102354 |

| | | | |
|---|---|---|---|
| 1 | 0.545675 | -1.067484 | 1.979138 |
| 8 | 0.462795 | 2.577390 | -0.783063 |
| 1 | 0.883417 | 3.404632 | -0.496411 |
| 1 | -0.390353 | 2.858336 | -1.253550 |
| 8 | 0.342484 | 2.846796 | 2.212243 |
| 1 | 0.947879 | 3.532134 | 1.842399 |
| 1 | -0.538825 | 3.248013 | 2.245177 |
| 8 | 4.761283 | -4.117065 | 0.276950 |
| 1 | 3.880765 | -4.558763 | 0.274602 |
| 8 | -5.110194 | -0.726596 | -0.602444 |
| 1 | -4.556572 | -1.316329 | -1.162181 |
| 12 | 0.146293 | 1.267591 | 0.773746 |
| 1 | -5.203095 | 0.212831 | 1.919539 |
| 1 | -3.008398 | -0.321091 | -3.338070 |
| 1 | -3.258134 | 0.279594 | -1.802487 |

(19) Frisch, M. J.; Trucks, G. W.; Schlegel, H. B.; Scuseria, G. E.; Robb, M. A.; Cheeseman, J. R.; Scalmani, G.; Barone, V.; Petersson, G. A.; Nakatsuji, H.; Li, X.; Caricato, M.; Marenich, A. V.; Bloino, J.; Janesko, B. G.; Gomperts, R.; Mennucci, B.; Hratchian, H. P.; Ortiz, J. V.; Izmaylov, A. F.; Sonnenberg, J. L.; Williams-Young, D.; Ding, F.; Lipparini, F.; Egidi, F.; Goings, J.; Peng, B.; Petrone, A.; Henderson, T.; Ranasinghe, D.; Zakrzewski, V. G.; Gao, J.; Rega, N.; Zheng, G.; Liang, W.; Hada, M.; Ehara, M.; Toyota, K.; Fukuda, R.; Hasegawa, J.; Ishida, M.; Nakajima, T.; Honda, Y.; Kitao, O.; Nakai, H.; Vreven, T.; Throssell, K.; Montgomery, J. A., Jr.; Peralta, J. E.; Ogliaro, F.; Bearpark, M. J.; Heyd, J. J.; Brothers, E. N.; Kudin, K. N.; Staroverov, V. N.; Keith, T. A.; Kobayashi, R.; Normand, J.; Raghavachari, K.; Rendell, A. P.; Burant, J. C.; Iyengar, S. S.; Tomasi, J.; Cossi, M.; Millam, J. M.; Klene, M.;

Adamo, C.; Cammi, R.; Ochterski, J. W.; Martin, R. L.; Morokuma, K.; Farkas, O.; Foresman, J. B.; Fox, D. J. Gaussian 16 Revision C.01. 2016; Gaussian Inc. Wallingford CT.

(20) Grimme, S.; Antony, J.; Ehrlich, S.; Krieg, H. A consistent and accurate ab initio parametrization of density functional dispersion correction (DFT-D) for the 94 elements H-Pu. *J. Chem. Phys* **2010**, *132*.

(21) Weigend, F.; Ahlrichs, R. Balanced basis sets of split valence, triple zeta valence and quadruple zeta valence quality for H to Rn: Design and assessment of accuracy. *Phys. Chem. Chem. Phys.* **2005**, *7*, 3297–3305.

(22) Marenich, A. V.; Cramer, C. J.; Truhlar, D. G. Universal solvation model based on solute electron density and on a continuum model of the solvent defined by the bulk dielectric constant and atomic surface tensions. *J. Phys. Chem. B* **2009**, *113*, 6378–6396.

(23) Bader, R. F. W. *Atoms in Molecules: A Quantum Theory*; Oxford University Press: Oxford, New York, 1994.

(24) Lu, T.; Chen, F. Multiwfn: A multifunctional wavefunction analyzer. *J. Comput. Chem.* **2012**, *33*, 580–592.

(25) Guo, Z.; Thirumalai, D. Kinetics and thermodynamics of folding of a de Novo designed four-helix bundle protein. *J. Mol. Biol.* **1996**, *263*, 323–343.

(26) Wang, J.; Williams, B., II; Chirasani, V. R.; Krokhotin, A.; Das, R.; Dokholyan, N. V. Limits in accuracy and a strategy of RNA structure prediction using experimental information. *Nucleic Acids Res.* **2019**, *47*, 5563–5572.

(27) Hori, N.; Denesyuk, N. A.; Thirumalai, D. Shape changes and cooperativity in the

folding of the central domain of the 16S ribosomal RNA. *Proc. Nat. Acad. Sci.* **2021**, *118*.

(28) Kumar, S.; Reddy, G. TPP riboswitch populates holo-form-like structure even in the absence of cognate ligand at high $Mg^{2+}$ concentration. *J. Phys. Chem. B* **2022**, *126*, 2369–2381.

(29) Humphrey, W.; Dalke, A.; Schulten, K. VMD – Visual Molecular Dynamics. *J. Mol. Graph.* **1996**, *14*, 33–38.

(30) Kabsch, W. A discussion of the solution for the best rotation to relate two sets of vectors. *Acta Cryst. A* **1978**, *34*, 827–828.

(31) Eargle, J.; Wright, D.; Luthey-Schulten, Z. Multiple alignment of protein structures and sequences for VMD. *Bioinformatics* **2006**, *22*, 504–506.

(32) Stone, J. An efficient library for parallel ray tracing and animation. M.Sc. thesis, Computer Science Department, University of Missouri-Rolla, 1998.
